# Large-scale discovery, analysis, and design of protein energy landscapes

**DOI:** 10.1101/2025.03.20.644235

**Authors:** Állan J. R. Ferrari, Sugyan M. Dixit, Jane Thibeault, Mario Garcia, Scott Houliston, Robert W. Ludwig, Pascal Notin, Claire M. Phoumyvong, Cydney M. Martell, Michelle D. Jung, Kotaro Tsuboyama, Lauren Carter, Cheryl H. Arrowsmith, Miklos Guttman, Gabriel J. Rocklin

## Abstract

All folded proteins continuously fluctuate between their low-energy native structures and higher energy conformations that can be partially or fully unfolded. These rare states influence protein function, interactions, aggregation, and immunogenicity, yet they remain far less understood than protein native states. Although native protein structures are now often predictable with impressive accuracy, conformational fluctuations and their energies remain largely invisible and unpredictable, and experimental challenges have prevented large-scale measurements that could improve machine learning and physics-based modeling. Here, we introduce a multiplexed experimental approach to analyze the energies of conformational fluctuations for hundreds of protein domains in parallel using intact protein hydrogen-deuterium exchange mass spectrometry. We analyzed 5,778 domains 28-64 amino acids in length, revealing hidden variation in conformational fluctuations even between sequences sharing the same fold and global folding stability. Site-resolved hydrogen exchange NMR analysis of 13 domains showed that these fluctuations often involve entire secondary structural elements with lower stability than the overall fold. Computational modeling of our domains identified structural features that correlated with the experimentally observed fluctuations, enabling us to design mutations that stabilized low-stability structural segments. Our dataset enables new machine learning-based analysis of protein energy landscapes, and our experimental approach promises to reveal these landscapes at unprecedented scale.

## Main

All proteins continuously move between different conformational states, including (typically) a low-energy native folded state, a higher-energy unfolded state, and diverse excited states with different levels of native-like structure. Although each protein molecule occupies its high-energy states only a small fraction of the time, these states have large impacts across biology and protein engineering, influencing protein function (Rossi et al. 2022; Glasgow et al. 2023), interactions (Hodge et al. 2022), aggregation (Dumoulin et al. 2005; Codina et al. 2019; Volz et al. 2025; Pal and Udgaonkar 2024a, 2024b), and immunogenicity (de Taeye et al. 2015; Calvaresi et al. 2024). The rare and often transient nature of high-energy states makes them challenging to study experimentally, and these states are often described as “invisible” to traditional structural biology (Alderson and Kay 2020). As a result, we know much less about protein excited states compared to protein native states, with no comparable resource to the Protein Data Bank to guide the development of artificial intelligence (AI) and machine learning (ML) and methods (but see Pancsa et al. 2016). AI methods trained to predict native (lowest energy) protein structures have shown little ability to predict protein folding stabilities or the energies of different conformational states without additional data (Kim et al. 2022; Buel and Walters 2022; Pak et al. 2023; Chakravarty, Lee, and Porter 2025).

Along with the experimental difficulties, high-energy states are also challenging to understand because they are highly sensitive to a protein’s specific amino acid sequence. Every protein sequence has its own *conformational energy landscape* describing the energies (and hence populations) of its different conformational states. Energy landscapes can vary significantly between structurally similar proteins, and single mutations can strongly perturb energy landscapes without altering the native protein structure (Spudich et al. 2004; Dumoulin et al. 2005; Shaw et al. 2006; Volz et al. 2025). AI methods for predicting native structures rely on the conservation of protein structures across highly diverged sequences, but this conservation does not hold for energy landscapes (Cota et al. 2000; Oyeyemi et al. 2011; Lim and Marqusee 2018; Chisholm et al. 2024). To develop next-generation AI models that can predict and engineer conformational energy landscapes, we need new experimental methods that can characterize energy landscapes across sequence space and reveal the rules for how protein sequences determine their energy landscapes in a particular environment. Recent advances in measuring global folding stability at scale (Rocklin et al. 2017; Tsuboyama et al. 2023) have accelerated the use of AI methods in biophysics (Dieckhaus et al. 2024; Diaz et al. 2024; Lewis et al. 2024). However, these high-throughput methods do not yet have the ability to resolve the details of conformational fluctuations or identify the range of excited states populated by each protein sequence.

Here, we introduce a multiplexed hydrogen-deuterium exchange mass spectrometry (mHDX-MS) strategy to investigate protein energy landscapes for hundreds of protein domains simultaneously. Unlike experiments that only probe global stability (Araya et al. 2012; Rocklin et al. 2017; Walker et al. 2019; Weng et al. 2024; Faure et al. 2022; Beltran et al. 2025; Tsuboyama et al. 2023), hydrogen-deuterium exchange measures the energies of individual residues transitioning from “closed” conformations (typically in secondary structure) to higher energy “open” conformations. Within the same protein, different residues open by accessing different states on the energy landscape, including nearly native states that expose only a few additional residues, alternative folded conformations, partially unfolded states, and the globally unfolded state. This enables HDX experiments to measure energies for states that are invisible to other approaches. Traditionally, HDX experiments have been limited to studying one or a few purified proteins at a time (Rossi et al. 2022; Glasgow et al. 2023; Hodge et al. 2022; Dumoulin et al. 2005; Bryan Francis Shaw et al. 2006; Codina et al. 2019; Volz et al. 2025; Pal and Udgaonkar 2024a, 2024b; de Taeye et al. 2015; Calvaresi et al. 2024; Wales et al. 2024), although recent studies have employed simplified cell lysates (Fang et al. 2023; Moroco et al. 2025). To enable large-scale HDX analysis of both natural and designed protein domains, we leveraged DNA oligo pool library synthesis to produce customized synthetic proteomes comprising up to 1,300 small protein domains in a single mixture (28-64 amino acids in length). Analyzing these mixtures by mHDX-MS revealed the exchange rate distributions and approximate opening energy (ΔG_open_) distributions for each protein domain.

Overall, we measured the opening energy distributions of 5,778 protein domains from ten families under identical experimental conditions (3,590 domains after removing low stability domains). Our dataset revealed diverse energy landscapes across sequences with the same overall fold, differences in landscapes between domains sharing the same global folding stability, and systematic differences between domain families. The unique scale of our data (to our knowledge >500-fold larger than previous comparative studies of energy landscapes (Orban et al. 1995; Hollien and Marqusee 1999; Goedken and Marqusee 2001; Dumoulin et al. 2005; Wales and Engen 2006) enabled us to use machine learning to discover common determinants of energy landscapes across a broad range of sequences. Our analysis also enabled us to design mutations that enhanced local stability by dampening conformational fluctuations, demonstrating the potential of data-driven approaches to modulate protein energy landscapes.

### The multiplex hydrogen deuterium exchange mass spectrometry method

Protein domains have individual energy landscapes that influence how backbone amide hydrogens exchange for deuterium. In an idealized two-state protein with no conformational fluctuations, all protected amides (typically those donating hydrogen bonds in secondary structure, but not always (McAllister and Konermann 2015; Skinner et al. 2012) can only exchange from the globally unfolded state (**Fig. 1A left**). This leads to a uniform distribution of opening free energies (Δ*G*_open_) across all protected residues because all protected amides exchange from the same state (**Fig. 1B, blue**). If the energy landscape includes additional states at intermediate energies that open a subset of amides (**Fig. 1A right**), the distribution of Δ*G*_open_ will be less uniform (**Fig. 1B, red**). We developed multiplexed hydrogen-deuterium exchange mass spectrometry (mHDX-MS) to measure these opening energy distributions for hundreds of protein domains in parallel (**Fig. 1C**).

**Figure 1:**
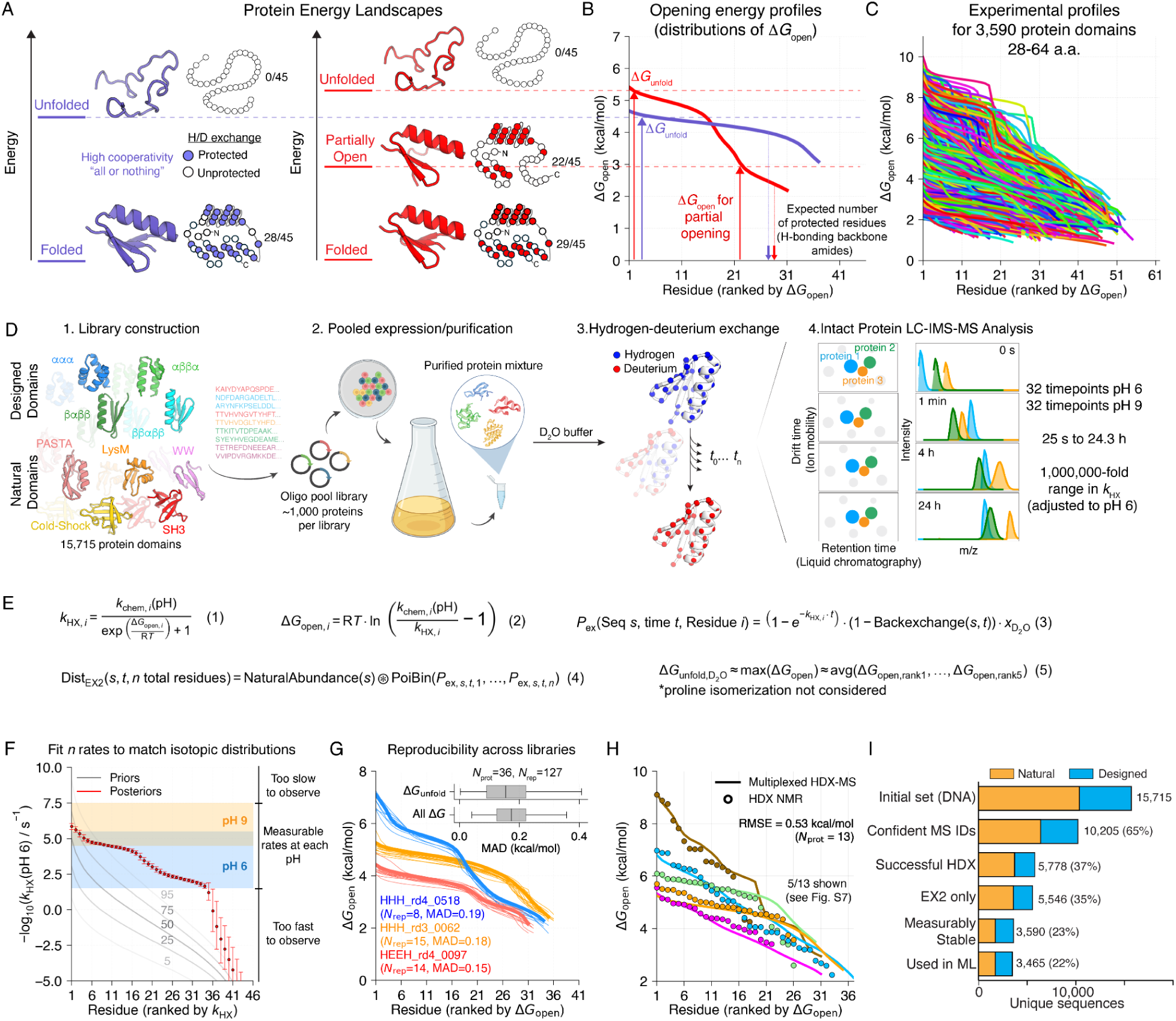
Multiplex hydrogen-deuterium exchange mass spectrometry (mHDX-MS) to measure energy landscapes. **A)** Idealized, hypothetical energy landscapes for a perfectly two-state protein (left) and a protein with a partially open state (right). Many residues are protected from exchange (filled circles) in the folded state and exchange (open) through conformational fluctuations to higher energy states. **B)** mHDX-MS opening energy profiles for two proteins resembling the idealized energy landscapes in A. The two-step ΔG_open_ distribution (red) reveals many residues opening through low energy fluctuations (not global unfolding). **C)** Diverse opening energy profiles discovered by mHDX-MS. **D)** The mHDX-MS method. Protein domains are encoded in a DNA oligo pool, expressed together in a single *E. coli* culture, purified as a mixture, and analyzed together at 64 HDX timepoints by liquid chromatography-ion mobility-mass spectrometry (LC-IMS-MS). LC retention time and IMS drift time remain constant at each timepoint and uniquely identify each protein signal. **E)** Kinetic model for HDX. Under our experimental conditions (EX2), first-order residue exchange rates *k*_HX_ are determined by equilibrium opening free energies Δ*G*_open_and pH-dependent open-state exchange rates *k*_chem_ (Eq. 1-2, Konermann et al. 2008). Residue exchange probabilities *P_ex_* at time *t* depend on *k*_HX_, the back exchange probability, and the mole fraction *x*_D2O_ (Eq. 3). In this EX2 regime (independent exchange), protein mass distributions are the convolution of natural isotopic abundance and a Poisson Binomial (PoiBin) distribution determined by all residues’ *P*_ex_ (Eq. 4). We measure mass distributions over time, then infer all *k*_HX_ and all Δ*G*_open_ based on approximate *k*_chem_ (Methods). Δ*G*_unfold_ is computed from the most stable residues’ Δ*G*_open_ (Eq. 5, Huyghues-Despointes et al. 1999). **F**) Bayesian approach to infer *n* exchange rates (*k*_HX_) for *n* exchangeable residues, see also **Fig. S2**. Rates are pH-dependent due to *k*_chem_ (Nguyen et al. 2018); slower rates are resolved at pH 9 and converted to a pH 6 scale (Methods) **G)** Reproducibility of opening energy profiles (measurable Δ*G*_open_ values) for the same domains analyzed in *N*_rep_ different libraries, along with the Mean Absolute Deviation (MAD) between all replicates and the highest-quality replicate (thick line). Inset shows MAD distributions from 36 domains with > 1 replicate, see also **Fig. S5** and Methods **H)** Opening energy profiles from mHDX-MS are consistent with HDX NMR; all comparisons in **Fig. S7. I)** Experimental outcomes of all sequences.

Our approach begins by constructing customized mixtures of protein domains. Each sample of 108-1,334 domains is encoded as a synthetic DNA oligo pool, cloned into a vector, and expressed and purified as a mixture from a single *E. coli* culture (**Fig. 1D**). We then incubate the mixture in deuterium oxide (D_2_O) for timepoints ranging from 25 seconds to 24 hours, quench the exchange reaction, and analyze each timepoint (32 at pH 6 and 32 at pH 9) by liquid chromatography ion mobility mass spectrometry (LC-IMS-MS, **Fig. 1D**). Using a customized computational pipeline, we extract the full isotopic distributions for each domain at each timepoint. Our pipeline incorporates a novel tensor factorization approach to deconvolute overlapping isotopic distributions and an optimization algorithm that identifies the most likely signal for each domain at each timepoint (**Fig. S1**). All analysis is fully automated, with no manual intervention beyond establishing global data quality thresholds.

We use Bayesian inference to infer each domain’s set of exchange rates (*k*_HX_, *n* rates for *n* exchangeable residues in a domain) based on that domain’s isotopic distributions at each timepoint (**Fig. 1E**: Eq. 1-4; **Fig. 1F**, **Fig. S2**). Although our analysis infers *n* rates for each domain, our measurements on intact domains do not resolve which rates stem from which residues in the sequence. The Bayesian procedure naturally infers the experimental uncertainty for each rate, and these uncertainties are typically low due to the large number of timepoints measured. Rates that cannot be measured on our timescales show high uncertainty, as expected (**Fig. 1F**). We then compute an approximate opening energy (Δ*G*_open_) distribution for each domain based on its exchange rate (*k*_HX_) distribution and its expected exchange rates in the unfolded state (*k*_chem_, **Fig. 1E**: Eq. 2, **Fig. S3**, see also Methods). These Δ*G*_open_ distributions also reveal the global stability (Δ*G*_unfold_) of each domain based on the opening energy of the most stable residue (Huyghues-Despointes, Scholtz, and Pace 1999; Bai et al. 1994); to reduce noise we average the five most stable residues (**Fig. 1E**: Eq. 5 and **Fig. S4**). Our experiments and analysis protocol produce reproducible *k*_HX,_ Δ*G*_open_, and Δ*G*_unfold_ measurements when the same domains are analyzed in different libraries (**Fig. 1G, Fig. S5**).

Our inferred *k*_HX_ and Δ*G*_open_ distributions depend on several assumptions. First, we assume all exchanges are independent and follow first-order kinetics (EX2 assumption; **Fig. 1E**: Eq. 3-4, see Methods and **Fig. S6** for details on removing EX1 data). Second, for many domains we combine data collected at pH 6 and pH 9 into a single overall model; we assume this pH shift does not alter the energy landscape (measuring exchange at each pH expands the measurable range of Δ*G*_open_ by modulating *k_c_*_hem_, **Eq. 1**). Third, to model the loss of deuterium during LC-IMS-MS analysis (back exchange), we assume all exchanged residues in a domain have an equal probability of back exchange (see Methods). Finally, our Δ*G*_open_ distributions are only approximate because they depend on approximate estimates of *k*_chem_ (see **Fig. S3** and Methods). Although these assumptions will not hold for every domain, their overall validity is supported by (a) the consistency between our observed and modeled data (see **Table S1**: **Dataset_8** for all raw data and fits), and (b) the consistency between mHDX-MS, HDX NMR, and cDNA display proteolysis data (see below), including for domains with unusual energy landscapes and unique dynamics that were initially discovered by mHDX-MS.

### Multiplexed HDX-MS measurements are accurate

We analyzed the accuracy of our approach using comparisons to HDX NMR and cDNA display proteolysis measurements. HDX NMR measures “gold standard” values for *k*_HX_ and Δ*G*_open_ one protein at a time (requiring several days), whereas cDNA display proteolysis measures Δ*G*_unfold_ for up to 900,000 domains in parallel (Tsuboyama et al. 2023). Across 13 different domains, mHDX-MS measurements were highly consistent with HDX NMR measurements, with RMSE of 1.9-fold for *k*_HX_ distributions and 0.53 kcal/mol for Δ*G*_open_ distributions (**Fig. 1H**; all 13 domains shown in **Fig. S7**). Discrepancies between mHDX-MS measurements and HDX NMR measurements stem from the assumptions above as well as small experimental differences in temperature, pH, the fraction of D_2_O during exchange, and the exact protein constructs (see Methods). Global stabilities (Δ*G*_unfold_) measured for 4,464 domains by mHDX-MS (in D_2_O) and cDNA display proteolysis (in H_2_O) were also strongly correlated (R=0.78, **Fig. S8**). Stabilities were typically 1.6 kcal/mol higher in mHDX-MS experiments, likely due to the stabilizing effect of D_2_O (Efimova et al. 2007; Pica and Graziano 2018; Stadmiller and Pielak 2018; Haidar and Konermann 2023). mHDX-MS measurements also resolved a wider range of folding stability (approximately 2-9 kcal/mol in mHDX-MS compared to 0-5 kcal/mol in cDNA display proteolysis, **Fig. S8**). Comparing mHDX-MS and cDNA display proteolysis revealed that nucleic acid binding domains have inflated folding stability in cDNA display proteolysis, likely due to stabilization conferred by binding DNA (**Fig. S8**).

### Measuring energy landscapes across domain families

From an initial pool of 15,715 synthetic DNA sequences, we successfully analyzed 5,778 domains by mHDX-MS. These domains came from four families of *de novo* designed sequences (ααα, βαββ, αββα, and ββαββ, Rocklin et al. 2017; Kim et al. 2022), six families of naturally occurring domains from the Pfam database (LysM, PASTA, WW, SH3, Pyrin, and Cold-Shock domains, (Mistry et al. 2021), and an assortment of other small domains taken from the Protein Data Bank (PDB). Within each family, pairwise sequence identities averaged 10% to 41% (**Fig. S9**). Domains were assayed in 18 separate libraries ranging in size from 108 to 1,334 sequences. Different sequences failed at different stages owing to differences in protein expression, signal intensity, or HDX data quality (**Fig. 1I**). Success also varied by protein family (**Fig. S10**). 54% of successfully analyzed natural domains had fully completed exchange before our first time point; this indicates a low folding stability (Δ*G*_unfold_ < 2 kcal/mol). In contrast, only 10% of designed domains were this unstable, likely because we selected our designed sequences based on previous experimental stability data (Rocklin et al. 2017; Kim et al. 2022). The largest number of domains successfully analyzed in one library after applying HDX data quality filters was 519, from a library with 1,311 initial sequences (**Fig. S10C**). All domains became fully deuterated after 24 hours at pH 9 according to our criteria (Methods) except for 42 extremely stable domains (mainly PASTA domains, **Fig. S11**).

### Quantifying opening cooperativity using opening free energy distributions

Our mHDX-MS experiments revealed diverse Δ*G*_open_ distributions across our set of 3,590 stable domains. Unlike large-scale measurements of fitness (Jacquier et al. 2013; Fowler and Fields 2014; Notin et al. 2023) or global stability (Araya et al. 2012; Rocklin et al. 2017; Walker et al. 2019; Faure et al. 2022; Tsuboyama et al. 2023; Weng et al. 2024; Beltran et al. 2025), Δ*G*_open_ distributions are inherently multidimensional and can differ between domains in complex ways. One axis of variation is global stability (Δ*G*_unfold_ measured based on the most stable residues, **Fig. 1E**: Eq. 5; (Bai et al. 1994; Huyghues-Despointes et al. 1999), which varied from below 2 to ∼9 kcal/mol (in D_2_O). However, most residues across our domains exchange through conformational fluctuations that are lower in energy than Δ*G*_unfold_ (**Fig. 1C**, **Fig. 2A**). To quantify this and reduce the dimensionality of our data, we computed the average Δ*G*_open_ over all exchangeable residues for each domain (Δ*G*_avg_, unmeasurably fast residues are set to a lower bound of 0 kcal/mol, **Fig. 2A**). Domains with the same Δ*G*_unfold_ and similar native structures often differed substantially in Δ*G*_avg_ (**Fig. 2A**). In some cases, this reflects differences in hydrogen bonding in different domains, because residues lacking amide H-bonds typically have low ΔG_open_ regardless of conformational stability (these residues are “open” in the native state, McAllister and Konermann 2015; Skinner et al. 2012). The remaining variation in Δ*G*_avg_ (after hydrogen bonding has been accounted for) reveals hidden differences in conformational fluctuations between different domains—even between domains with similar native structures and similar global stability.

**Figure 2:**
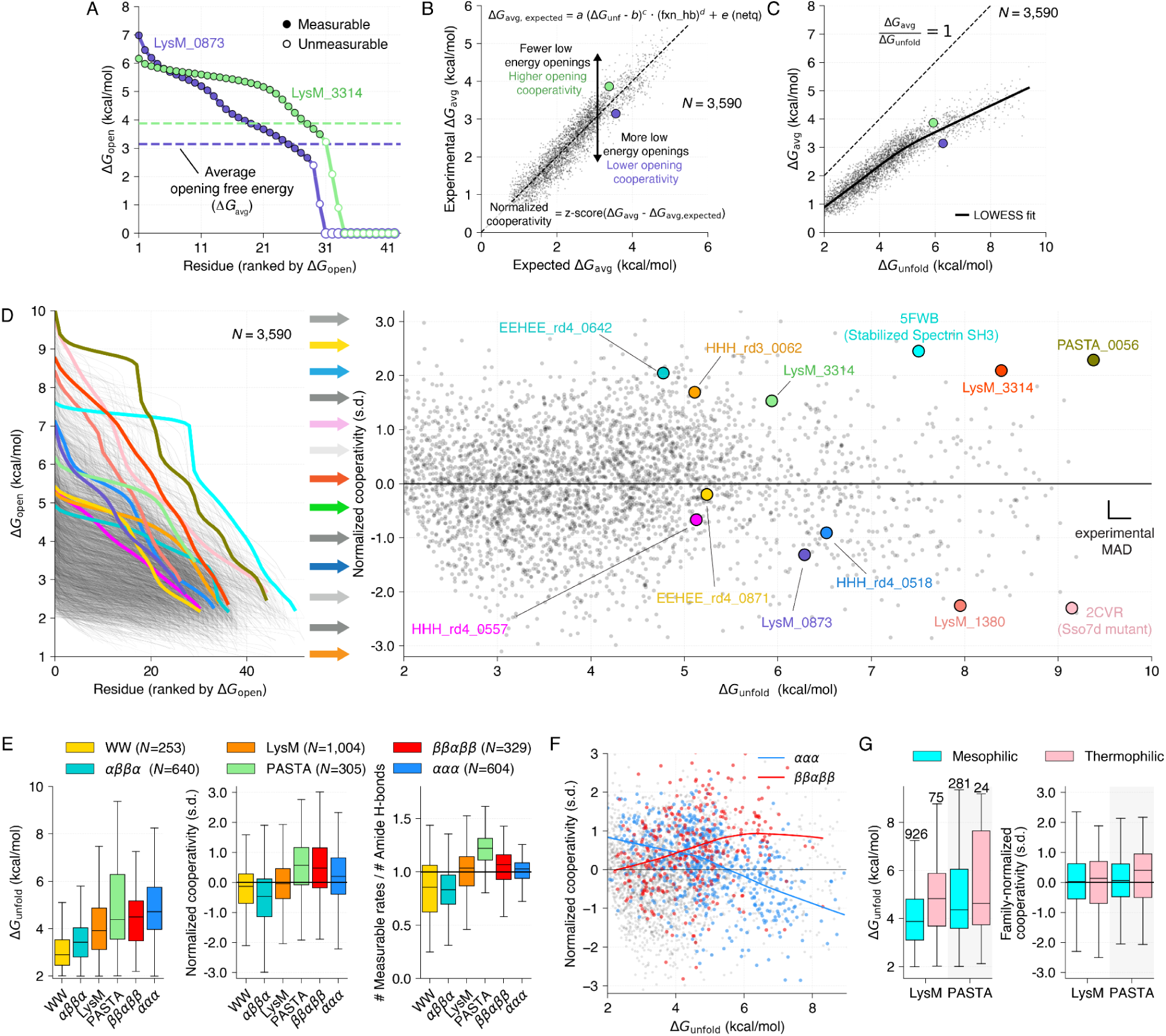
Small domains vary in stability and opening cooperativity. **A)** We calculate the average opening energy (Δ*G*_avg_) over all exchangeable residues as a proxy measure for the energies of partially open states. Low stability residues with unmeasurably fast exchange rates are included in the average and set at 0 kcal/mol. **B)** Five-parameter empirical model to predict Δ*G*_avg_from (1) Δ*G*_unfold_, (2) the fraction of backbone amide residues donating hydrogen bonds (fxn_hb), and (3) the protein net charge (netq). We define *normalized cooperativity* as the standardized difference (z-score) between each protein’s observed Δ*G*_avg_ and the model’s expected Δ*G*_avg_. Green and blue circles show the protein domains from (A). **C)** Experimental Δ*G*_avg_ increases sublinearly with Δ*G*_unfold_, highlighting the proteins in (A). **D)** Opening energy profiles for all domains shown in full (left) or simplified to two dimensions: Δ*G*_unfold_ (x-axis) and normalized cooperativity (y-axis). Twelve proteins are highlighted and shown in the same color in each plot. Scale bars show experimental mean absolute deviations (MAD) between replicates measured in different libraries. **E)** Distributions of Δ*G*_unfold_ (left), normalized cooperativity (middle), and the ratio of measurable rates to backbone amide H-bonds (right) for stable domains in the six families with the most experimental data. **F)** Normalized cooperativity trends with Δ*G*_unfold_ for two protein families; colored lines show LOWESS fits. **G)** Distributions of Δ*G*_unfold_ (left) and family-normalized cooperativity (right) for domains from mesophilic and thermophilic organisms.

To isolate these differences in conformational fluctuations, we built a simple five-parameter empirical model of Δ*G*_avg_ based on Δ*G*_unfold_, hydrogen bonding, and protein net charge (included for technical reasons, see Methods). This model explains 89% of the variance in Δ*G*_avg_ (**Fig. 2B**). The remaining variance in Δ*G*_avg_ (the model residuals) results from differences in conformational fluctuations. Domains with a positive residual have high conformational stability (high Δ*G*_open_) throughout the full domain (**Fig. 2A** and **2B**, green), compared to the typical domain in our dataset with similar global stability. Domains with a negative residual (**Fig. 2A** and **2B**, blue) have low energy fluctuations (low Δ*G*_open_) across many residues that lead to a low Δ*G*_avg_, compared to what our model expects given Δ*G*_unfold_ and hydrogen bonding. We define each domain’s normalized residual (a z-score) as its *normalized cooperativity*; a metric reflecting the level of conformational fluctuations in the domain compared to the typical domain with similar stability in our dataset. In protein folding, proteins are considered cooperative when they fold in a two-state, “all-or-nothing” manner. Hydrogen exchange studies have complicated our understanding of cooperativity by revealing numerous partially folded states across many proteins that previously appeared “two-state” (Bai et al. 1995; Chamberlain et al. 1996; Llinás et al. 1999; Englander et al. 2002; Maity et al. 2003). Our normalized cooperativity metric describes a continuum in cooperativity between domains where partially open states are relatively rare and high in energy, approximating ideal “two-state” behavior (**Fig. 2A**, green) and domains with significant low energy fluctuations to partially open states affecting many residues (**Fig. 2A**, blue). We use the term *opening cooperativity* to refer to the degree of all-or-nothing character in each domain’s opening energy landscape (conformational “openings” are a broad category that includes global and local “unfolding” as well as smaller local fluctuations, see Bai et al. 1995; Chamberlain et al. 1996; Llinás et al. 1999; Maity et al. 2003).

How does opening cooperativity relate to folding stability? We compared Δ*G*_avg_ to Δ*G*_unfold_ across all 3,590 stable domains and found Δ*G*_avg_ increases sub-linearly with Δ*G*_unfold_ (**Fig. 2C**, **Fig. S12**), indicating decreased opening cooperativity in higher stability proteins (the highest stability proteins have significant openings occuring at much lower energies). This has been described before: high stability proteins often appear less cooperative when examined by hydrogen exchange (Llinás et al. 1999) because they have a wider range of energies below Δ*G*_unfold_ where partially open states can be found (Englander, Mayne, and Rumbley 2002). Our empirical model (**Fig. 2B**) captures this sub-linear dependence so that normalized cooperativity (the normalized model residuals) is nearly orthogonal to Δ*G*_unfold_. By defining normalized cooperativity as relative to a typical domain with the same stability using our empirical model, we decouple normalized cooperativity from stability (**Fig. 2D**), enabling us to analyze how protein features directly influence cooperativity independently from stability.

We observed systematic differences in stability and cooperativity between different domain families (**Fig. 2E-F**). However, the variation in cooperativity within each family was larger than the average differences between families. This indicates that the individual sequences of the domains strongly influence cooperativity beyond the average tendency of the overall fold. Of the six families with the most data, PASTA domains (an α/β fold) and *de novo* designed ββαββ domains showed the highest average cooperativity (**Fig. 2E**), possibly owing to their β-sheet content and the placement of one (PASTA) or both (ββαββ) terminal strand(s) fully inside the sheet. The highly cooperative PASTA domain family also showed a relatively high number of protected residues compared to the number of H-bonding backbone amides in the predicted reference structures (**Fig. 2E**; cases of unexpected protection have been found previously; McAllister and Konermann 2015; Skinner et al. 2012). Although normalized cooperativity is (intentionally) nearly orthogonal to stability across all 3,590 stable domains (**Fig. 2D**), certain domain families showed clear relationships between stability and cooperativity within the family (**Fig. 2F**). To better compare normalized cooperativity within families, we defined an additional *family-normalized cooperativity* metric by fitting the empirical model from **Fig. 2B** separately for each different family (**Fig. S12, Table S2**). Finally, we compared stability and cooperativity between mesophiles and thermophiles within the LysM and PASTA families (the only families with enough data from thermophiles, (Engqvist 2018), **Fig. 2G**). LysM domains from thermophiles showed significantly higher global stability on average (mean difference of 0.8 ± 0.4 kcal/mol, mean ± 95 C.I. from bootstrapping); PASTA domains showed a similar but statistically insignificant difference (0.4 ± 0.9 kcal/mol). We did not observe any significant differences in normalized cooperativity (mean differences of 0.0 ± 0.2 and 0.2 ± 0.5 for LysM and PASTA domains respectively).

### The spatial distribution of stability

Highly cooperative domains are all alike; every less cooperative domain fluctuates in its own way. Low stability residues might be dispersed throughout the structure (perhaps due to secondary structure elements fraying at the ends), or they could be clustered in specific unstable elements of the structure. To investigate the spatial distribution of residue stability, we selected five low cooperativity domains discovered by mHDX-MS to analyze by site-resolved hydrogen exchange NMR, along with three high cooperativity contrasting examples (**Fig. 3A-B**). In four of five low cooperativity domains, the unstable residues were clustered in specific regions of the structure (**Fig. 3C**). For example, the *de novo* designed protein HHH_rd4_0557 (magenta, family-normalized cooperativity -0.8 standard deviations (s.d.)) showed a tiered stability landscape for its three helices: α1 is the most stable, followed by α2, followed by α3. Only four residues in α3 had measurable protection by NMR, compared to nine residues in α1. The design HHH_rd4_0518 (blue, -0.6 s.d.) showed larger differences in stability between helices: the cores of α1 and α2 open near 6 kcal/mol, whereas the only measurably protected residues in α3 open below 3 kcal/mol (**Fig. 3C**). We solved the solution structure of HHH_rd4_0518 by NMR and found it was very similar to the designed structure and AlphaFold model, with the unstable α3 helix correctly folded in the native state (**Fig. 3D** and **Table S3**). This indicates that the faster exchange from α3 occurs through a (relatively low-energy) excited state, not the native state. As expected, the highly cooperative example HHH_rd3_0062 (orange, +2.0 s.d.) showed uniform opening energies across all three helices (**Fig. 3C**).

**Figure 3:**
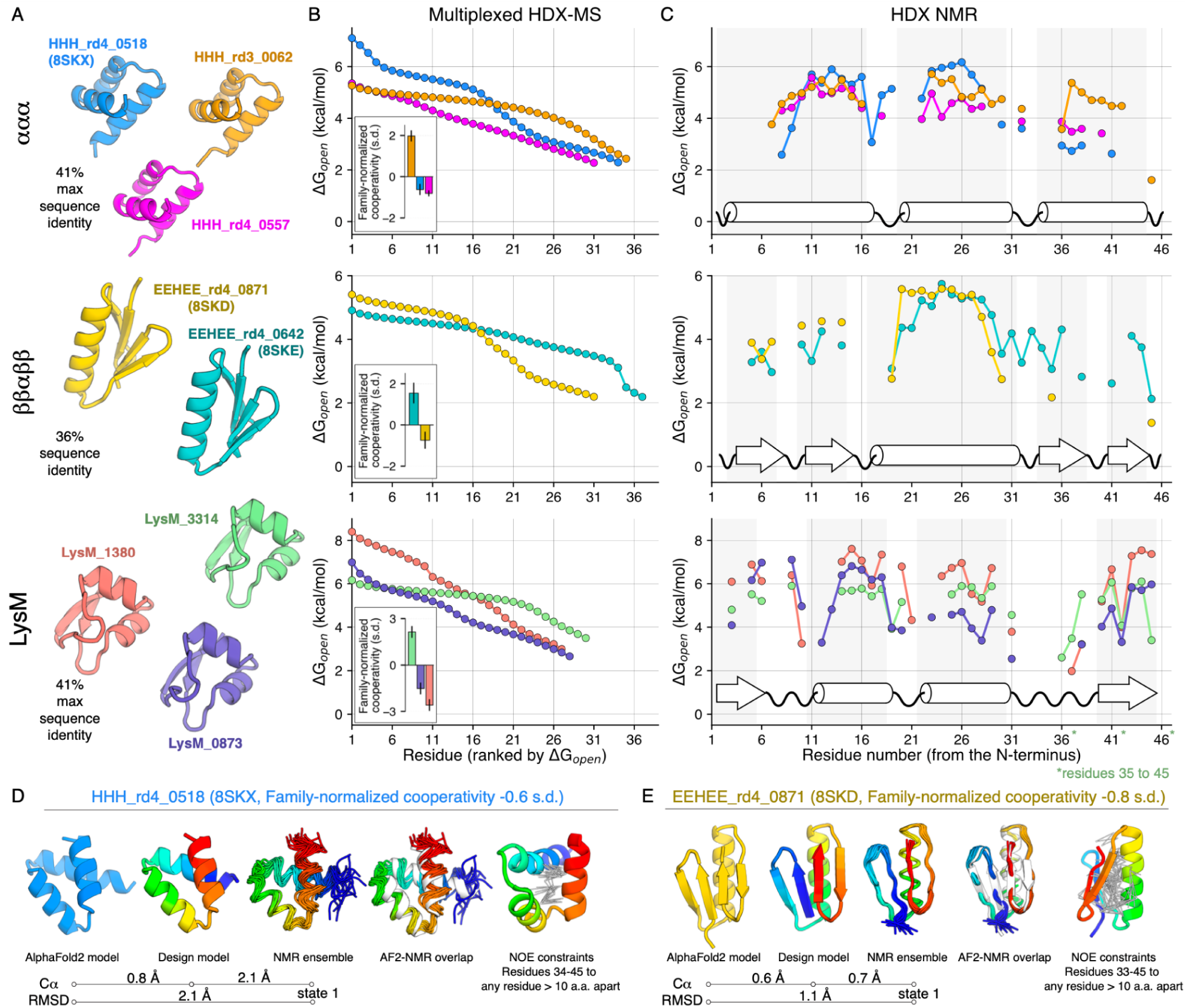
Low cooperativity domains show clustering of unstable residues. **A)** AlphaFold models of exemplar domains. **B)** mHDX-MS opening energy profiles for exemplar domains. Insets show family-normalized cooperativity with confidence intervals showing mean absolute deviations (MAD) from 8-15 replicates in different libraries for ααα and ββαββ domains and average experimental MAD across 36 proteins for LysM domains. **C)** Opening energies for exemplar domains from HDX NMR collected under multiple pH and temperature conditions as needed; see **Fig. S7** and **Table 1: Dataset_4**. Secondary structures are shown at bottom and shaded; some residues in secondary structure (helical first turns and edge-pointing strand residues) do not donate H-bonds and are not expected to be protected. For LysM_3314, residues 35-45 are plotted at sites 36-46 based on structural alignment to the other LysM domains. **D)** Solution NMR structure of the low cooperativity *de novo* protein HHH_rd4_0518 compared to the AlphaFold 2 and computational design models; in the NMR-derived structure, α3 adopts the correct fold and tertiary contacts despite its low stability. **E)** As in D, for the low cooperativity *de novo* protein EEHEE_rd4_0871. The C-terminal hairpin is correctly folded in the NMR ensemble, where it contacts the helix and N-terminal hairpin.

The low cooperativity designed protein EEHEE_rd4_0871 (**Fig. 3** yellow, family-normalized cooperativity -0.8 s.d.) and the low cooperativity natural domain LysM_0873 (**Fig. 3** dark blue, -1.5 s.d.) also showed clustering of low stability residues. In EEHEE_rd4_0871, the C-terminal β-hairpin is much less stable than the rest of the structure and is essentially unmeasurable by NMR (**Fig. 3C**). We solved the structure of both EEHEE_rd4_0871 and the high-cooperativity example EEHEE_rd4_0642 by NMR and both were similar to the designed models (**Fig. 3E** and **Table S3**). As with HHH_rd4_0518, the unstable C-terminal β-hairpin of EEHEE_rd4_0871 folds as designed, despite its low-energy fluctuations. In the low cooperativity natural domain LysM_0873, the unstable residues are clustered in α2, with lowered stability in β2 as well (**Fig. 3C**). The highly cooperative domains EEHEE_rd4_0642 (**Fig. 3** teal, +1.5 s.d.) and LysM_3314 (**Fig. 3** green, +2.1 s.d.) showed more uniform stabilities across all secondary structures, although the helix was more stable than either β-hairpin in EEHEE_rd4_0642.

The fifth low cooperativity natural domain LysM_1380 (**Fig. 3** red, -2.6 s.d.) did not show significant clustering of unstable residues. Instead, the variation within each secondary structure was greater than in the other LysM domains. Many residues in LysM_1380 (such as positions 6, 9, 10, 17, 27, 28, and 38) showed lower stability than the homologous positions in the other two LysM domains despite the higher global stability of LysM_1380 (**Fig. 3C**).

Overall, these results show that low opening cooperativity typically results from specific unstable elements of the structure - though not always. Bimodal Δ*G*_open_ distributions like those seen for HHH_rd4_0518, EEHEE_rd4_0871, and LysM_0873 may be evidence of spatial clustering of stability, but this need not always be the case. It is also important to note that residues sharing the same Δ*G*_open_ (below Δ*G*_unfold_) may open together or through different (but isoenergetic) partially open states. Finally, our results illustrate that highly stable small domains can still have relatively unstable structural elements. HHH_rd4_0518, EEHEE_rd4_0871, and LysM_0873 have relatively high global stability (Δ*G*_unfold_) compared to the other measured domains in their families (86%ile, 76%ile, and 94%ile respectively) despite the dynamics observed in their less stable elements.

### Structural determinants of cooperativity

To identify biophysical determinants of opening cooperativity, we used our modeled protein structures to calculate thousands of sequence and structural features for each domain. We then analyzed the Pearson correlation coefficient (PCC) between each feature and our experimental measurements of global stability and family-normalized cooperativity (**Fig. 4A**). Features included primary sequence properties (e.g. amino acid composition), physical energetic terms computed using Rosetta (Alford et al. 2017), and machine learning metrics from tools such as AlphaFold (Jumper et al. 2021), secondary structure predictors (McGuffin, Bryson, and Jones 2000), and disorder predictors (Hu et al. 2021; Redl et al. 2023). The large number of features makes it likely that some correlations would be observed by chance, but many observed correlations were stronger than those found using permuted data (**Fig. S13**). Overall, correlations were strongest in the ααα and ββαββ families, so we focused our analysis on these families (**Fig. 4A, Fig S14**). No individual feature dominated the correlations (maximum absolute PCCs with cooperativity 0.38±0.07 for ααα domains and 0.27±0.09 for ββαββ domains, mean±95CI from bootstrapping), indicating that cooperativity is influenced by many factors.

**Figure 4:**
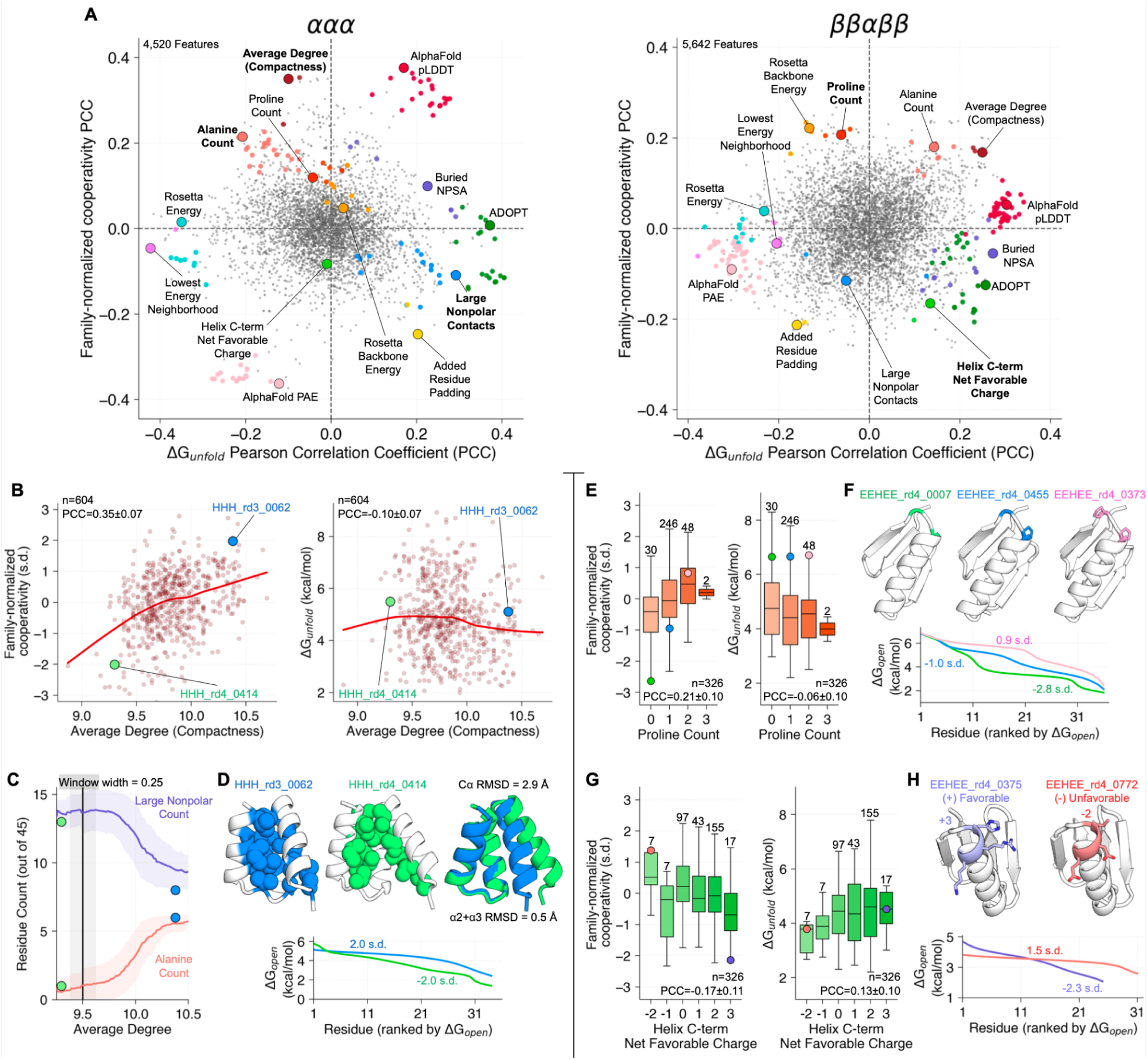
Protein features correlating with cooperativity. **A)** Pearson correlation coefficients (PCC) between protein features and family-normalized cooperativity (y-axis) or Δ*G*_unfold_ (x-axis), for ααα (left) and ββαββ (right) protein families. Large colored circles highlight notable features; small colored circles show features that are closely related to the same-color labeled feature (inter-feature PCC > 0.75); features in bold are highlighted below. **B)** Average degree compactness (average Cα count within 9.5 Å of each Cα) positively correlates with family-normalized cooperativity in the ααα family but has a small negative correlation with Δ*G*_unfold_. Large circles highlight two proteins shown in D. PCC 95% confidence intervals were computed by bootstrapping. **C)** Moving average (± std in shading) of alanine count (pink) and large nonpolar (FILMVWY) count (purple) across the range of average degree. Large circles show the two domains in D. **D)** Examples of high (blue) and low (green) compactness domains. Top: AlphaFold models with hydrophobic core residues highlighted. Bottom: Opening energy profiles and family normalized cooperativity. **E)** Proline count positively correlates with family-normalized cooperativity in the ββαββ family but negatively correlates with Δ*G*_unfold_. Circles highlight three proteins shown in F. Box plots are labeled with the number of examples in each bin. **F)** Example ββαββ domains with zero (green), one (blue), and two (pink) proline residues. Top: AlphaFold models. Bottom: Opening energy profiles and family-normalized cooperativity. **G)** Helix C-term Net Favorable Charge (HKR count minus DE count in the last three helical residues) negatively correlates with family-normalized cooperativity in the ββαββ family but positively correlates with Δ*G*_unfold_. Circles highlight two example proteins shown in H. **H)** Example ββαββ domains with three favorable (+) charges (blue) and two unfavorable (-) charges (red). Top: AlphaFold models. Bottom: Opening energy profiles and family-normalized cooperativity.

We identified protein features that correlated with cooperativity alone, stability alone, both, or neither (**Fig. 4A**). In both the ααα and ββαββ families, the Rosetta model’s “total energy” score (Alford et al. 2017), the amount of buried nonpolar surface area (NPSA), and the score from the ADOPT disorder predictor (Redl et al. 2023) all have relatively strong correlations with global stability, but relatively weak correlations with cooperativity (**Fig. 4A**). In contrast, the count of proline residues shows minimal correlation with folding stability, but shows one of the strongest correlations with cooperativity in the ββαββ family, and a similar (but relatively weaker) correlation with cooperativity in the ααα family. AlphaFold 2’s pLDDT confidence metric positively correlated with both stability and cooperativity in both families, but the correlation with cooperativity was stronger in the ααα family. Overall, features typically had directionally similar (but different strength) correlations between the two families.

Many features had opposite relationships with cooperativity and global stability. In the ααα family, the average degree compactness metric (average Cα count within 9.5 Å of each Cα) had one of the strongest positive correlations with cooperativity, but a negative correlation with global stability (**Fig. 4B)**. Alanine count also had a positive correlation with cooperativity but a negative correlation with global stability in the ααα family (**Fig. 4A**). This suggests an underlying tradeoff in the ααα family: highly compact domains are more cooperative, but this compactness is achieved through greater alanine content and fewer large nonpolar amino acids (**Fig. 4C,D**), which is modestly destabilizing.

In the ββαββ family, proline count had one of the strongest positive correlations with cooperativity but a negative correlation with global stability (**Fig. 4E**). Prolines were always located in the same two positions in the loop connecting the second strand to the helix (**Fig. 4F**). The side chains in the final helical turn also had opposing influences on stability and cooperativity. Helices are stabilized by positively charged side chains in the final turn that counteract the helical backbone dipole (Baker et al. 2015). We observed this trend across our 326 ββαββ domains: increased positive charge in the final turn showed a small positive correlation with stability (PCC 0.13±0.10, **Fig. 4G**). However, this was associated with a stronger negative effect on cooperativity (PCC -0.17±0.11, **Fig. 4G**, examples in **Fig. 4H**). This suggests that these charges primarily stabilized the helix with minimal cooperative increase in the stability of the sheet. If the helix has residual stability in the unfolded state (Mayor et al. 2003), this would also lower our inferred cooperativity by increasing the stability difference between the helix and the remaining residues.

These correlations provide insight into the biophysical determinants of opening cooperativity, but they depend on the domains in our dataset, which are not necessarily representative of all folded domains or all possible sequences. Correlations between different features (including correlations introduced by Berkson’s paradox and the minimal requirement of folding) also make it difficult to infer causal relationships. Still, the scale of our dataset enabled the discovery and quantification of correlates of opening cooperativity across a uniquely broad range of sequences. The consistency of many trends between the ααα and ββαββ families also suggests these trends may generalize further.

### Predicting stability and cooperativity

Individual feature correlations with global stability (Δ*G*_unfold_) and family-normalized cooperativity were generally modest, indicating that these properties are governed by the cumulative effects of multiple factors. To examine this, we built machine learning (ML) models to predict Δ*G*_unfold_ and family-normalized cooperativity for the four protein families with the most data (329–1,004 examples for the ββαββ, αββα, ααα, and LysM families). We used the engineered features from **Fig. 4** (including feature expansion and selection strategies, see Methods) as well as embeddings from protein language models (PLMs) to train regularized linear models (Lasso and Ridge regression) and evaluated these models using five-fold cross-validation. To improve model generality within each family, sequences were clustered by identity and clusters were assigned to the same fold (maximum identity between folds 30-55% for the four families).

The overall accuracy at predicting family-normalized cooperativity was relatively low: the best coefficients of determination (R²) for the four families ranged from 0.16 to 0.24 on unseen data (all cross-validation folds were used as separate hold-out sets for separate models and combined to calculate R², **Fig. S15B**). Still, these multi-feature models had higher correlations (even on unseen data) than the strongest single features examined in **Fig. 4A** (highest single-feature R² 0.05-0.14 without cross-validation for the four families). In contrast, models trained to predict Δ*G*_unfold_ using the same model architectures and training data were stronger, with R² values ranging from 0.40-0.53. This highlights the relative difficulty of predicting fine differences in energy landscapes compared to predicting global folding stability. Whereas language model embeddings led to the most accurate models at predicting stability (even for *de novo* designed families), models trained from manually engineered interpretable features (and explicitly modeled structures) led to the highest accuracy predictions of cooperativity for all four families (**Fig. S15B**).

### Designed mutations increase cooperativity

To understand sequence-cooperativity relationships on a more granular level, we examined how individual residues in individual proteins influenced opening cooperativity. We chose the proteins HHH_rd4_0518 and EEHEE_rd4_0871 as exemplars due to their characteristic low cooperativity from unstable C-terminal segments (**Fig. 3 A-C**), which is not intuitively explainable to our eyes. Both proteins have native structures that are fully folded and consistent with their computationally designed models and predicted structures (**Fig. 3 D-E**). For each protein, we used our family-specific models to identify double mutations predicted to increase opening cooperativity while preserving or increasing stability (**Fig. 5A**, **Fig. S16** and see Methods). Because mutations that increase both properties were predicted to be rare (only 6% and 4% of all mutations in HHH_rd4_0518 and EEHEE_rd4_0871 respectively, **Fig. S16**), identifying these mutations is also a substantial prospective challenge for these models. We selected 70 top-ranking double mutants from each wild type as our designed set for experimental testing. In EEHEE_rd4_0871, the most frequently designed mutations were spread across the whole protein (**Fig. 5A**). However, in HHH_rd4_0518, the most frequently designed mutations were clustered on the unstable C-terminal helix (**Fig. 5A**). The models never encountered spatially resolved opening energies, yet they inferred that mutating the least stable helix was the best route to increase opening cooperativity without sacrificing global stability. Along with these designed mutants, we also experimentally tested 70 randomly chosen double mutants of each wild type to analyze the impact of machine learning-guided design.

**Figure 5:**
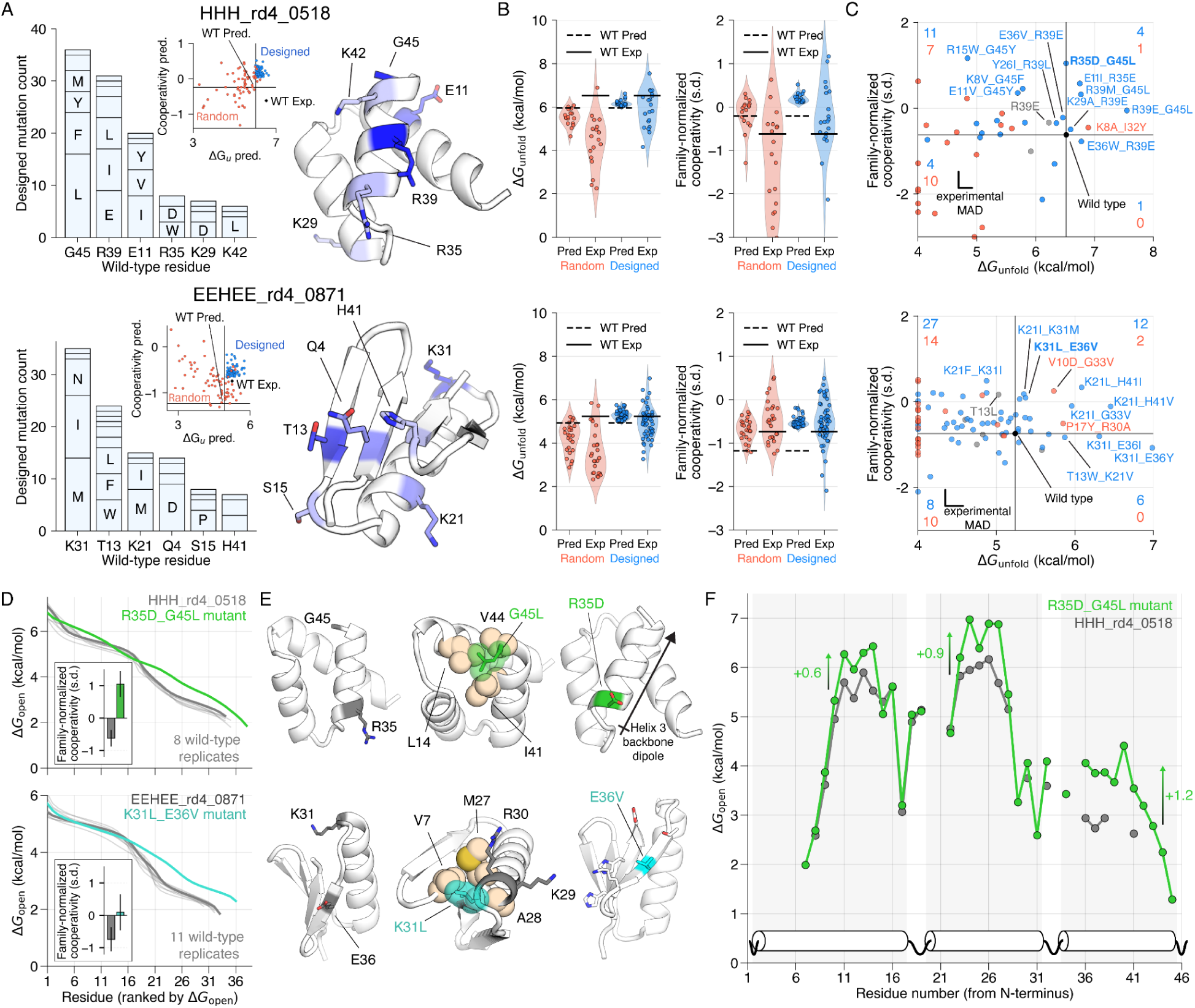
Data-driven design of improved cooperativity. **A)** Mutation frequency in designed double-mutant libraries of HHH_rd4_0518 (top) and EEHEE_rd4_0871 (bottom) for the six most frequently mutated sites (left). Structures (colored by mutation frequency) are AlphaFold models. Insets: Predicted Δ*G*_unfold_(x-axis) and family-normalized cooperativity (y-axis) for designed (blue) and randomly chosen (red) double mutants from linear machine learning models (see Methods). Lines show the predicted values for the (held-out) wild-type sequence from the same models; experimental wild-type values are marked by a black diamond. **B)** Comparison of predicted (Pred) and experimental (Exp) stability (left) and cooperativity (right) measurements for stable designed (blue) and random (red) double mutants of HHH_rd4_0518 (top) and EEHEE_rd4_0871 (bottom). Each wild-type sequence is shown as a black line. **C)** Experimental results of stable designed (blue) and random (red) double mutants of HHH_rd4_0518 (top) and EEHEE_rd4_0871 (bottom). Wild-type sequences are shown in black, single mutants in gray. Sequences outside the plot domain are shown on the edges. Each quadrant shows the count of designed and random mutants in that quadrant. Experimental mean absolute deviations (MAD) are shown and computed from replicates of other sequences (see *Methods* and **Fig. S5**). See **Fig. S17** for results organized by mutation. **D)** Opening energy profiles of double mutants with improved family-normalized cooperativity (colors). Replicates of wild-type measurements from different libraries are shown in gray with the highest quality measurement as a thick line. Inset: Family-normalized cooperativity for wild type and mutant sequences. Confidence intervals indicate MADs from replicates of the wild type in different libraries or the MAD across our whole dataset for the mutants. **E)** Structural effects of the mutations from D shown on AlphaFold structures; see main text. **F)** HDX NMR analysis of HHH_rd4_0518 (gray) and the G45L/R35D double mutant (green). Stability differences between the wild-type and mutant were calculated by averaging the three highest Δ*G*_open_ values in each helix.

We tested these 280 variants in small libraries due to the high sequence identity between mutants (108 sequences per library including internal standards) and successfully measured opening energy profiles for 38 HHH_rd4_0518 variants (20 designed, 18 random) and 80 EEHEE_rd4_0871 variants (54 designed, 26 random). Designed variants typically showed increased opening cooperativity, although this often came at the cost of lower global stability (**Fig. 5B**). Still, five HHH_rd4_0518 variants (four designed, one random) and 14 EEHEE_rd4_0871 variants (12 designed, two random) achieved simultaneous gains in both properties (**Fig. 5C,D**). In HHH_rd4_0518, mutations at R35, G45, and R39 (all in the unstable C-terminal helix) repeatedly yielded increases in both properties, as did mutations at K21, H41, and K31 in EEHEE_rd4_0871 (**Fig. 5C, Fig. S17**). In HHH_rd4_0518, the G45L mutation likely secures the C-terminus through multiple new hydrophobic interactions, while R35D/E converts an unfavorable interaction with the helix backbone dipole (Baker et al. 2015) into a favorable interaction (**Fig. 5E**). In EEHEE_rd4_0871, K31L likely stabilizes the hydrophobic core, E36V increases the strand propensity in the unstable C-terminal hairpin, and K21I/L introduces new hydrophobic contacts between the helix and the unstable hairpin, similar to contacts made with the N-terminal hairpin (**Fig. 5E**).

To examine how these mutations improved opening cooperativity at the residue level, we analyzed one variant of each protein by HDX NMR (**Fig. 5D,F**). In EEHEE_rd4_0871_K31L_E36V, we observed increased stability at the C-terminal end of the helix and in several residues of the unstable C-terminal hairpin, although most of the unstable hairpin still exchanged too quickly to resolve (**Fig. S18**). In HHH_rd4_0518_R35D_G45L, the two mutations stabilized the entire protein, with the greatest stabilization in the least stable helix α3 (+1.2 kcal/mol in α3 by comparing the three most stable residues between the mutant and wild-type, compared to +0.9 for α2 and +0.6 for α1, **Fig. 5F**). The number of measurably protected residues in α3 increased from four to eleven and included all H-bond-donating amides in the helix (**Fig. 5F**). The added stability in the least stable helix explains the improved opening cooperativity observed by mHDX-MS, although the added stability in the other helices (changes in the slowest rates) observed by NMR was larger than what we observed by mHDX-MS (see **Fig. S7** for complete comparisons of NMR-measured rates and MS-inferred rates).

Overall, these results illustrate how residues in unstable segments (such as R35D and G45L in HHH_rd4_0518) and outside unstable segments (such as E11 in HHH_rd4_0518 or K21 and K31 in EEHEE_rd4_0871) can increase or decrease local stability. As expected from the modest correlations, the machine learning results were not quantitatively accurate. However, they improved our exploration of sequence space by identifying rare mutations that could increase opening cooperativity while maintaining folding stability, with substantially improved success compared to random mutations. This illustrates the potential for rational data-driven engineering of protein energy landscapes.

## Discussion

It is widely appreciated that proteins are constantly in motion, continuously shifting between numerous conformational states on their energy landscapes. Yet the energetic details of these states — and the sequence features mediating them — remain almost entirely unknown. The mHDX-MS method makes it possible to experimentally analyze these fluctuations on a far larger scale than was previously possible, enabling new approaches to understand and predict fluctuations across sequence space. Our experiments revealed hidden variation in energy landscapes in both natural and designed domains, including substantial variation between related sequences. In fact, the variation within each fold family due to differences in sequence was often larger than the average difference between fold families (**Fig. 2E**). The low cooperativity domains highlighted in **Fig. 3** had entire pieces of secondary structure that were much less stable than the overall fold. These domains were not especially unusual: they represent the lowest 25%ile (HHH_rd4_0518), 22%ile (EEHEE_rd4_0871), and 6%ile (LysM_0873) opening cooperativity scores in their families. Hundreds of domains in our set showed similarly low opening cooperativity, and these instabilities remain largely invisible to methods that predict native protein structures. Analyzing these fluctuations across a wide sequence space revealed general protein properties that correlate with opening cooperativity, although our best models could predict only a limited fraction (16-24%) of the variance in opening cooperativity. Given experimental noise, we would expect perfect models to show correlations of R²=0.74-0.78, indicating how much remains to be discovered about the determinants of conformational fluctuations.

Our approach has important limitations. Our inferred rates depend on approximations about back exchange and the validity of combining data measured at different pH. Our inferred opening energies are also approximations because they depend on approximate chemical exchange rates (**Fig. 1E**: Eq. 2). Although our HDX NMR and cDNA display proteolysis measurements, data quality filters, and experimental reproducibility indicate that our high-throughput measurements are reliable, automated data processing can still introduce inaccuracies. To verify individual results, all raw MS data, extracted signals for each protein, and Bayesian fits are provided (see **Data Availability** and **Table S1: Dataset_8**), along with quality metrics for each step. The conformational details of our observed partially open states also remain unknown, and our multiplexed measurements do not localize the high- and low-stability segments of each domain. Resolving these details remains a key next step, especially for the hundreds of low-cooperativity domains discovered by our large-scale analysis. For the domains examined by NMR, the Δ*G*_open_ patterns of neighboring residues suggest that most openings involve entire secondary structures unfolding or fraying from the ends (Maity et al. 2003), but HDX experiments cannot establish this definitively.

Despite these limitations, we expect multiplexed HDX-MS to transform our ability to measure, predict, and model protein conformational fluctuations. Improved mass spectrometry technology will increase throughput even further. Larger domains should be amenable to our approach as well (Dumoulin et al. 2005). Combining our approach with bottom-up or top-down fragmentation strategies (Karch et al. 2018; Bhattacharjee and Udgaonkar 2021; Langford et al. 2024; Fang et al. 2023; Moroco et al. 2025; Filandr et al. 2024) should enable library-scale measurement with enhanced spatial resolution of conformational fluctuations. Profiling conformational fluctuations at scale will also reveal new links between these high energy states and protein function, aggregation, immunogenicity, and other critical properties. These fluctuations remain difficult to predict computationally (van Gunsteren et al. 2018; Orioli et al. 2020; Wan et al. 2020; Robustelli, Piana, and Shaw 2018), and large-scale mHDX-MS data offer a new route to directly optimize physics-based models (Robustelli, Piana, and Shaw 2018), machine learning potentials (Clementi et al. 2023; Majewski et al. 2023; Peng et al. 2022), and generative AI approaches (Lewis et al. 2024) to accurately model energy landscapes. The small proteins studied here are ideal for optimizing simulations due to the lowered computational cost, and our experiments uncovered a vast diversity of energy landscapes even amongst these structurally simple domains. Large-scale measurements, computational modeling, and machine learning make a powerful combination that have already transformed our understanding of protein native states (Jumper et al. 2021). The space of non-native states is far larger, and still remains elusive. Multiplexed HDX-MS offers a powerful approach for mapping this space to empower data-driven modeling, discover new biology, and accelerate protein engineering.

## Methods

### Library design

The initial set of 15,715 domain sequences was organized into five batches and further divided into 19 libraries (Mix 1-4, Lib 1,4 and 7-15, and mutants 2-4): a) Mix 1-4: De novo designed ααα, βαββ, and ββαββ sequences (Rocklin et al. 2017); b) Libraries 1 and 4: *de novo* designed αββα proteins (Kim et al. 2022); c) Libraries 7-14: Natural domains from the Pfam database, including LysM, PASTA, WW, SH3, Pyrin, and Cold-Shock; d) Library 15: PDB-derived monomeric proteins devoid of cysteine residues and metal cofactors. e) mutant libraries containing single and double mutants from EEHEE_rd4_0871 and HHH_rd4_0518 low cooperativity proteins. Sequences were randomly assigned to libraries within each batch, ensuring a minimum mass difference of 50 ppm between nearest-neighbor sequences for mass spectrometry compatibility (except Lib15 where two sequences are 36 ppm apart). After SUMO cleavage (see below), all proteins begin with the dipeptide HM (the scar from the NdeI ligation). Some sequences were modified with C-terminal padding (G, S, GG, or GS) to optimize mass spacing. All sequences were reverse-translated and codon-optimized for E. coli using DNAworks2.0 (Hoover and Lubkowski 2002). To standardize amplification efficiency, a ’GGS’ sequence was appended post-stop codon. Oligo libraries encoding the original 15,715 sequences were purchased from Agilent Technologies, while the 280 designed mutations were sourced from Twist Bioscience.

### Cloning of Twist oligo libraries into the pGR02 plasmid

Oligo libraries were resuspended and amplified via qPCR for restriction enzyme cloning. A preliminary qPCR run determined optimal amplification cycles, preventing overamplification by terminating reactions at ∼50% of maximum fluorescence intensity. Purified qPCR products were digested with XhoI and NdeI and ligated into the pGR02 plasmid, which encodes an N-terminal 10×His-SUMO tag. Ligated constructs were electroporated into 10-β Electrocompetent E. coli (New England Biolabs) and recovered in SOC medium at 37°C for 1 hour before plating onto selective MDAG-11+B1+K agar plates (Studier 2018). Serial dilutions determined transformation efficiency, and all colonies were pooled to maximize sequence diversity. Plasmid DNA was extracted from pooled cultures using the QIAprep Spin Miniprep Kit (Qiagen).

### Library expression and purification

Each library’s plasmid pool (5 μL) was electroporated into 25 μL BL21(DE3) Electrocompetent E. coli (Sigma-Aldrich), recovered in SOC media (1 mL, 37°C, 1 hour), and plated on selective MDAG-11+B1+K agar plates. Colonies were pooled and used to inoculate 2–4 L of LB broth with 50 μg/mL kanamycin. Cultures were grown at 37°C until OD600 = 0.6, then induced with 1 mM IPTG and incubated at 16°C overnight (∼16 hours). Cells were harvested via centrifugation and resuspended in lysis buffer (20 mM Tris, 500 mM NaCl, 30 mM imidazole, 0.25% CHAPS, 1 mg/mL lysozyme, 10 U/mL Benzonase, 1X Pierce Protease Inhibitor Cocktail, pH 8.0). Sonication (QSonica, 5 min total, 60% amplitude, 1 min on/off cycles) was followed by centrifugation (12,500 × g, 30 min, 4°C; repeated at 14,000 × g for clarification). The soluble fraction was purified via Ni-NTA agarose gravity columns (Qiagen). After washing with buffer (20 mM Tris, 500 mM NaCl, 30 mM imidazole, 0.25% CHAPS, 5% glycerol, pH 8.0), proteins were eluted (20 mM Tris, 300 mM NaCl, 500 mM imidazole, 5% glycerol, pH 8.0). Eluted proteins were dialyzed overnight into PBS, and SUMO tags were cleaved using a 1:100 molar excess of ULP1 (4°C, ∼20 hours). A second Ni-NTA purification removed SUMO and ULP1, collecting cleaved proteins in the flow-through. Proteins were concentrated (3 kDa Amicon Ultra filters) and further purified via Superdex 75 10/300 GL size-exclusion chromatography (Cytiva) on an NGC FPLC system (Bio-Rad). Monomeric fractions were pooled, re-concentrated, filtered (0.22 μm Millex-GP filter), flash-frozen in liquid nitrogen, and stored at -80°C until use.

### Labelled protein expression and purification for NMR analysis

We selected 13 proteins for individual expression, purification, and NMR analysis. The DNA sequences were codon-optimized for E. coli and cloned into pET-28a(+) (thrombin cleavage site) from Twist Biosciences or pET-28a(+)-TEV from GenScript. The plasmids were transformed into chemically competent BL21(DE3) cells. A small starter culture (5 mL) was inoculated in LB Miller broth with 50 μg/mL kanamycin and grown overnight at 37°C, 220 rpm. The starter culture (25 μL) was then diluted into 50 mL of labeled M9 media (42 mM Na2HPO4, 22 mM KH2PO4, 8.6 mM NaCl, 8.6 mM 15NH4Cl [Cambridge Isotope], 11 mM D-glucose [13C, Cambridge Isotope], 1 mM MgSO4, 0.2 mM CaCl2, 0.15 mM Thiamine, 1% [v/v] Trace Elements [3 mM FeCl3, 0.37 mM ZnCl2, 0.074 mM CuCl2, 0.042 mM CoCl2•H2O, 0.162 mM H3BO3, 6.84 mM MnCl2•H2O]) with 50 μg/mL kanamycin and grown overnight at 37°C, 220 rpm. Larger cultures of M9 media were inoculated with overnight M9 small culture (50 mL per 1 L) and grown at 37°C, 220 rpm to OD600 ∼ 0.6. Expression was induced with 0.5 mM IPTG, and cells were incubated at 16°C overnight (∼16-18 hours). Cells were harvested, resuspended in lysis buffer (20 mM Tris, 500 mM NaCl, 30 mM Imidazole, 0.25% CHAPS, pH 8.0, 1 mg/mL lysozyme, 10 U/mL Benzonase, 1X Pierce Protease Inhibitor EDTA-free), and lysed by sonication. The lysate was clarified by centrifugation (13,000 × g, 30 min). Proteins were purified by immobilized metal affinity chromatography (IMAC) using Ni-NTA agarose. The column was washed with buffer (20 mM Tris, 500 mM NaCl, 30 mM imidazole, 0.25% CHAPS, 5% glycerol, pH 8.0), and proteins were eluted in elution buffer (20 mM Tris, 300 mM NaCl, 500 mM imidazole, 5% glycerol, pH 8.0). Eluted proteins were dialyzed into buffer (50 mM Tris, 200 mM NaCl, 5% glycerol, pH 8.0) using Pur-A-Lyzer dialysis tubes (Sigma). His-tags were cleaved using either TEV protease (produced in-house, pRK793 plasmid; Addgene #8827) or Thrombin CleanCleave kit (Sigma), depending on the construct. TEV protease was added at a protease:target protein ratio of 1:10 with 0.5 mM DTT and incubated overnight at room temperature. Thrombin cleavage followed the manufacturer’s protocol, incubating overnight at room temperature. A second IMAC Ni-NTA purification was performed to remove the tag and uncleaved protein. Proteins were further purified by size-exclusion chromatography using a Superdex 75 10/300 column in phosphate-buffered saline. Monomeric fractions were identified based on elution profiles of a standard mixture (bovine serum albumin, ovalbumin, ribonuclease A, aprotinin, and vitamin B12), pooled, and concentrated using Amicon Ultra-4 Centrifugal Filters. Protein concentration was determined using the Pierce BCA assay (Thermo Fisher).

### Nuclear Magnetic Resonance (NMR) structure determination

NMR spectra for HHH_rd4_0518, EEHEE_rd4_0871 and EEHEE_rd4_0642 structure calculations were acquired at 288 K, on Briker spectrometers operating at 600 and 800 MHz, equipped with TCI cryoprobes with the protein buffered in 20 mM sodium phosphate (pH 7.5, 150 mM NaCl) at concentrations of 0.5 to 1 mM. Resonance assignments for ^15^N/^13^C-labeled proteins were determined using FMCGUI (Lemak et al. 2011) based on a standard suite of 3D tripe and double-resonance NMR experiments collected as described previously (Lemak et al. 2008). All 3D spectra were acquired with non-uniform sampling in the indirect dimensions and were reconstructed by the multi-dimensional decomposition software qMDD (Kazimierczuk and Orekhov 2011), interfaced with NMRPipe (Delaglio et al. 1995). Peak picking was performed manually using NMRFAM-Sparky (Lee, Tonelli, and Markley 2015). Torsion angle restraints were derived from TALOS+ (Shen et al. 2009). Automated NOE assignments and structure calculations were conducted using CYANA 2.1 (Güntert 2004).The best 20 of 100 CYANA-generated structures were refined with CNSSOLVE (Brünger et al. 1998) by performing a short restrained molecular dynamics simulation in explicit solvent (Linge et al. 2003). The final 20 refined structures comprise the NMR ensemble. Structure quality scores were performed using Procheck analysis (Laskowski et al. 1993) and the PSVS server (Bhattacharya, Tejero, and Montelione 2007).

### HDX NMR analysis

NMR HDX rates were measured for 13 proteins. A detailed overview of the experimental conditions, including buffer composition, temperature, pH, and protein constructs, can be found in **Fig. S7** and **Table S1: Dataset_4**. Exchange experiments were conducted at 600 MHz by tracking the decay of amide peak intensities in ¹H-¹⁵N HSQC spectra over a 24-hour period. Proteins were lyophilized in their respective analysis buffers, and exchange was initiated by dissolving them in an equivalent volume of D₂O. Each HSQC time point required approximately 5 minutes for acquisition, with the first measurement occurring ∼5 minutes after exchange initiation. Peak intensity data were fitted to a single exponential decay, and opening free energies were derived from these rates as described previously (Connelly et al. 1993; Nguyen et al. 2018; Bai et al. 1993).

### Multiplex hydrogen–deuterium exchange mass spectrometry (mHDX–MS)

Library samples (0.1 – 1 mg/mL) were diluted 1:9 into deuterated buffer (95% D₂O), either 25 mM MES (for pH_read_ ≈ 6) or 25 mM bicine (for pH_read_ ≈ 9). At specified incubation times, 95 µL of the exchange solution was mixed with 25 µL of quench buffer (0.5 M Gly–HCl, pH = 2.1-2.3). All sample handling was fully automated using a PAL3 LEAP HDxTool: incubation in D₂O buffer took place in a chamber at 20 °C, while quenching was performed in a chamber maintained at 0 °C. All buffers were individually adjusted prior to the experiment so that final pH_read_ values were 6.0 ± 0.05 (MES buffer) or 9.0 ± 0.05 (bicine buffer) and quench conditions were pH_read_ 2.45 ± 0.05 after diluting with the sample. The quenched samples were then analyzed on a Waters Synapt G2-Si Q–TOF mass spectrometer equipped with a Waters HDX Manager. Chromatographic separation was performed at 0 °C on a 1 mm × 100 mm BEH C4 column (300 Å, 1.7 µm particles) using a 30 min gradient, with the entire system maintained at 0 °C to minimize back exchange. Each library was measured in two batches - one at pH 6 and one at pH 9 - collecting 32 timepoints (log-spaced from 25 s to 24 h) plus three undeuterated replicates distributed throughout the experiment. All MS runs are performed in MS1 only mode at 1 scan per second. Every 10 seconds, one scan of Sodium Formate solution wes collected for usage in post processing calibration.

### Computational pipeline for mHDX-MS

We developed a computational pipeline for mHDX-MS analysis that is organized into two complementary repositories: “mhdx_pipeline” (available at https://github.com/Rocklin-Lab/mhdx_pipeline), which handles the processing of LC-IMS-MS data (converted from mzML) through protein identification, signal deconvolution via tensor factorization, and the assembly of time-dependent mass distributions using a path optimization module; and “hdxrate_pipeline” (available at https://github.com/Rocklin-Lab/hdxrate_pipeline), which applies back exchange correction, performs rate fitting via Bayesian optimization, and converts these rates into opening free energies. These pipelines are implemented using Snakemake (Köster and Rahmann 2012; Mölder et al. 2021), enabling reproducible, scalable, and parallel processing across compute clusters, thereby enhancing workflow efficiency and facilitating easy re-execution of the entire analysis. We also made available mhdx_analysis (https://github.com/Rocklin-Lab/mhdx_analysis) to provide readers with useful code for post processing of mHDX-MS results. Our entire codebase is freely available under CC BY 4.0 license. Below we describe the main components of both pipelines.

### Protein identification

We used IMTBX and Grppr to extract the set of protein-like isotopic clusters to allow for automated processing of the undeuterated samples (Avtonomov et al. 2018). We use default parameters except we don’t apply automatic mass correction. Calibration was performed as described below. Protein identification is performed solely based on the MS1 intact mass, with our library designed so that each protein exhibits a unique mass (> 50 ppm, except Lib15 where two sequences are within 36 ppm). In our pipeline, we have three sequential quality control steps for protein identification: step 1) mass error filtering: proteins are filtered based on their absolute protein monoisotopic mass error < 10 ppm of theoretical values; step 2) isotopic distribution matching: identified proteins are further filtered based on a dot product of experimental and theoretical isotopic distributions (idotp), with a threshold idotp > 0.98; step 3) we also implemented a false discovery rate (FDR) estimation and control: a target-decoy strategy to estimate and control for false discovery rate. In this strategy, we generate a decoy sequence database that is twice the size of the initial target library. Decoy sequences are sequences randomized within the expected mass range of target sequences (±50 Da) and are designed to be at least 50 ppm apart from their nearest mass neighbor. At this stage, proteins passing the mass error and idotp thresholds often include an (expected) greater number of target sequences compared to decoy sequences. This pre-filtered dataset is then used for FDR estimation. We trained a regularized logistic regression model to discriminate identifications from the target database from the decoy database. We extract 36 features, including charge-based features, ppm deviation, idotp, intensity metrics, and retention time (RT) residual errors. RT residual errors are calculated using a linear regression model trained on the target dataset to predict RT based on amino acid composition. The squared difference between experimental and predicted RT is used as an additional descriptor for our logistic regression model. The trained logistic regression model outputs probabilities for both the target dataset and the held-out decoy set. At each probability threshold, we estimate the FDR as the ratio of decoys to targets above the threshold. For individual identifications, q-values are calculated as the proportion of decoys relative to targets observed above the given probability threshold. Proteins are filtered based on a q-value threshold of 0.025, corresponding to an estimated FDR of 2.5%. In this work, steps 1 and 2 were applied as initial filters for downstream analyses, while step 3 served as a post-processing step. The regularized logistic regression model was trained using all library results (excluding mutant libraries) to derive a unified classification model. This model was then applied to mutant libraries, using the probability threshold (p = 0.803) observed to achieve an FDR of 2.5% in the training data.

### Mass spectrometry calibration

To ensure mass accuracy during the analysis, we implemented a lockmass calibration strategy using a reference compound. In this work, we used a solution with Sodium Formate (1:1:18 0.1 M NaOH:10% Formic Acid:Acetonitrile) as a calibrant. In our pipeline, raw mass spectrometry data files in .mzML format are parsed to extract individual scans and their respective retention times. For each scan, the experimental spectrum is filtered to retain peaks within a predefined mass-to-charge (m/z) tolerance of 100 ppm around theoretical reference masses. Filtered peaks with intensities exceeding 500 are grouped into chromatographic time bins (5 bins across a 30-minute runtime). This allows localized analysis of reference peaks to account for time-dependent variations. For each reference m/z value, a gaussian function is fitted to the experimental peaks in the filtered spectrum. Peaks with residual errors exceeding the defined 100 ppm tolerance or intensities below the threshold are excluded from calibration. For valid peaks, the observed m/z values are matched to the corresponding theoretical values. We build a linear regression curve to map observed m/z values to theoretical reference values. The calibration curve coefficients are stored and applied to correct all experimental m/z values within the dataset either at the stage of protein identification (see protein identification section) or at the tensor extraction stage (see tensor extraction section).

### Tensor extraction

To isolate individual protein signals from our liquid chromatography–ion mobility–mass spectrometry (LC–IMS–MS) data, we represented each experiment as a 3D tensor spanning retention time (RT), drift time (DT), and m/z dimensions. For each protein and charge state at each timepoint, we extracted a sub-tensor centered on the protein’s observed RT and DT (and window of ±0.4 min for RT, ±6% of DT center for DT, plus m/z range corresponding to the expected isotopic envelope extended to the maximum possible deuteration. RT and DT centers were empirically defined by averaging what was observed across undeuterated replicates to account for experimental variability. We applied m/z calibration corrections before extracting each sub-tensor to refine mass accuracy according to the strategy described above.

### Tensor factorization for signal deconvolution

We then performed iterative factorization via a rank-decomposition approach (e.g., nonnegative matrix factorization in multiple dimensions, see **Fig. S1A** for schematics, A. Marmoret 2020). We smoothed the sub-tensor in the RT and DT dimensions using small Gaussian kernels (σ_RT_,σ_DT_=(3,1)) to improve signal-to-noise ratio. Beginning with an initial guess of *k*=5 factors, if the minimum correlation across RT, DT or m/z dimensions between any pair of factors exceeded a specified threshold of 0.17, we considered the data over-factorized and reduced *k* accordingly. This adaptive process continued until factors were sufficiently distinct. If factors were initially sufficiently distinct, we increased *k* until we observed over factorization, keeping the previous iteration. Each resulting factor was further filtered by their Gaussian fit quality in RT and DT (R^2^ ≥ 0.90) to ensure they represented coherent elution or drift profiles. Next, we examined each factor for multiple isotopic clusters. We projected each factor’s m/z dimension to create an integrated mass profile, then used a peak-finding algorithm to locate individual clusters. If more than one cluster was detected, we split the factor accordingly, treating each isotopic cluster as a distinct signal component. This procedure allowed us to separate closely spaced isotopic envelopes and reduce noise-induced over-segmentation, ultimately yielding the collection of signals for downstream analyses.

### Path optimizer for assembly of time course mass distributions

After extracting and deconvoluting signals for each protein and timepoint, we assembled a consistent, time-resolved mass profile for each protein by selecting the most plausible isotopic cluster (IC) at each timepoint using the Path Optimizer (PO) module in our pipeline. The resulting “path” spans from the undeuterated (initial) to increasingly deuterated (late) states. All ICs - regardless of their charge state - are analyzed together by converting each m/z signal into a neutral mass representation (i.e., baseline-integrated mass distribution). Before path optimization, we derive nine quantitative features for each IC: 1) RT/DT errors: we compute the deviation in each IC’s retention time (seconds) and drift time (percentage) relative to the undeuterated reference signal; 2) RT/DT profile similarity: the cross-correlation between each IC’s elution profile (RT or DT) and that of the undeuterated reference quantifies overall shape similarity; 3) Peak error: we compare the observed mass to a theoretical isotopic envelope, penalizing large discrepancies from the expected mass; 4) full width at half maximum (FWHM) deviation: we calculate the FWHM difference relative to the undeuterated reference; 5) Intensity deviation: each IC’s integrated intensity is compared to the baseline intensity of the undeuterated protein; unusual gains or losses raise suspicion; and 6) Neighbor correlation: if a protein exhibits multiple charge states at a given timepoint, we compute a correlation across m/z and RT dimensions to ensure they represent the same underlying species. ICs lacking other charge states are assigned zero correlation. 7) Signal noise estimation: the integrated mass distribution is fitted to a Gaussian, higher noise or incomplete peaks indicate lower quality, and result in higher values.

To reduce the number of low-quality or redundant isotopic clusters (ICs) before optimization, we apply two complementary filtering strategies: 1) User-defined thresholds: each IC is evaluated against empirical cutoffs for multiple features (e.g., maximum RT/DT errors, minimum RT/DT cross-correlation, maximum mass error). ICs exceeding these thresholds (or failing to meet minimum criteria) are excluded outright, ensuring only clusters within typical experimental bounds proceed to the next stage. 2) Weak Pareto dominance: within each timepoint, we compare ICs that have similar baseline-integrated neutral masses. If one IC is strictly worse than another on multiple metrics (e.g., higher RT/DT errors, lower RT/DT profile fits, greater peak error), it is Pareto-dominated and removed. This pruning further refines the set of plausible ICs by discarding those demonstrably inferior across key features.

After prefiltering, we create an array of hypothetical deuteration trajectories, each characterized by a starting fractional uptake and a slope parameter in logarithmic space. For each trajectory, our algorithm selects the best undeuterated IC (based on the dot product with the theoretical isotopic distribution, idotp) and next proceeds across subsequent timepoints, selecting the IC whose baseline-integrated mass is closest to the trajectory’s expected deuteration level. Each of these initial sampled paths undergoes a greedy local refinement search where we iteratively swap out a single IC at a time if a replacement lowers the path score. Specifically, at each timepoint, we consider all ICs and check whether substituting any single IC yields a better overall path. This iterative process continues until no single substitution can further reduce the path score. Each potential time-course path is evaluated using a multidimensional scoring function which sums several penalty terms that capture low data quality or physically implausible behavior. The physically implausible behavior is assessed whether mass addition between consecutive timepoints does not decrease (no negative uptake) and does not change abruptly (average deuterium uptake doesn’t become faster with time). From the final paths, after the greedy swaps, we choose the one with the lowest score as the winner. This path represents the most coherent set of ICs from undeuterated through late timepoints.

Before downstream analysis, the pipeline runs the PO module in a temporary mode, generating a set of best-fit paths from all proteins to collect statistics (e.g., typical RT/DT error, baseline mass deviations). These statistics are aggregated to define empirical thresholds (e.g., two standard deviations above/below the mean) that exclude outlier signals. For each protein, the pipeline then repeats path optimization as described above with these thresholds in non temporary mode, ignoring candidate clusters failing the newly established criteria. This two-stage approach filters low-quality clusters and yields more robust final timecourses. Final paths with path score (PO total score) lower than 50 were selected for rate fitting.

### Back exchange correction in mHDX-MS

In HDX-MS, deuterated residues can “back exchange” to hydrogen during the quenching step and during liquid chromatography (performed in a non-deuterated buffer). The level of back exchange varies from protein to protein (affecting all measurements of the same protein in a uniform way) and also varies timepoint to timepoint based on small differences in conditions (affecting all proteins in the sample in a uniform way). We correct all measurements using both timepoint-specific and protein-specific corrections to determine the original level of deuteration before any back exchange. Although different residues in each protein may back exchange at different rates, our model assumes a single overall back exchange percentage for all residues in a given protein at a given timepoint (**Fig. 1E**: Eq. 3).

#### Protein-speciifc back exchange correction (Fig. S2A)

We determine the percentage of back exchange for each protein based on the deuteration level observed in the longest timepoint (typically 24h) when the total deuteration is no longer changing (i.e. the protein achieved full deuteration and any missing deuteriums are the result of back exchange). Back exchange percentages are computed separately for pH_read_ 6 experiments and pH_read_ 9 experiments for each protein. For proteins that do not reach full deuteration in pH_read_ 6 experiments, we estimate their pH_read_ 6 back exchange level based on an empirical linear correlation between back exchange measured at pH 6 (for other, fully deuterated proteins) and back exchange measured at pH_read_ 9 (for those same proteins, **Fig. S2A**). Overall, protein-specific back exchange varied from 6-45%.

### Timepoint-specific back exchange correction (Fig. S2B)

We identify “fully deuterated” proteins by checking whether their final five timepoints show ≤1 Da variation in mass. Next, we check which subset of proteins are fully deuterated in the preceding timepoints by checking if their centroid masses remain within 6% of that final mass. Owing to the diverse stabilities in our samples, there is typically a subset of unstable proteins that rapidly reach full exchange, serving as internal controls for timepoint specific back exchange across all timepoints. If fewer than five such proteins are found at a given timepoint, we apply the average back exchange computed from later timepoints where enough fully deuterated proteins were found. The timepoint-specific back exchange correction (which varied from -5% to +4% across all timepoints) is added to each protein’s back exchange value to determine Backexchange(*s*,*t*) in **Fig. 1E**, Eq. 3.

### Inferring exchange rates from isotopic distributions

We sample a set of exchange rates *k*_HX_ (one rate per exchangeable residue) to obtain the posterior rate distribution that is consistent with the observed isotopic envelopes according to the model described in **Fig. 1E**, Eq. 3 and 4. The N-terminal two residues are excluded because these fully back exchange. For domains where we combine data from pH_read_ 6 and pH_read_ 9, we also sample the time scaling factor which determines “effective pH_read_ 6 measurement times” for all pH_read_ 9 measurements (**Fig. S2 C-F**). This allows for small deviations from a theoretical factor of 10^3^ due to pH-dependent changes in protein dynamics or charge effects on *k*_chem_.

The inputs to the model are the experimentally measured integrated mass distributions at each timepoint (non-zero intensity only), the corresponding timepoints, the theoretical isotopic distribution in the undeuterated state (Gohlke 2022), the estimated level of back exchange at each timepoint, and the fraction of D₂O in the exchange buffer. Next, we use Bayesian inference (via No-U-Turn Sampling, NUTS, as implemented in the numpyro package, Phan et al. 2019) to infer the set of amide exchange rates ln(*k*_HX_). We define a hierarchical prior that first samples a “slowest rate” from a truncated normal centered near e^-7^ s^-1^ (with scale e^10^ and bounded below e^-15^), then linearly spaces a set of *N* log-rates from this slowest rate up to ln(e^10^). Each of the *N* log-rates is assigned a broad normal prior distribution (σ = 5), reflecting minimal prior knowledge (**Fig. 1F**). For each proposed set of rates, we compute a Poisson-binomial (Hong 2013; Straka 2017) distribution of exchanges - adjusted by the inferred back exchange and deuterium fraction (**Fig. 1**, Eq. 3) - to yield a convolved “theoretical” integrated mass envelope per timepoint. Discrepancies between these theoretical envelopes and the measured intensities are captured by a Gaussian distribution with a global noise parameter (drawn from an exponential prior, σ = 1). We run the Markov chain Monte Carlo (MCMC) algorithm with four parallel chains, using 100 warm-up iterations and 250 posterior draws in each chain. Post-sampling, we discard any problematic chains (e.g., poor R-hat or chain-specific RMSE) and re-run if necessary. Finally, samples from all chains are combined, and each sample’s posterior rate distribution is sorted from slowest to fastest, yielding posterior distributions for each *i*’th slowest rate (with *i* ranging from 1 to *N*) as shown in **Fig. 1F**.

### Chemical intrinsic rates approximation in mHDX-MS analysis

To convert our Bayesian-inferred exchange rates *k*_HX,*i*_ into residue opening free energies Δ*G*_open,i_, we must know each site’s intrinsic rate *k*_chem,*i*_ (**Fig. 1E**, Eq. 2). However, each residue has a unique *k*_chem,*i*_ based on its local sequence context (Pedersen et al. 2019; Nguyen et al. 2018), and our data do not reveal which measured rate is associated with which residue’s *k*_chem_. A straightforward approximation is simply to use the median (or mean) value among the intrinsic rates. To improve on this approximation, we examined the empirical relationship between *k*_chem,*i*_ and *k*_HX,*i*_ using site-resolved HDX NMR data on 11 proteins (all from **Fig. S7** except double mutants HHH_rd4_0518_R35D_G45L and EEHEE_rd4_0871_K31L_E36V). We found a weak but significant trend across nearly all proteins whereby residues with slower *k*_HX,*i*_ also had slower *k*_chem,*i*_ (**Fig. S3A**). We incorporated this trend into our estimation of *k*_chem,*i*_ using a simple scaling factor between normalized *k*_HX,*i*_ and normalized *k*_chem,*i*_:

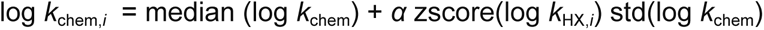

where median(log *k*_chem_), zscore(log *k*_HX,*i*_), and std(log *k*_chem_) are computed separately for each protein, and a universal scaling factor *α* is used for all proteins. As a result, a residue with the average log *k*_HX,*i*_ for its protein (i.e. zscore = 0) will be analyzed using that protein’s median *k*_chem,*i*_, and faster or slower residues will be analyzed using faster or slower *k*_chem,*i*_ based on the scaling factor *α*. Here, we used a scaling factor *α* = 0.38, which nearly minimizes the error between our inferred Δ*G*_open_ distributions and the Δ*G*_open_ distributions from NMR across all 11 proteins (this was computed using a smaller set of NMR proteins than our final set). This adjustment yields Δ*G*_open_ distributions that are more accurate than assuming the median *k*_chem_ for all positions (**Fig. S3B**).

### Net charge correction to all Δ*G*_open_

To account for electrostatic effects on hydrogen exchange rates, we applied a correction to the estimated Δ*G*_open_ values based on the estimated net charge of each protein at pH 6 (as computed by ProtParam module within the Biopython package, Cock et al. 2009; Gasteiger et al. 2005). It has been shown that negatively charged proteins have slower *k*_chem_ due to decreased local concentration of the hydroxide catalyst, and vice versa (Shaw et al. 2010; Dass et al. Mulder 2021), although these corrections are not explicitly considered in the exchange framework of (Bai et al. 1994). Our data reproduced this electrostatic dependency. To empirically correct for this effect, we derived a linear adjustment that removes the dependency between net charge and Δ*G*_unfold_. For proteins with Δ*G*_unfold_ > 4 kcal/mol, we found the nearly optimal correction coefficient of 0.12 kcal/mol per unit charge, which was then applied to all Δ*G*_open,i_ values.

### Δ*G*_unfold_ calculation

We derive Δ*G*_unfold_ as the average across the five more stable residues (averaging across 1-6 residues leads to highly correlated Δ*G*_unfold_ estimates, **Fig. S4A**). When deriving Δ*G*_unfold_, we did not account for the contribution of the unfolded state cis-trans isomerization to the measured Δ*G*_unfold_ (Huyghues-Despointes, Scholtz, and Pace 1999; Bai et al. 1994). Thus, all Δ*G*_unfold_ measurements are in reference to an unfolded state where prolines have their native-state isomerization state, rather than a fully-equilibrated unfolded state. However, analysis of proline isomerization states in AlphaFold 2 models (**Fig. S4B**) indicates that cis-prolines are predominantly present in PASTA domains, with minimal occurrence in other protein families analyzed (**Table S1: Dataset_3**). Therefore, potential biases due to cis-prolines are largely confined to PASTA and do not substantially affect the other protein families examined. We also did not correct results for the small effects of trans-prolines.

### Detecting and filtering EX1 Behavior

Our pipeline is primarily designed for unimodal, EX2-like exchange profiles. It attempts to split multimodal or poorly fitted peaks into separate unimodal peaks, which is not ideal for analyzing EX1 kinetics. We therefore do not recommend using this code, in its current form, to analyze EX1 data. Nonetheless, we implemented an automated procedure to identify and exclude these potential EX1-type profiles from our dataset. Under EX1 kinetics, backbone amides often exhibit an abrupt shift in their isotopic distribution, leading to characteristic anomalies in the experimental and fitted exchange profiles. (1) Width criterion: for each timepoint, we compared the observed isotopic distribution width to the width of the theoretical distribution from our EX2-only fit. We then computed the ratio of these widths across the exchange progress—specifically focusing on timepoints where the majority of amides (80–90%) had undergone exchange. Samples were flagged as potential EX1 if two conditions were met: (i) the average width ratio in that progress window exceeded 1.25, and (ii) the maximum ratio was greater than 1.35. Any domain meeting both thresholds at either pH 6 or pH 9 was classified as ex1_width_criteria. (2) Jump criterion: To capture large, discrete transitions in the exchange profiles, we tracked the difference between the centroid of the fitted distribution (EX2 assumption) and the maximum peak of the observed distribution over time. An exchange profile was labeled as ex1_jump_criteria if it displayed a shift exceeding a predefined range (from less than -2Da to more than +2Da) coupled with the transition happening below 90% overall exchange. These constraints were set to identify “jumps” characteristic of EX1-like coordinated exchange rather than the smoother transitions expected under EX2 kinetics. See **Fig. S4** for schematics on each criteria.

### Determining full deuteration and protein classification in mHDX-MS

For protein classification, we defined full deuteration as a delta mass of less than 2 Da between the back exchange corrected centroid masses measured at the last and fifth-to-last timepoints. Based on this criterion, our data are classified into four groups: (1) unmeasurable proteins, for which full deuteration occurs too early for accurate exchange rate modeling. We also classify a unmeasurable protein if any of following conditions are met: slowest rate constant greater than 10⁻⁴ s⁻¹, Δ*G*_unfold_ value is lower than 2 kcal/mol, or fewer than 20% of residues present measurable exchange rates (k_HX_ < 10^-3.45^ s⁻¹); (2) proteins that achieve full deuteration in the pH 6 experiment; (3) proteins that only reach full deuteration in the pH 9 experiment - requiring the integration of data from both pH 6 and pH 9; and (4) proteins that do not reach full deuteration even in the pH 9 experiment. Proteins belonging to groups (2) and (3) are subsequently referred to as measurably stable.

### Filtering low-quality data

To maximize the reliability of our mHDX-MS measurements, we applied a series of filtering criteria (**Fig. S19**). First, low-quality identifications were removed based on mass error (> 10 ppm), isotopic distribution matching (idotp < 0.98), and q-values (> 0.025) and paths with a path optimizer score above 50 were excluded (or above 40 for HHH_rd4_0518 and EEHEE_rd4_0871 mutants) (**Table S1: Dataset_1**). For all proteins that reach full deuteration, we require overall back exchange to remain below 45% (we do not consider back exchange for proteins in Group 4 which do not reach full deuteration). For proteins fully deuterated under both conditions, we required that the back exchange difference between pH 6 and pH 9 be less than 10%. Additionally, we assessed the consistency between the experimental and computed mass distributions (from the inferred rate constants) by requiring that 90% of timepoints have an RMSE lower than 0.3. This filtered dataset is available as **Table S1: Dataset_2**. Finally, for each protein, the mass distribution path with the lowest path optimizer score was selected as the representative opening energy distribution, and proteins exhibiting EX1 kinetics (**Fig. S6**), unmeasurable protection, or incomplete deuteration at pH 9 (**Fig. S11**) were excluded from the measurably stable dataset (**Table S1: Dataset_3**).

### Protein structure prediction

We predict the 3D structures of the proteins from their amino acid sequences using AlphaFold 2 (Jumper et al. 2021). The protein sequences, provided in FASTA format, were used as input to the ColabFold implementation of AlphaFold 2 (Mirdita et al. 2022), which was run on Quest high performance computing facility at Northwestern University. The pipeline generated five models for each sequence, and the best-scoring model, based on the predicted pLDDT, was refined using amber. The best AlphaFold model for each sequence was further refined using the Rosetta Relax protocol (Conway et al. 2014). This process aimed to optimize the predicted structures by minimizing their energy. When not explicitly mentioned otherwise, the relaxed structure was selected for downstream analysis (**Table 1: Dataset_9**).

### Deriving normalized cooperativity and family**-**normalized cooperativity

We first computed each protein’s average opening free energy (Δ*G_a_*_vg_) as the average of all Δ*G*_open_ values, with unmeasurably fast residues set to a lower bound of 0 kcal/mol. Here, Δ*G*_avg_ serves as a simplified proxy for the energies of partially open states on the energy landscape. We then built an empirical model of the typical average opening energy (Δ*G*_avg,expected_) for a given domain based on its global stability (Δ*G*_unfold_), its fraction of exchangeable backbone amides that donate hydrogen bonds (fxn_hb), and its net charge (netq). In our Bayesian regression model, we define the expected average free energy as

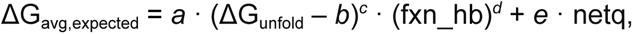

where the parameters *a*, *b*, *c*, *d*, and *e* are sampled from non-informative priors (specifically, *a* ∼ Normal(1, 3), *b* ∼ Normal(0, 5), *e* ∼ Normal(–1, 3), log(*c*) ∼ Normal(–0.3, 2) with *c* = exp(log(*c*)), and d ∼ Normal(–0.3, 2) with *d* = exp(log(*d*))), and the observation noise is modeled with σ ∼ Exponential(1). We run Markov chain Monte Carlo using NUTS (as implemented in numpyro; Phan et al. 2019) with 1000 warm-up iterations and 1000 samples to obtain the posterior distribution of these parameters. We then use the mean values of the posterior distributions for the parameters *a*-*e* (**Table S2**) to compute Δ*G*_avg,expected_ for each domain. Finally, we define normalized cooperativity as the z-score of the residual between each domain’s experimental Δ*G*_avg_ and its predicted value, Δ*G*_avg,expected_.

The hydrogen bonding fraction fxn_hb is defined as the fraction of backbone amides that donate hydrogen bonds to any acceptor (sidechain or backbone) in the molecule. The first two residues (the N-terminus and the first amide) are excluded from both the numerator and the denominator because these residues fully back exchange during HDX-MS. Hydrogen bonds are determined using the Rosetta model (Alford et al. 2017; Chaudhury, Lyskov, and Gray 2010) based on AlphaFold 2 structural models that are subsequently relaxed with Rosetta. We call these predicted structures *reference structures*.

These reference structures are critical for the calculation of normalized cooperativity because they define our expectation for how many protected residues should be observed in a fully cooperative domain. Even if a domain’s native structure under our experimental conditions differs from the reference structure (which may occur because the reference structures are only computational predictions), normalized cooperativity is still computed based on the number of hydrogen bonds in the reference structure. In an extreme case, a protein might be substantially unfolded under our experimental conditions compared to the reference structure. For example, consider a protein with two subdomains where both are folded in the reference structure but only one subdomain is stable under our experimental conditions. Here, “cooperativity” could be interpreted in different ways. If we only consider the fraction of the structure that is stably folded under our experimental conditions, this subdomain might behave in a perfectly two-state manner and be considered highly cooperative. However, *the domain as a whole* (including both subdomains) is not cooperative because one subdomain can unfold while the other remains folded. Our analysis (based on the reference structure) considers the domain as a whole. In this scenario, Δ*G*_avg_ will be low, because the residues in the unfolded subdomain (with low Δ*G*_open_) are included in the average. Δ*G*_avg,expected_ will be much higher, because Δ*G*_avg,expected_ is computed assuming a larger number of protected residues than are experimentally observed. The result is a low normalized cooperativity, indicating that the full domain (according to the reference structure) is relatively low cooperativity compared to other domains with similar stability in our dataset.

We also include a linear correction for net charge (netq) in the calculation of Δ*G*_avg,expected_. As discussed in **Methods: Net charge correction to all Δ*G*_open_**, negatively charged proteins tend to exhibit slower *k*_chem_. As a result, negatively charged proteins tend to have a larger fraction of measurably stable residues compared to positively charged proteins (exchange is uniformly slower in negatively charged proteins regardless of conformational stability, so more residues can be measured). Although we correct for this bias by applying a charge correction to all Δ*G*_open_ measurements, there is still an uncorrected bias on the number of measurable residues. In fitting *k*_HX_, we use a prior distribution on *k*_HX_ that assumes that most residues are very unstable until they are shown to be stable in the data (**Fig. 1F**). As a result, the inferred *k*_HX_ (and Δ*G*_open_) end up being very different for residues that are barely slow enough to be measurable and residues that are barely too fast to be measured. Residues that are just barely measurable end up with Δ*G*_open_ ∼2 kcal/mol, compared to residues that are slightly too fast to be measured, which often have inferred Δ*G*_open_ much lower than 0 kcal/mol (**Fig. 1F**). These very low opening energies are set to the lower boundary of Δ*G*_open_ = 0 kcal/mol when calculating Δ*G*_avg_. As a result, the bias toward more measurable residues in negatively charged proteins ends up biasing Δ*G*_avg_. Including netq in Δ*G*_avg,expected_ mitigates this bias to enable a more fair comparison of normalized cooperativity in negatively and positively charged proteins.

We derived two distinct cooperativity metrics. The “normalized cooperativity” metric is obtained by fitting the model to the entire dataset, while the “family-normalized cooperativity” metric is derived by fitting the model separately within each protein family (using only families with more than 50 examples in the measurably stable category). For both metrics, we report their normalized versions—computed by subtracting the mean and dividing by the standard deviation of the residuals—yielding the normalized cooperativity and family-normalized cooperativity, respectively. A Jupyter Notebook for modeling cooperativity from new data, along with a script to derive cooperativity using our precomputed metrics, are available at https://github.com/Rocklin-Lab/mhdx_analysis.

### Reproducibility of opening energy distributions in mHDX-MS

Starting from the filtered dataset (**Table S1: Dataset_2**), we assessed the reproducibility of measured opening energy distributions across independent mHDX-MS experiments. First, for proteins analyzed in multiple libraries, if a protein was represented by more than one opening energy distribution within a given library, we selected the path with the lowest path optimizer (PO) score as the representative for that library. Across all libraries, 127 opening energy distribution replicates were identified for 36 different proteins. For each protein, the replicate with the lowest PO score was designated as the reference distribution, and the mean absolute deviation (MAD) was computed between the reference and every other replicate for that protein in a different library, considering only the measurable opening energies (i.e. excluding energies derived from rates outside the dynamic range). This MAD is shown in **Fig. 1G**.

We further observed that some proteins were detected at multiple retention times within a single experiment. To evaluate the reproducibility in this context, we started with the opening energy distribution corresponding to the lowest PO score as the reference. We then iteratively added additional distributions only if they were acquired at retention times at least 2 minutes apart from any previously selected distribution—thereby ensuring that the additional replicates reflected genuine distinct chromatographic events rather than minor experimental variability. Under these criteria, 180 proteins exhibited multiple distinct retention time replicates (**Fig. S18A**). Across these replicates, both the MAD for global stability (Δ*G*_unfold_) and the MAD computed over all measurable Δ*G*_open_ values were less than 0.5 kcal/mol (**Fig. S18B–D**).

### Comparison between mHDX-MS and HDX NMR

We compared exchange rate and opening energy measurements obtained from HDX NMR with those derived from mHDX-MS. For the rate comparison, each HDX NMR log-rate was compared to a simulated mHDX-MS log-rate, which was derived by adjusting the measured mHDX-MS rates by the ratio of the median *k*_chem_ values between the two conditions. The overall agreement between the two methods was quantified by computing root mean squared error (RMSE) across all 511 *k*_HX_ measurements collected from 13 proteins in multiple HDX NMR conditions (**Fig. S7**). For the opening energy distributions, each construct’s sorted average Δ*G*_open_ (averaged over residues when multiple HDX NMR conditions were available for the same residue) was compared to the corresponding distribution from mHDX-MS. In cases where the two distributions differed in length, both distributions were truncated to the length of the shorter distribution. We then computed a single overall RMSE across all 323 Δ*G*_open_ measurements from the 13 proteins.

### Feature extraction

Our feature extraction pipeline leverages both interpretable features and protein language model embeddings. For the interpretable features, we computed a comprehensive set of predictors that include sequence-based descriptors generated with iFeature (Chen et al. 2018) (e.g., amino acid composition, grouped amino acid composition, and composition–transition–distribution metrics), disorder predictors (ADOPT, (Redl et al. 2023) and flDPnn (Hu et al. 2021)), and structure-based features derived from AlphaFold2/Rosetta-relaxed models. These structure-based features encompass pLDDT scores from AlphaFold2, a set of Rosetta-based features extensively described previously (Rocklin et al. 2017), solvent accessibility (Mitternacht 2016), as well as custom-derived metrics such as side-chain contact scores, burial scores, hydrogen-bond metrics, and an expanded set of secondary structural element-specific descriptors (e.g., maximum hydrophobicity of helix-1, average pLDDT of helix-2, etc.). A complete set of scripts for computing these features is available at https://github.com/Rocklin-Lab/mhdx_analysis/tree/main/scripts/feature_extraction. In total, we extracted 2,800 features common across folds and an additional 2,000–3,700 topology-specific features. In addition, we extracted embeddings from protein language models - ESM2 (650M parameters) (Lin et al. 2023), Unirep (Alley et al. 2019), and SaProt (Su et al. 2023) - to capture contextual sequence information. For ESM2, we averaged the representations from layers 33–36 and flattened the result into an array of 2,560 features per protein sequence, while Unirep and SaProt yielded 1,900 and 1,280 features, respectively.

### Feature correlation analysis

Pearson correlation coefficients (PCCs) were computed for all derived sequence and structural features with target variables, global stability (Δ*G*_unfold_), normalized cooperativity, and family-normalized cooperativity within each protein topology. PCCs were computed using the pearsonr function from SciPy Stats. Bootstrapping was performed with 1,000 resamples to generate the 95% confidence interval for PCCs. 10,000 random permutations of global stability (dg_mean), normalized cooperativity (normalized_cooperativity_model_global) and family normalized cooperativity (normalized_cooperativity_pf) were generated for the HHH and EEHEE topologies. PCCs were calculated for each random permutation of the target variables with the derived sequence and structural features. The mean and 95% confidence intervals were calculated from the distribution of PCCs of the random permutations with each feature and compared to the experimentally calculated PCC.

### Training of machine learning models

We implemented a machine learning pipeline to predict key protein properties - Δ*G*_unfold_ and family-normalized cooperativity - using regularized linear models (LassoCV and RidgeCV). We independently evaluated features derived from protein language model embeddings (ESM-2, SaProt, Unirep) alongside interpretable features. Our pipeline systematically tests 20 model variations per feature set, comparing models that use the original features to those employing expanded features (including square, inverse, and logarithmic transformations), and optionally applying Pearson correlation filtering at thresholds of 5%, 10%, 25%, and 50% to remove weakly correlated features. Model training was performed using 5-fold cross-validation across three independent random splits. To ensure reproducibility and biological relevance, we applied dynamic clustering with MMseqs2 (Hauser, Steinegger, and Söding 2016) to group similar sequences. For each protein family, sequences were iteratively clustered using a range of minimum sequence identity thresholds (from 0.1 to 0.75), with clustering halted when the largest cluster dropped below 10% of the total sequence count - resulting in final thresholds between 0.45 and 0.70 across families. These sequence clusters were then randomly assigned to the 5 folds, ensuring that each fold was as distinct as possible while maintaining balanced representation across protein families (PFs).

In contrast, a first-generation set of models was trained on an earlier version of the dataset where protein language model embeddings were derived from ProteinMPNN (Dauparas et al. 2022), ESM Inverse Folding 1 (ESM_IF1) (Hsu et al. 2022), Tranception (Notin et al. 2022), ESM2 (with both 650M and 3B parameters) (Lin et al. 2023), and hybrid models combining ESM2 with ESM_IF1. For these first-generation models, sequences were assigned randomly to 5-folds without applying identity-based clustering. Despite this difference in preprocessing, the overall training strategy - employing 5-fold cross-validation across three independent random splits - remained consistent across both approaches.

In both cases, global models were trained on the full dataset and evaluated on individual PFs, while family-specific models were developed for PFs with more than 200 examples. Model performance was assessed using the R² score aggregated across held-out sets, and final model selection was based on the highest mean R² across the three splits to maximize generalization.

## Code availability

The code for the analyses can be found at https://github.com/Rocklin-Lab/mhdx_pipeline, https://github.com/Rocklin-Lab/hdxrate_pipeline, and https://github.com/Rocklin-Lab/mhdx_analysis.

## Data availability

All data supporting the findings of this study, including those presented in the main text, extended data figures, and tables, are available for download at https://forms.gle/RwJwvfw6WN4gjXaD9. The mass spectrometry proteomics data have been deposited to the ProteomeXchange Consortium via the PRIDE (Perez-Riverol et al. 2025) partner repository with the dataset identifier PXD061702.

## Acknowledgements

We thank Susan Marqusee, Tobin Sosnick, Kresten Lindorff-Larsen, Lauren Porter, Tonya Zeczycki, Sarah Fahlberg, Darcy Kim, Vani Lorish, and members of the Rocklin Lab for valuable discussions and feedback throughout this work. We are grateful to the Northwestern Proteomics Core for assistance with mass spectrometry experiments and to members of Waters Corporation for technical support. We also thank Yuriy Pyatkivskyy for insightful conversations and advice. This work was supported by the National Institutes of Health (NIH) Director’s New Innovator Award DP2GM140927 (G.J.R.), the São Paulo Research Foundation (FAPESP) Grant #20/14421-1 (A.J.R.F.), NIH T32GM008382 (M. Garcia), NIH T32GM149439 (C.M.M.), the PhRMA Foundation Predoctoral Fellowship in Drug Delivery (C.M.M.), National Science Foundation (NSF) NRT Award 2021900 (C.M.P.), NIH F31GM151811 (C.M.P.), Human Frontier Science Program Long-Term Fellowship (K.T.), JST PRESTO Grant JPMJPR21E9 (K.T.), and NSF Award 2304707 (M. Guttman). This research was supported in part through the computational resources and staff contributions provided for the Quest high performance computing facility at Northwestern University which is jointly supported by the Office of the Provost, the Office for Research, and Northwestern University Information Technology.

## Contributions

G.J.R. conceived the project. C.M.P., J.T., and M.J. cloned the DNA constructs, while J.T., A.J.R.F., C.M.P., M.Garcia, S.M.D., L.C., and C.M.M. expressed the protein libraries and individual proteins. A.J.R.F. performed the mHDX-MS experiments, with optimization of the protocols carried out by A.J.R.F., J.T., and S.M.D.. A.J.R.F., S.M.D., R.W.L., and G.J.R. developed the computational pipeline for mHDX-MS analysis. A.J.R.F., S.M.D., M.Garcia, J.T. and G.J.R. analyzed the mHDX-MS data. A.J.R.F., P.N., and C.M.M. further developed the machine learning models for predicting global stability and cooperativity. S.H. collected and analyzed the HDX NMR data under the supervision of C.H.A.. K.T. performed the cDNA display proteolysis experiments. M.Guttman provided guidance on HDX-MS experimental design and assisted with preliminary experiments. A.J.R.F. and G.J.R. wrote the manuscript with input from all authors. G.J.R. acquired funding for the project and supervised the research.

## Extended data

**Figure S1.**
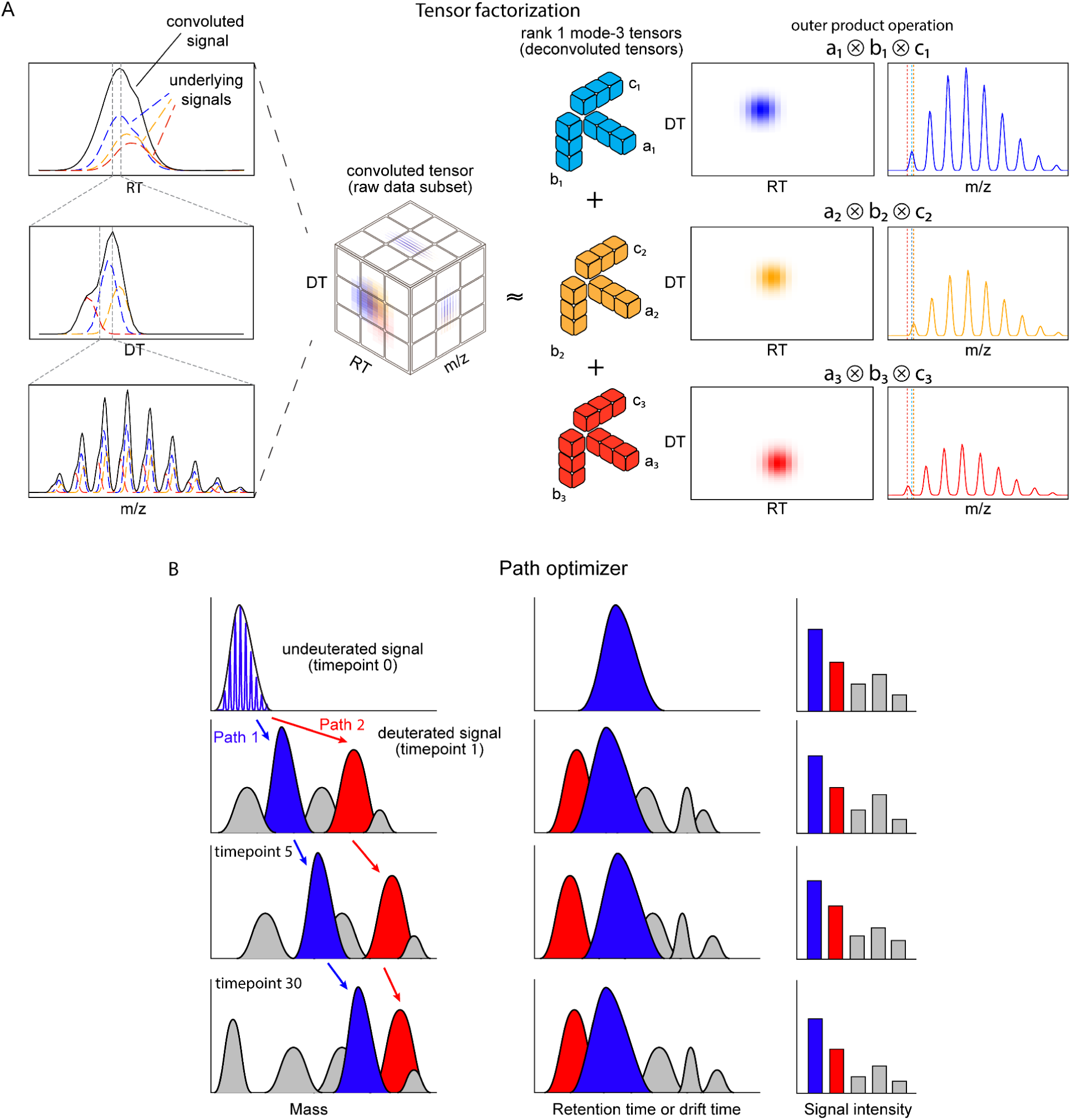
Tensor factorization and path optimization for mHDX-MS data. (A) Tensor factorization schematic: three underlying signals (blue, orange, red dashed lines) overlap in a single LC–IMS–MS experiment (black line). We represent the data as a 3D tensor spanning retention time (RT), drift time (DT), and m/z dimensions. By applying an iterative rank-decomposition approach (analogous to multi-dimensional nonnegative matrix factorization), each signal is approximated by a low-rank (rank-1) factor in each of the three modes. This effectively disentangles the individual signals, even when they overlap in multiple dimensions. (B) Path optimizer schematic: after deconvolving signals at each timepoint, the pipeline selects the most coherent set of isotopic clusters (ICs) to form a time-resolved mass profile. In this example, a “blue path” follows ICs that smoothly transition in mass, RT, and DT, whereas a “red path” includes clusters with abrupt transitions and poorer agreement, leading to a higher penalty. The optimizer computes a multidimensional score (penalizing implausible changes in mass uptake, large RT/DT errors, etc.) and iteratively refines the path until no single substitution of an IC can further improve the overall score. The final “winning” path provides the most physically consistent deuteration trajectory from the undeuterated to the fully deuterated state.

**Figure S2:**
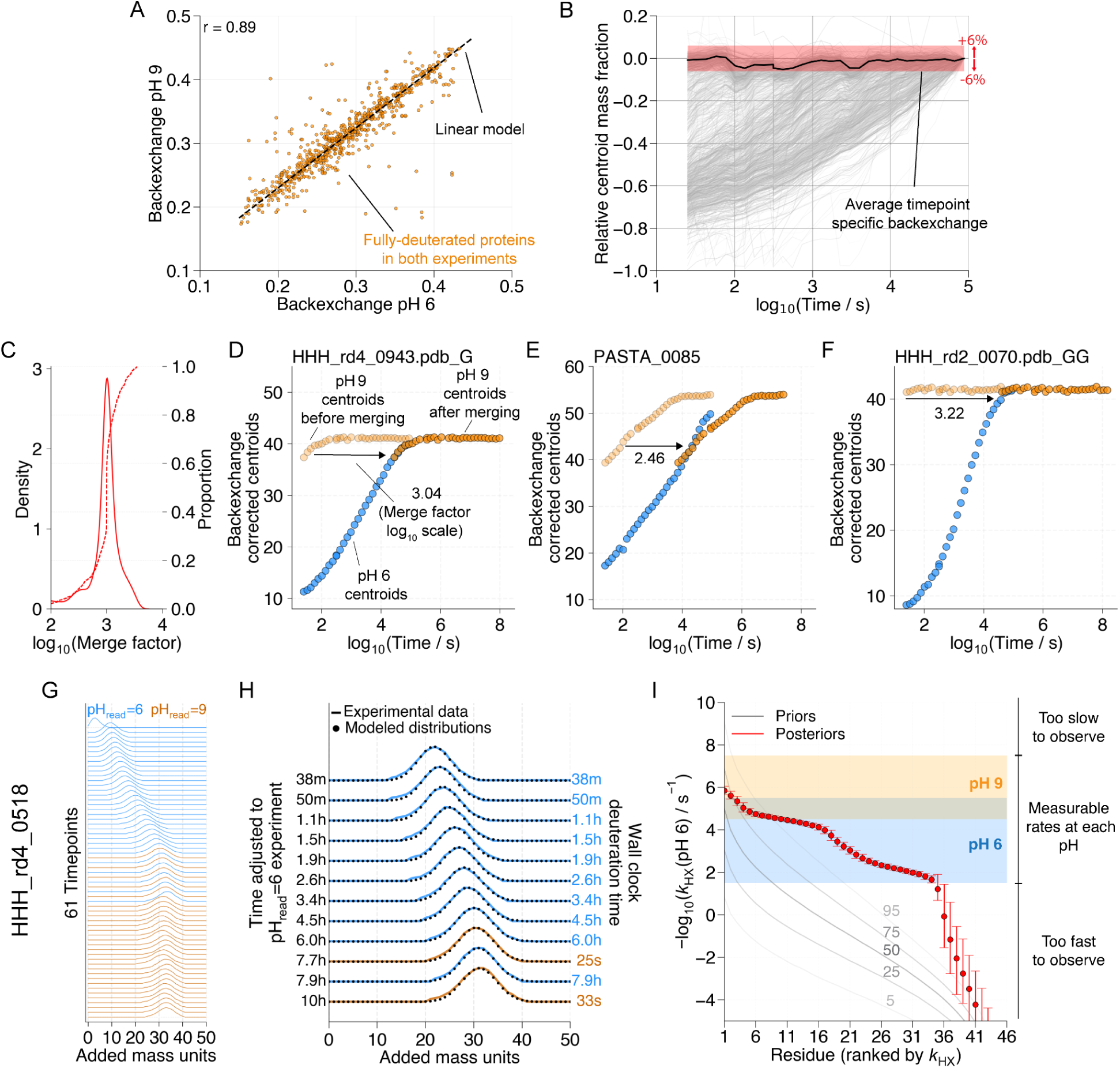
Inferring exchange rates (k_HX_) from mHDX-MS data. (A) Timepoint-specific back exchange correction: injection-to-injection variability leads to a different overall level of back exchange at each timepoint for all proteins. We identify the central trend (black line) using the subset of proteins that reach full deuteration by the fifth-to-last timepoint, with the red shaded area indicating the mass tolerance used to include centroids in computing the trend. This timepoint-specific back exchange is added to the protein-specific back exchange when we infer exchange rates. (B) Protein-specific back exchange normalization: we determine the average back exchange for all proteins based on the total deuteration in the last five timepoints, if the deuteration level is no longer increasing. For proteins that do not reach full deuteration at pH 6, we infer their pH 6 level of back exchange based on the back exchange observed at pH 9. We use a linear model to infer pH 6 back exchange based on pH 9 backexchange, parameterized using proteins identified as fully deuterated under both conditions. (C) Distribution of merge factors across proteins for which both pH 6 and pH 9 data were combined to infer exchange rates; pH 9 is expected to accelerate observed rates by a factor of 10³ (Connelly et al. 1993; Nguyen et al. 2018; Bai et al. 1993). (D–F) Representative centroid trajectories highlighting three examples: (D) HHH_rd4_0943.pdb_G with a merge factor near 10³, (E) PASTA_0085 with a lower factor (2.43), and (F) HHH_rd2_0070.pdb_GG with a higher factor (3.22). (G) The merged isotopic distributions (pH 6 in blue, pH 9 in orange) for HHH_rd4_0518, illustrating how timepoints from both conditions form a continuous series when properly merged. (H) A closer view of the overlapping timepoint region, where the observed distributions (dots) are compared with theoretical distributions based on the EX2 model (Fig. 1E Eq. 1-4) and inferred rates. (I) Ranked exchange rates (*k*_HX_) obtained via Bayesian inference, with the credible interval (5–95%) shown as error bars. The shaded blue and orange areas indicate the dynamic range covered by pH 6 and pH 9 experiments, respectively. The Bayesian priors are shown as grey lines.

**Figure S3.**
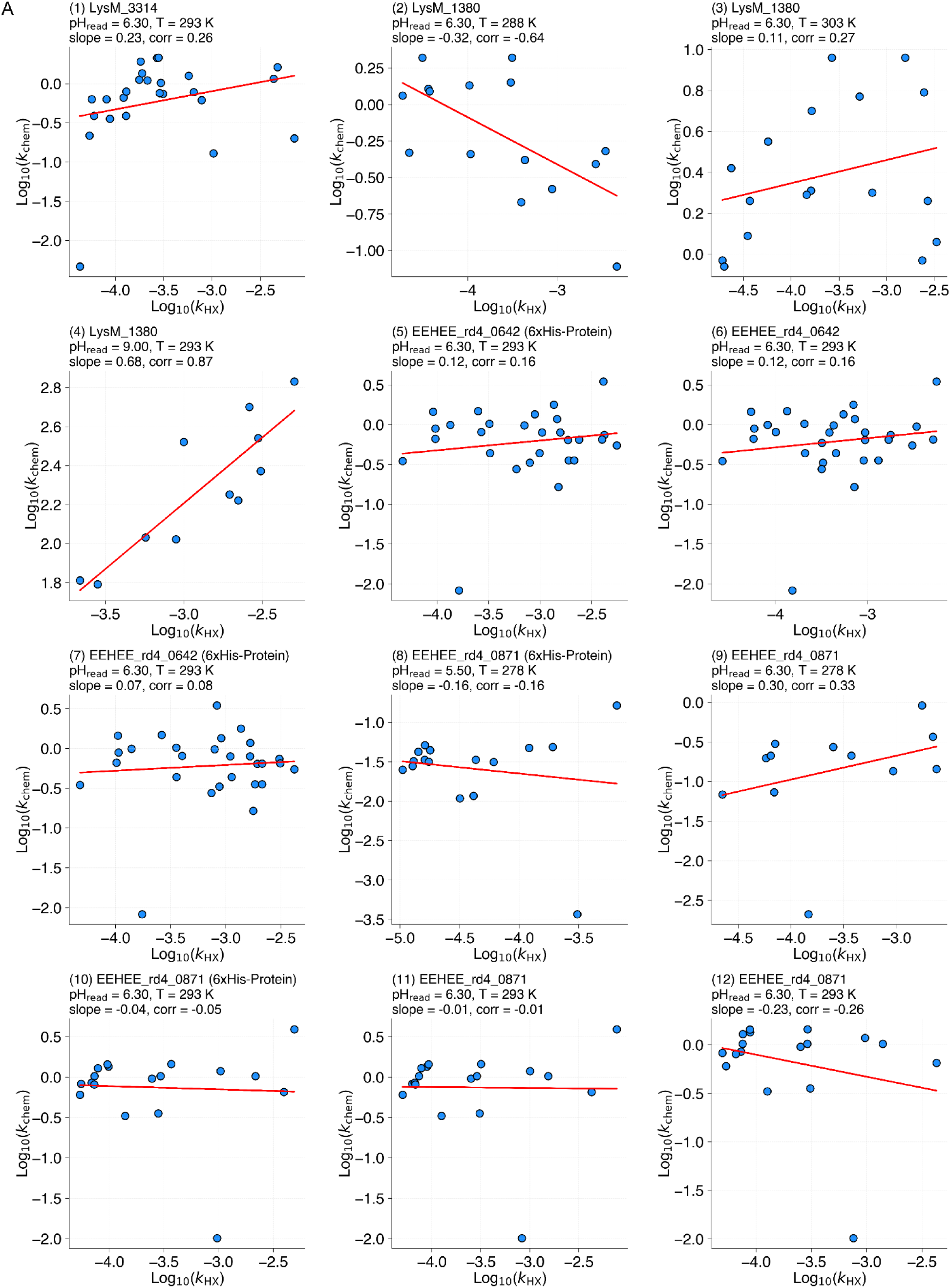
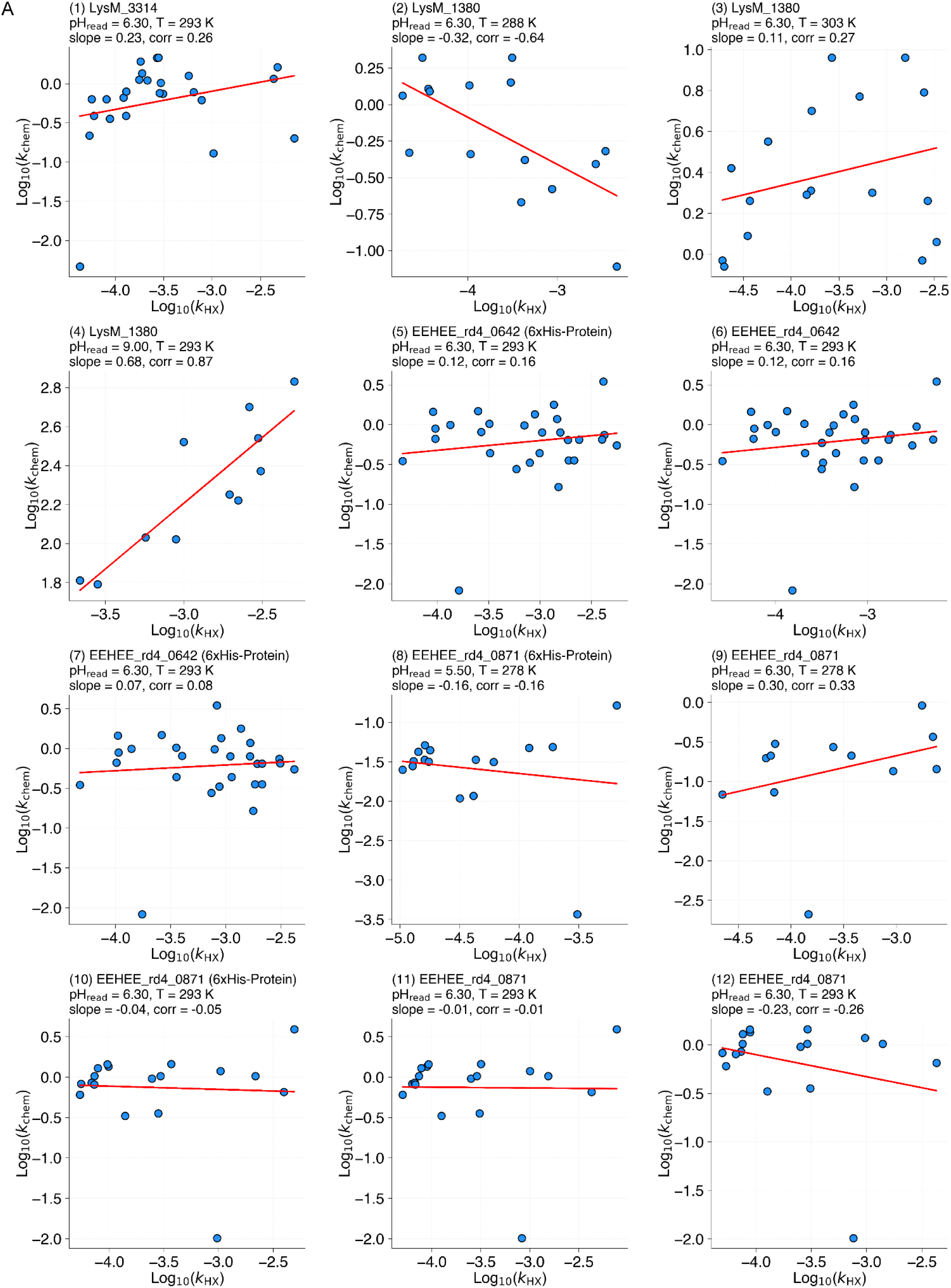
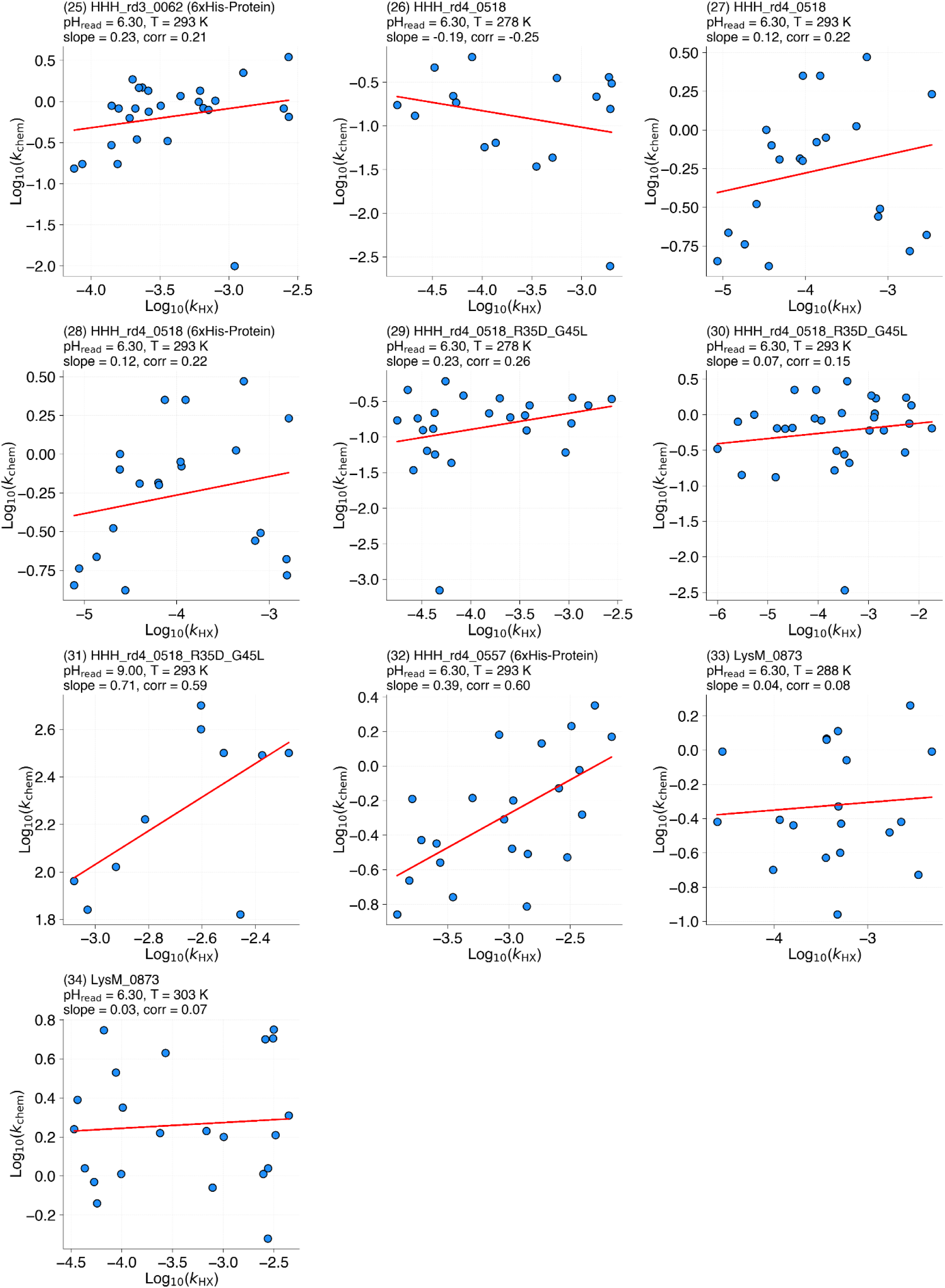
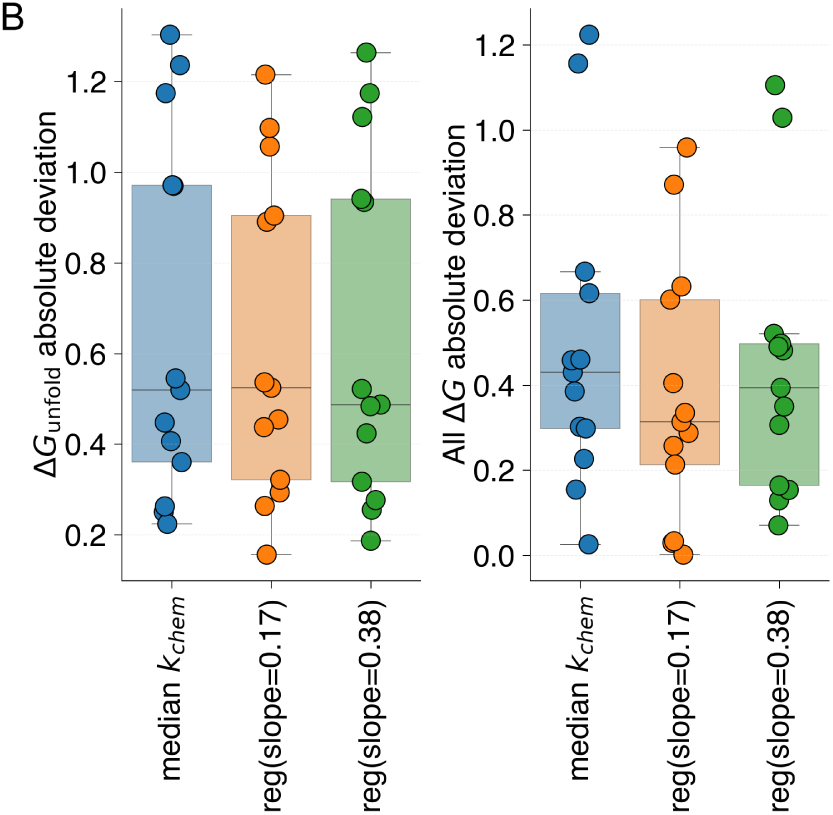
Estimation of intrinsic exchange rates and residue opening free energies (Δ*G*_open_). **(A)** Correlation plots comparing the logarithm of experimentally inferred exchange rates (log *k*_HX_) with intrinsic exchange rates (log *k*_chem_) derived from residue-resolved HDX NMR for 11 proteins. Each panel shows the relationship for one protein and one condition (temperature, pH), demonstrating that slower *k*_HX_ values tend to be associated with slower *k*_chem_. This trend is captured by fitting a linear model to z-scored log *k*_HX_ and log *k*_chem_ values, which forms the basis for our regression adjustment. (B) Boxplots comparing the absolute deviation in global stability (left) and across all measurable Δ*G*_open_ values (right) when estimating *k*_chem_ by (i) simply using the protein’s median *k*_chem_, (ii) using a regression model derived from a reduced dataset (slope = 0.17), and (iii) using a regression model based on the complete dataset (slope = 0.38). The regression-based adjustments yield Δ*G*_open_ estimates that more accurately recapitulate those obtained from HDX NMR, with the model using slope = 0.38 providing superior performance.

**Figure S4.**
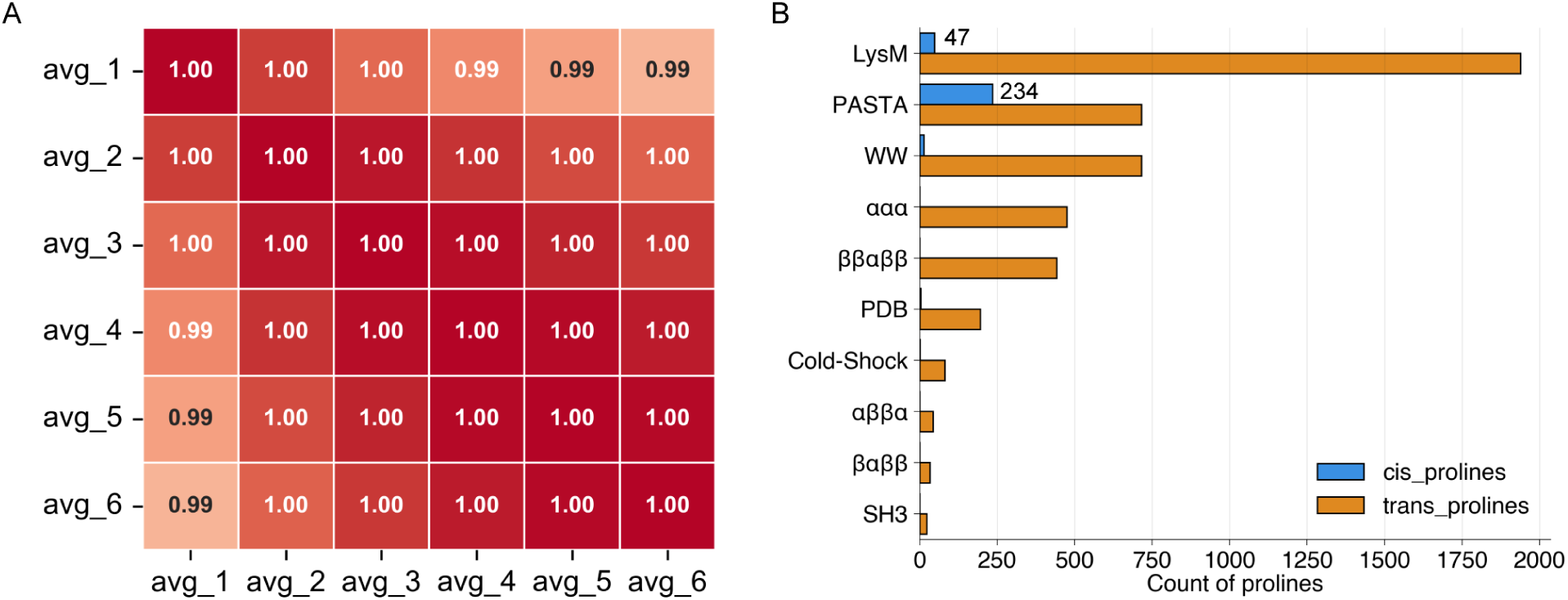
Robustness of Δ*G*_unfold_ estimation via averaging the most stable residues. (A) The correlation matrix displays pairwise Pearson correlation coefficients for stability values obtained by averaging the top 1 to 6 most stable residues (labeled avg_1 to avg_6). All pairwise correlations are ≥0.99, indicating that the stability metric is virtually invariant to the number of residues included in the average. (B) Counts of cis and trans prolines across protein families included in **Table S1 : Dataset_3**. While trans prolines are abundant across all families, cis prolines are predominantly found in PASTA domains, with few occurrences in LysM and WW domains. These data indicate that potential effects of cis prolines on Δ*G*_unfold_ estimations are largely confined to a subset of PASTA sequences.

**Figure S5.**
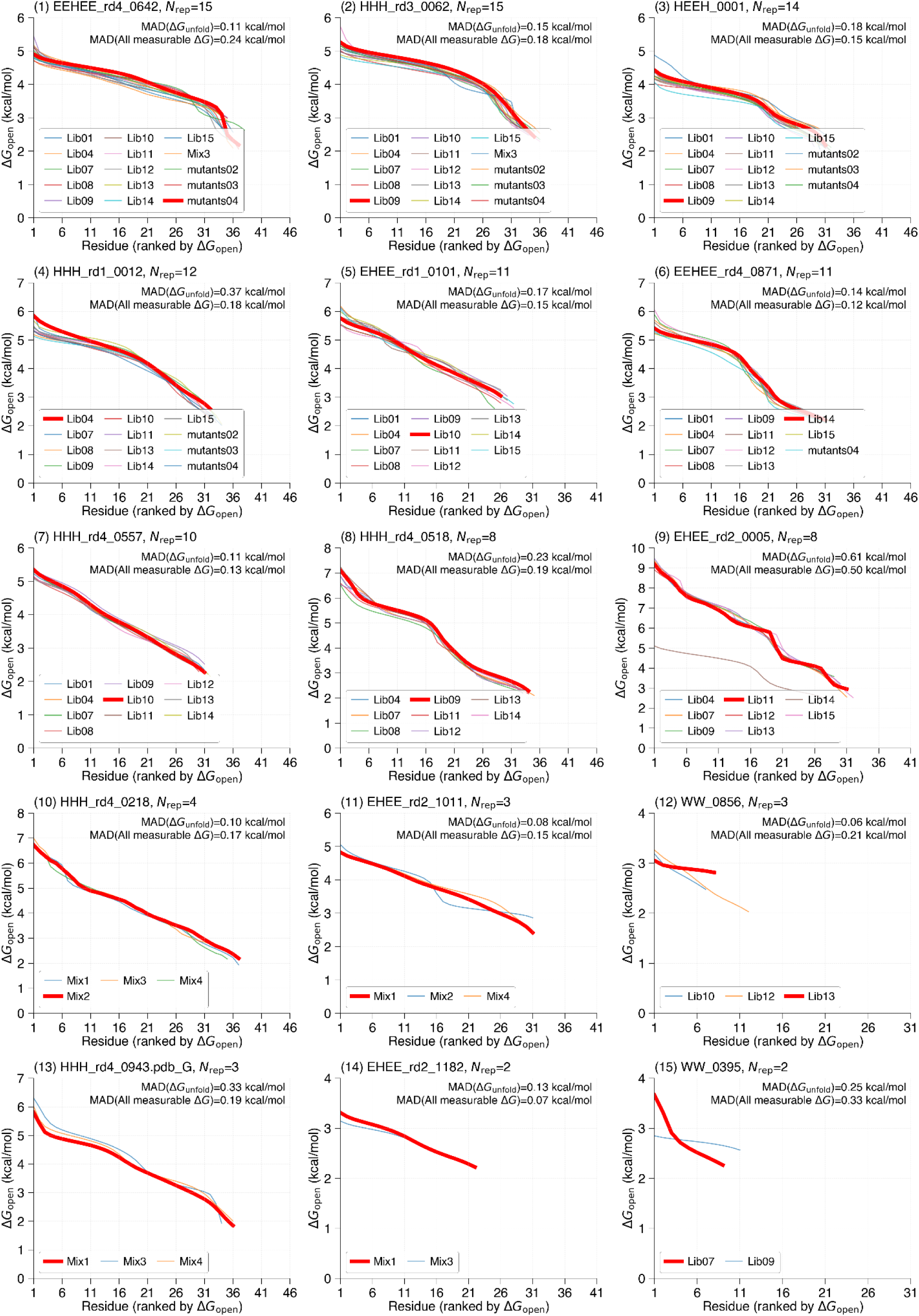
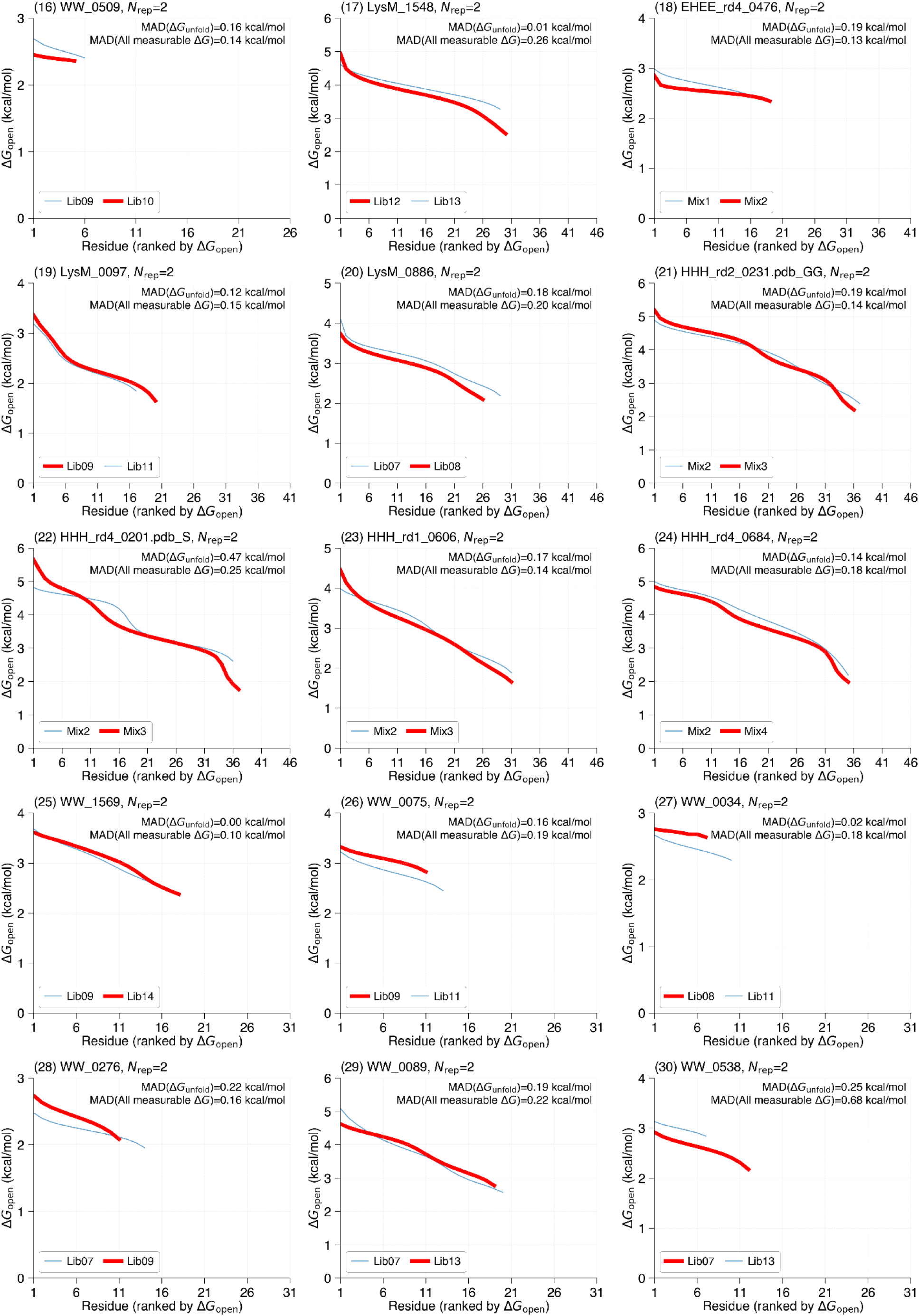
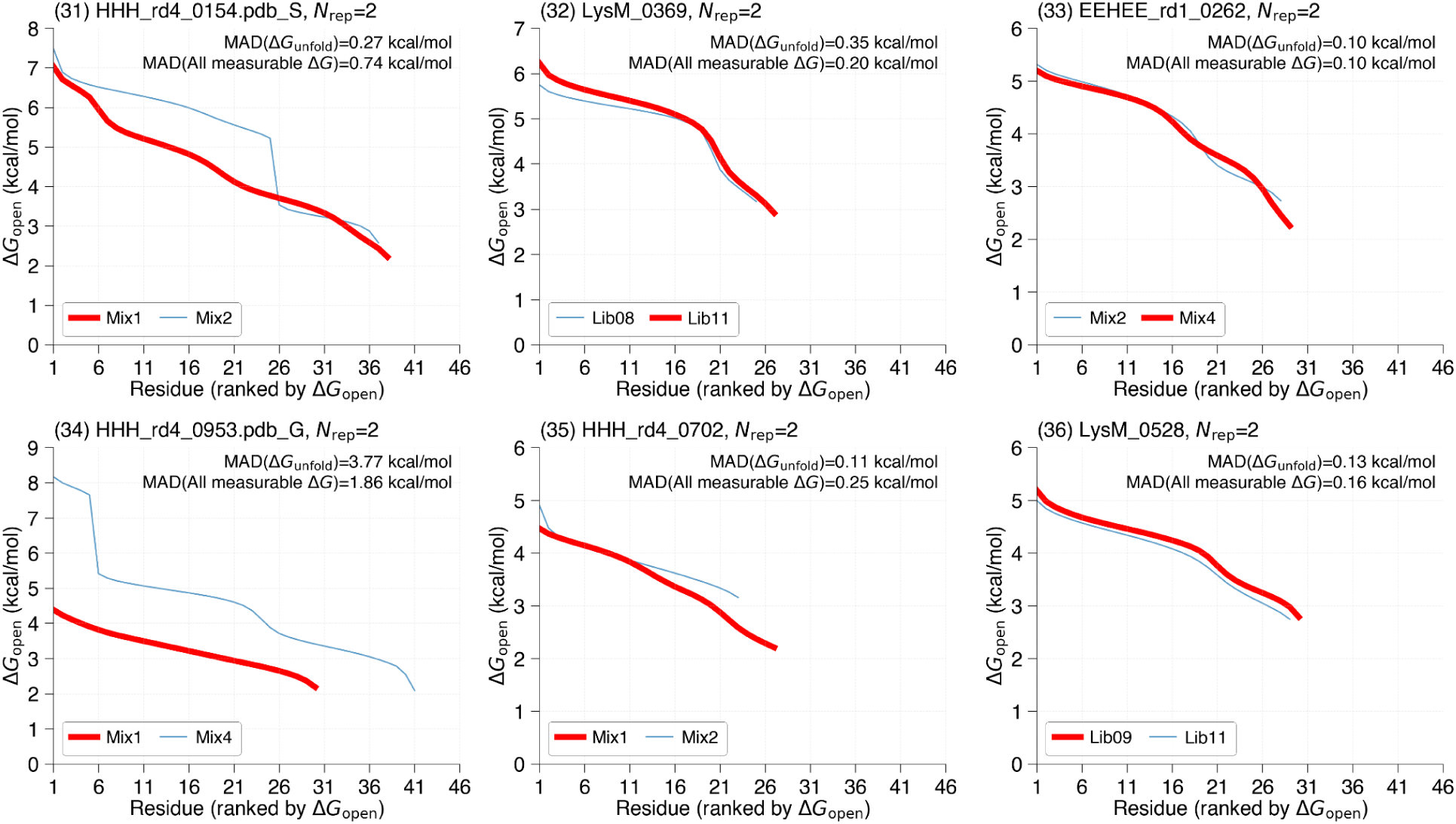
Reproducibility of mHDX-MS measurements across independent libraries. Each panel shows the reproducibility for one protein sequence that was measured in at least two independent mHDX-MS libraries (36 sequences in total). Panels are arranged in order of decreasing number of libraries in which the protein was detected. Within each panel, replicate measurements are displayed and the representative measurement - defined as the replicate with the lowest path optimizer score - is highlighted in red. The number of replicates (*N*_rep_) is indicated, and the mean absolute deviations for Δ*G*_unfold_ and for all measurable Δ*G*_open_ values are computed relative to the representative value.

**Figure S6.**
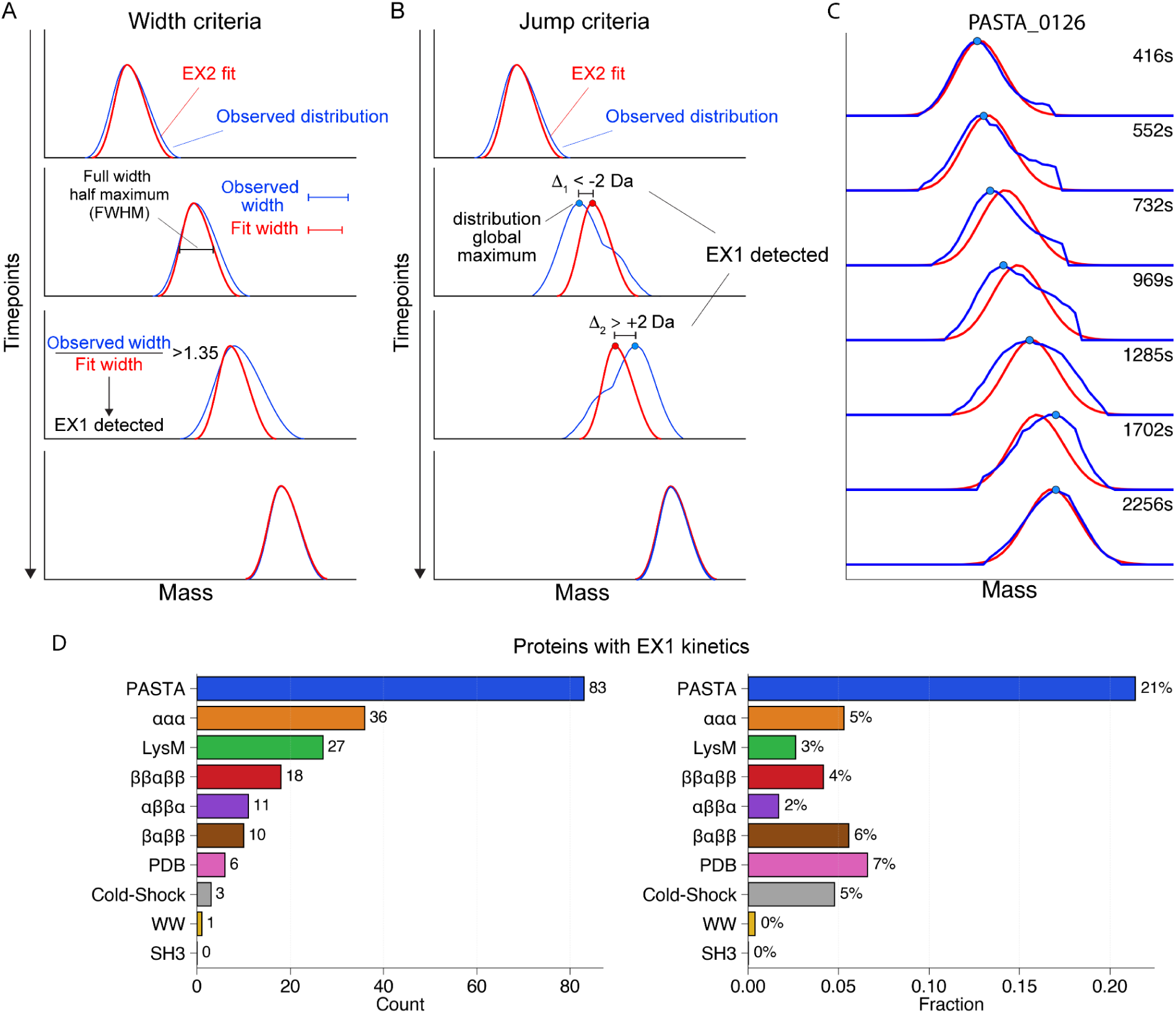
Detection and filtering of EX1 kinetics. (A) Illustration of the width criterion: the observed isotopic distribution widths (black) are compared with fitted reference distributions assuming EX2 kinetics (red), and the width ratio is computed during the exchange process (80–90% exchange). Samples with an average ratio >1.25 and a maximum ratio >1.35 are flagged as potential EX1 by the width criterion. (B) Illustration of the jump criterion: the difference between the centroid of the fitted EX2 distribution and the maximum peak of the experimental distribution is tracked over time, with abrupt shifts exceeding a predefined range (from <–2 Da to >+2 Da) before 90% exchange indicating EX1 behavior. (C) Example of PASTA_0126, whose exchange profile exhibits a pronounced “jump” consistent with EX1 kinetics as defined by the jump criterion. (D) Summary of EX1 cases identified in our dataset, shown as (left) counts and (right) percentages.

**Figure S7.**
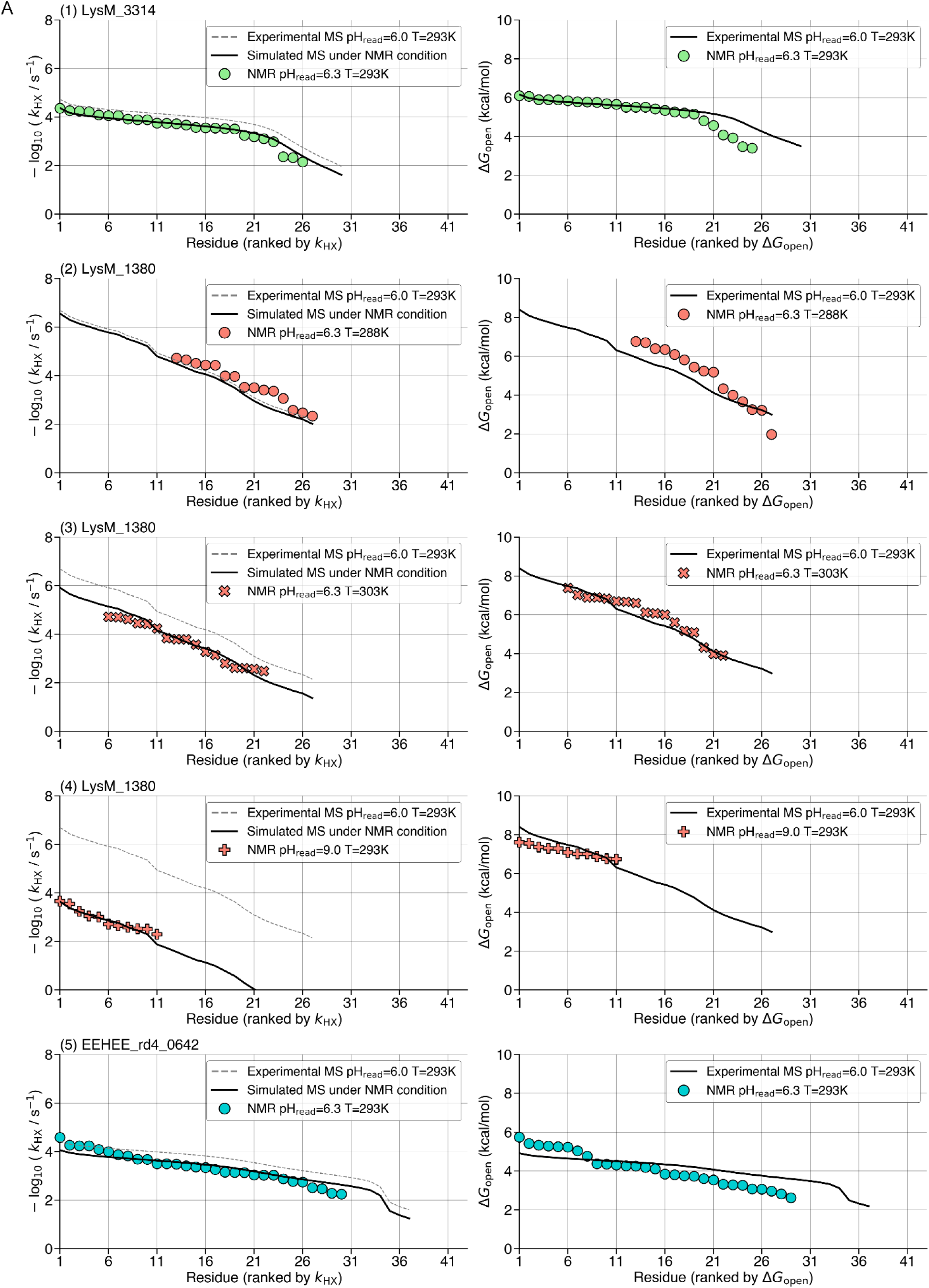
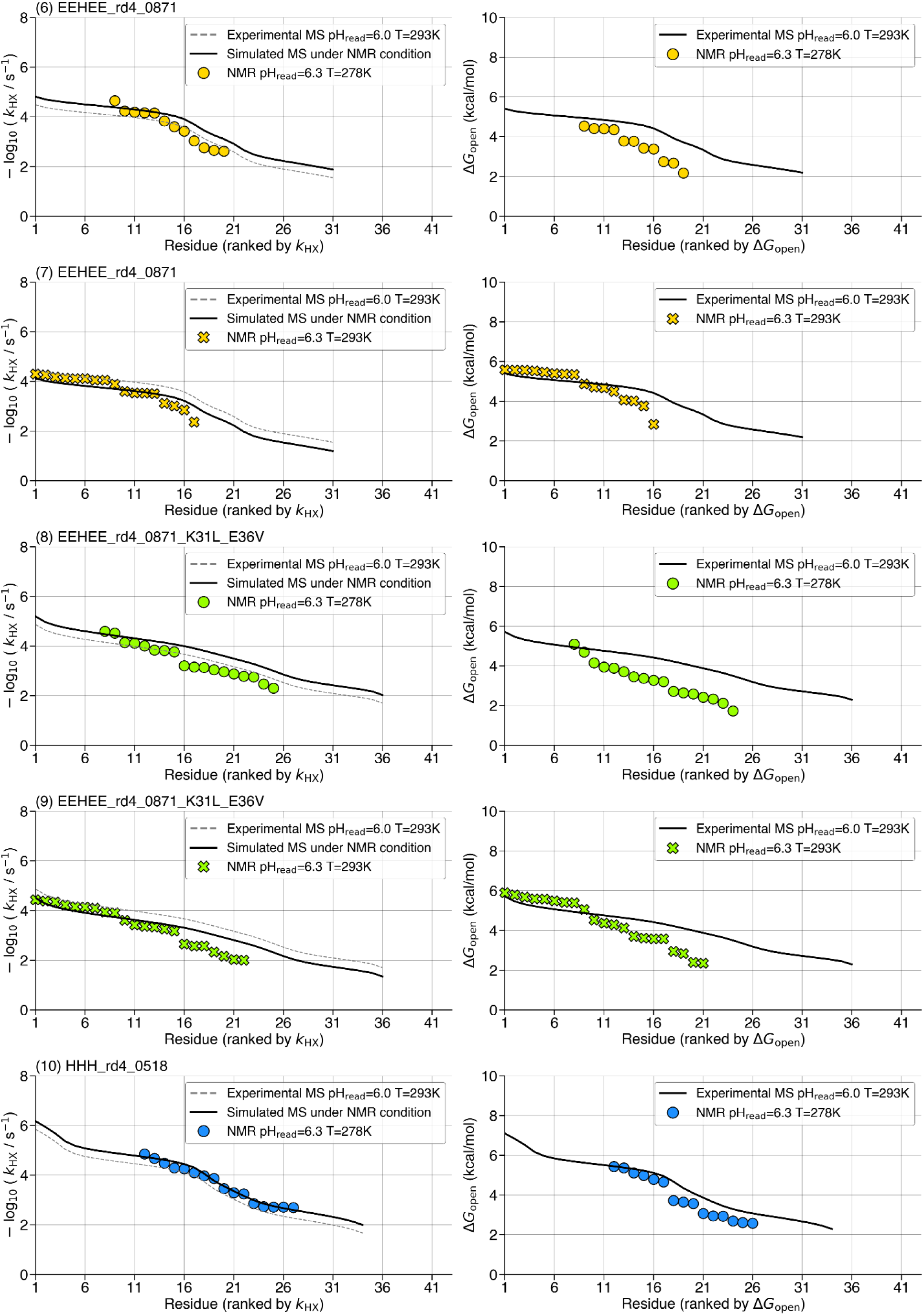
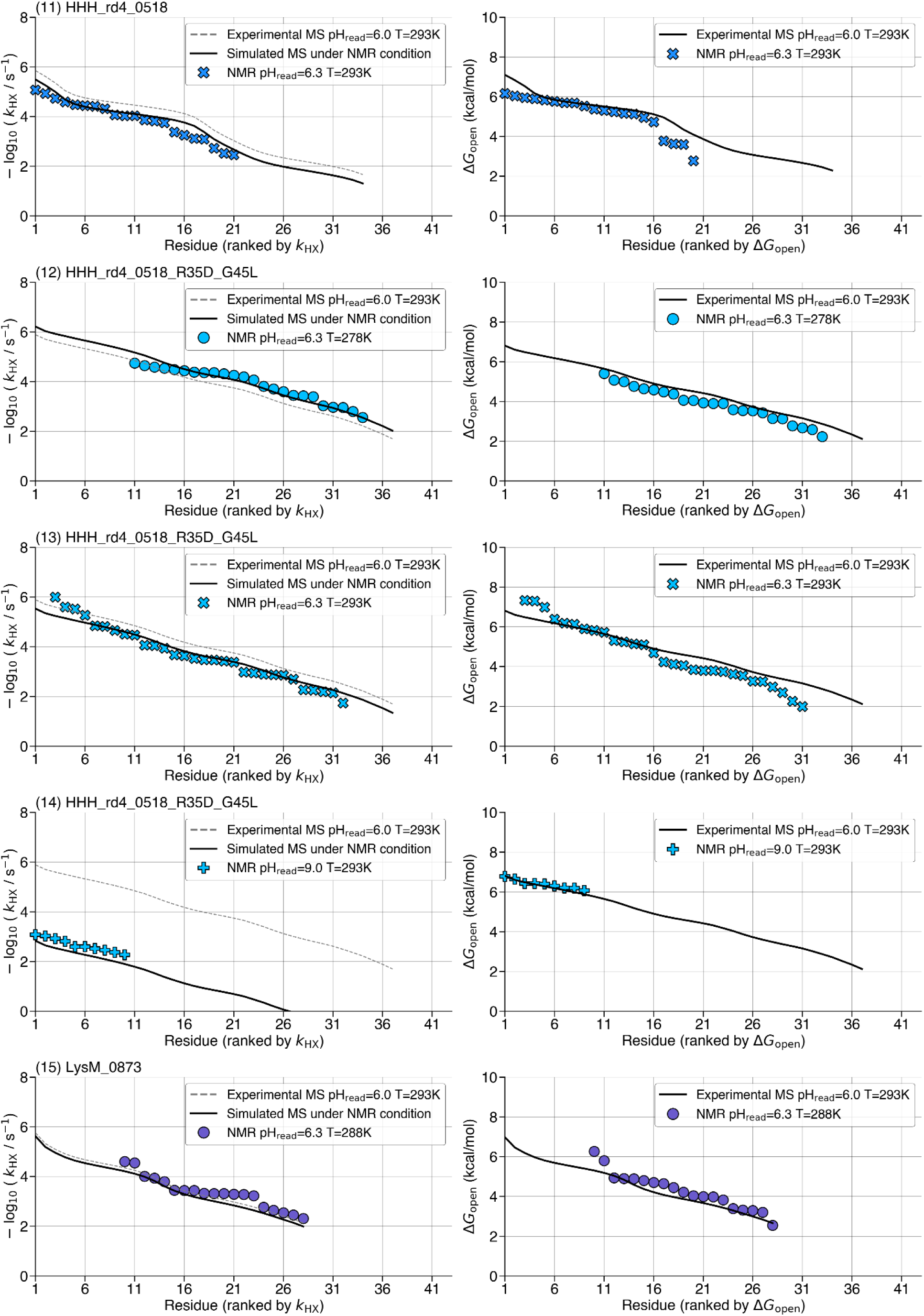
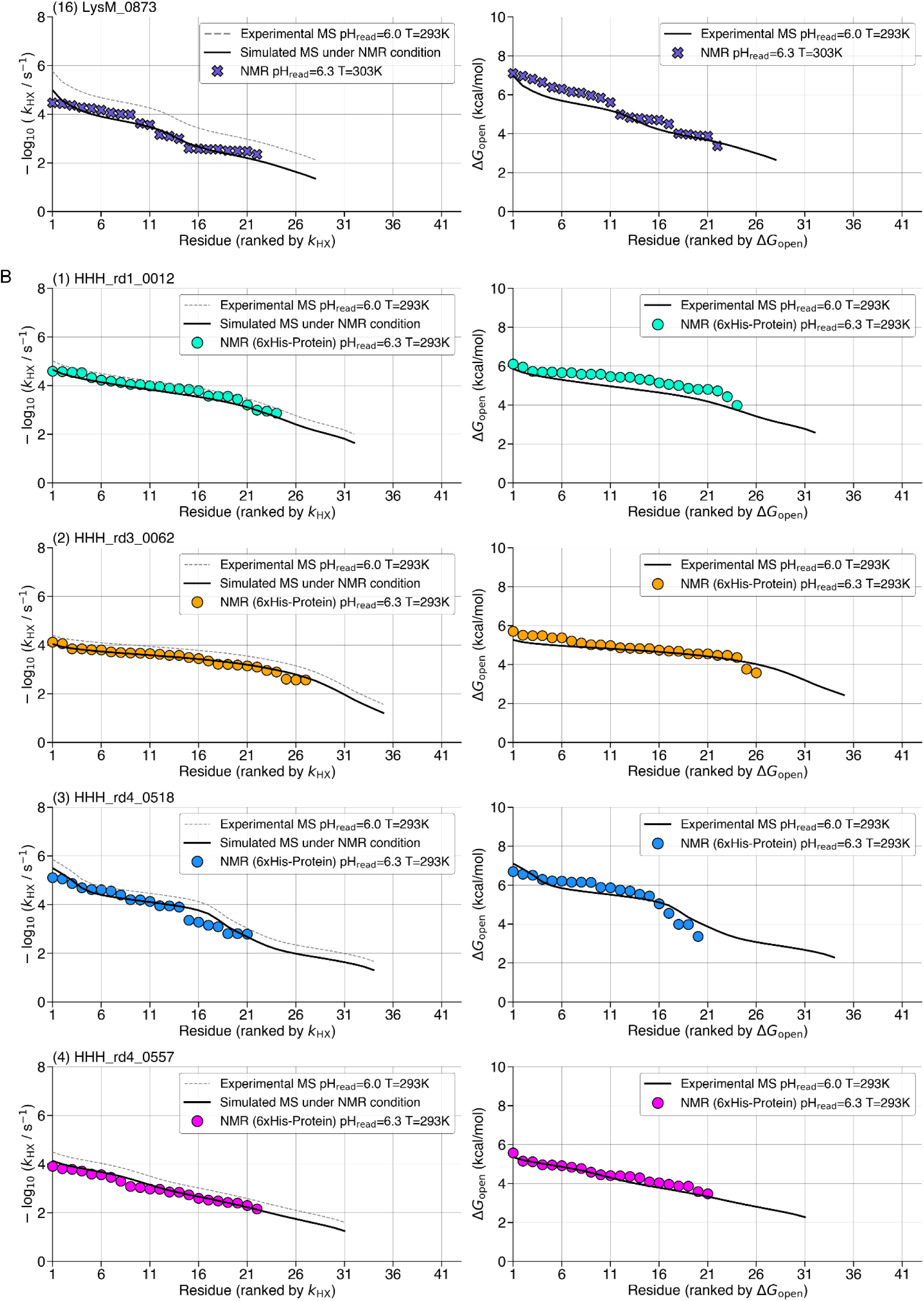
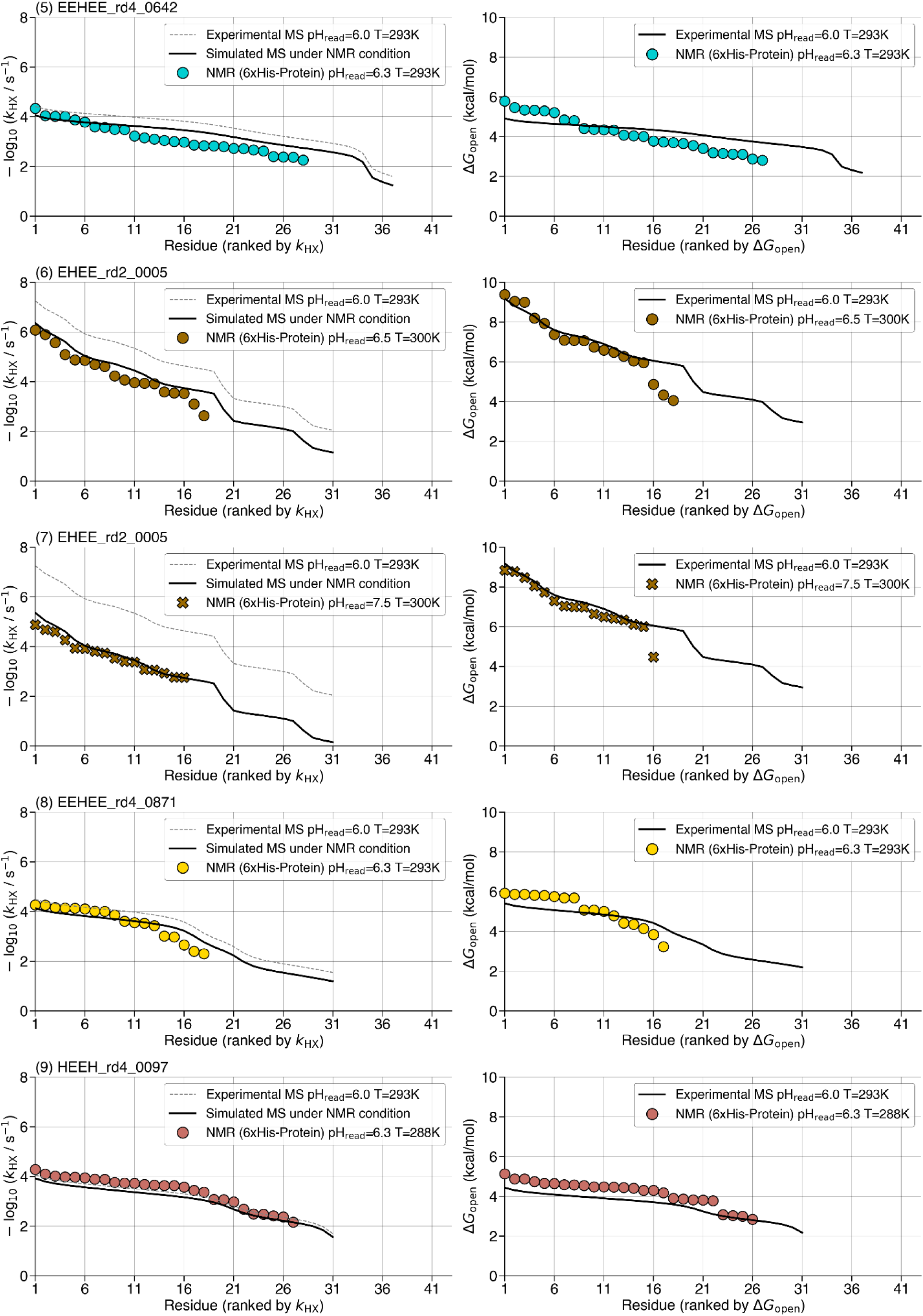
Comparative analysis of HDX NMR and mHDX-MS measurements. Thirteen unique protein domains were assessed using both multiplexed hydrogen-deuterium exchange mass spectrometry (mHDX-MS) and hydrogen-deuterium exchange NMR (HDX NMR) under various experimental conditions. All proteins analyzed by mHDX-MS had their 6×His-SUMO tags cleaved prior to measurement. Two classes of comparison are shown: (A) eight protein domains evaluated using nearly identical constructs between mHDX-MS and NMR, where the NMR constructs include only one or two extra residues at the N-termini; and (B) eight protein domain constructs where the NMR construct contains a long N-terminal tag (MGSSHHHHHHSSGLVPRGS). Each row corresponds to a specific protein under a given experimental condition, with multiple rows per protein representing different HDX NMR conditions. The left panels display exchange rates (*k*_HX_) from mHDX-MS (grey dashed lines) and HDX NMR (scatter), along with simulated mHDX-MS rates under replicated NMR conditions (black line), illustrating the agreement between the two methods. Simulated mHDX-MS rates were derived by modifying the measured rates according to the ratio between the median *k*_chem_ between the two conditions. The right panels show the corresponding opening energy distributions (Δ*G*_open_) from both mHDX-MS (black lines) and HDX NMR (scatter). Exchange rates were measured at pH 6.0-6.3 (MES buffer), pH 7.5 (PBS buffer), and pH 9.0 (bicine buffer). When some residues exchanged too slowly to resolve by NMR, these residues are not shown. For example, in #3 LysM_1380 at pH 6.3 303K, the slowest five residues were too slow to determine *k*_HX_ over 24 hours, so no NMR rate is shown for residues rank 1 to rank 5.

**Figure S8.**
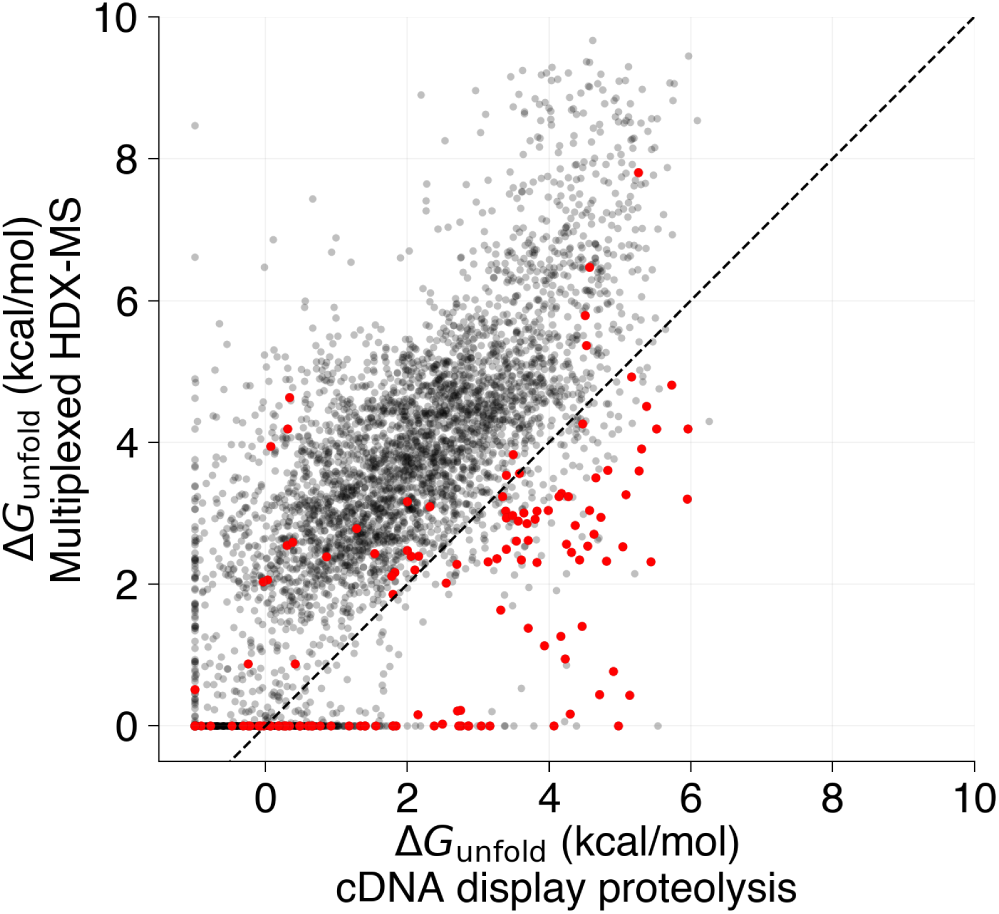
Correlation between Δ*G*_unfold_ measurements by mHDX-MS and cDNA display proteolysis. Scatterplots show the Δ*G*_unfold_ values obtained from both methods for 4,464 domains, with an overall Pearson correlation coefficient (r) of 0.75. Cold-Shock domains (highlighted in red), which are known to bind DNA, systematically exhibit greater stability in the cDNA display proteolysis assay. Excluding these domains (N = 4,327) improves the correlation to r = 0.78. ΔG_unfold_ values are clipped to 0 kcal/mol for mHDX-MS and to −1 kcal/mol for cDNA display proteolysis.

**Figure S9.**
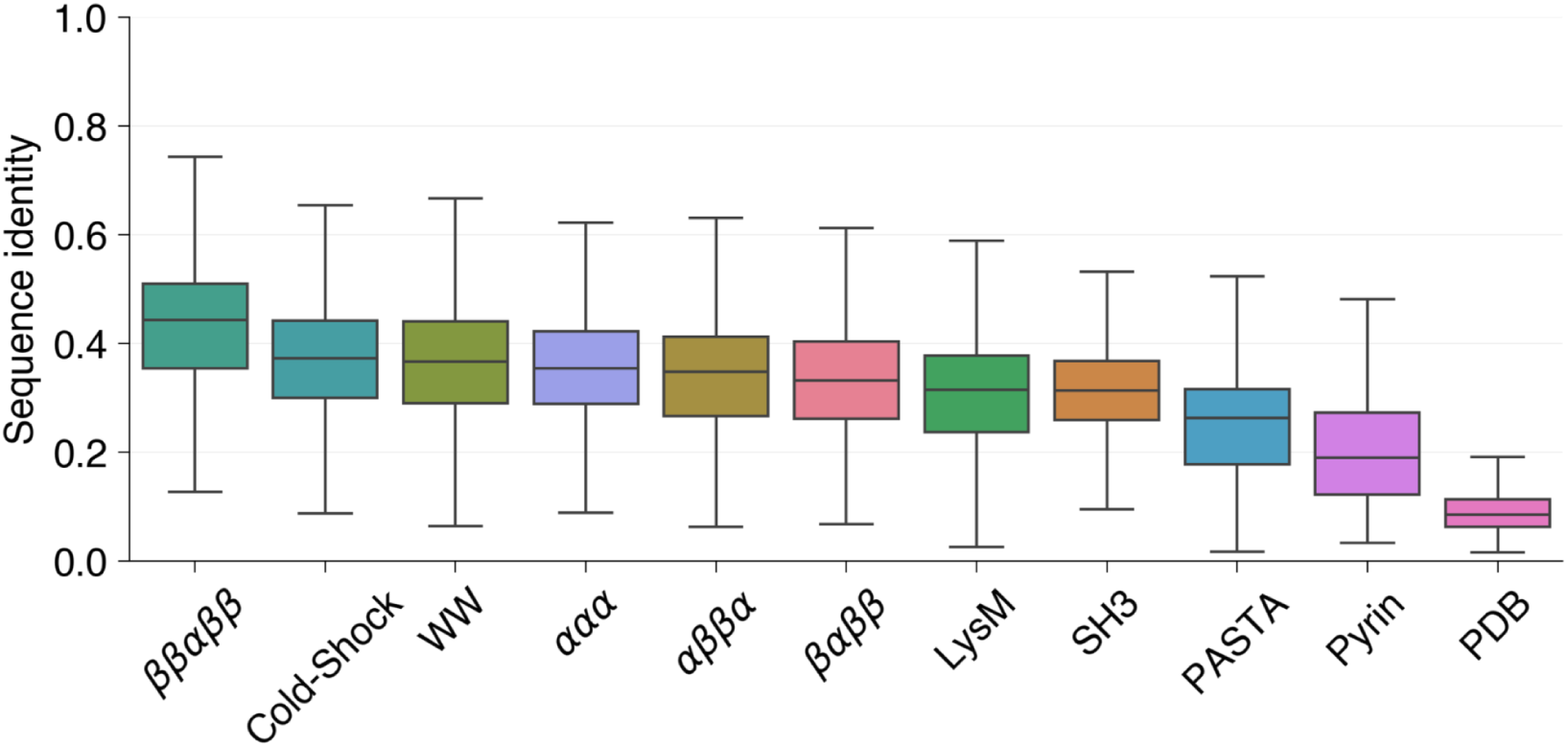
Distribution of pairwise sequence identity for different protein families. Sequences within each family were clustered using MMseqs2 (easy-cluster) with a dynamic threshold for minimum sequence identity (--min-seq-id ranging from 0.1 to 0.75), stopping when the largest cluster contained ≤10% of the total sequences. Representative sequences from the final clustering step were selected, and pairwise sequence identity was computed against all members within their respective clusters using MMseqs2’s search function. Alignments were filtered to remove redundant sequence pairs, retaining only the highest-scoring alignment per unique sequence pair. The identity values were further corrected by weighting sequence identity by alignment length relative to the shortest sequence in the pair. Alignments between identical sequences (query == target) were excluded, and spurious short alignments were removed. The final distribution aggregates pairwise identity values across all clusters within each protein family. Outliers were hidden in the boxplot for clarity.

**Figure S10.**
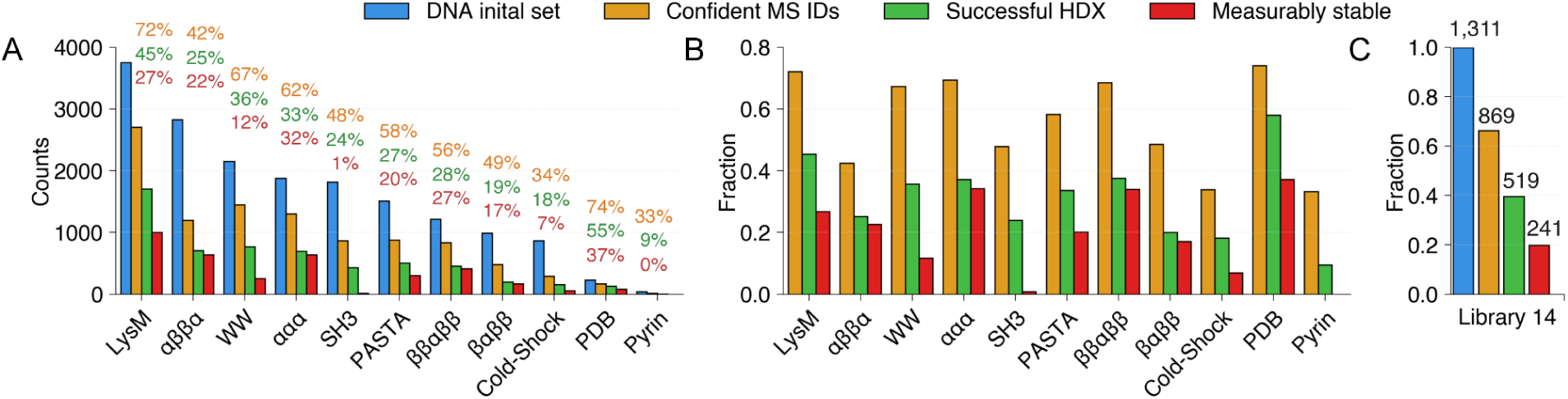
Success rate of mHDX-MS data processing across protein families. (A) Raw counts of protein families at successive stages of the analysis pipeline: the set of confident protein identifications (Confident IDs), proteins with successful HDX measurements (Successful HDX), and proteins classified as measurably stable. (B) The same three categories are expressed as percentages relative to the initial DNA set, illustrating the attrition at each processing stage. (C) A detailed breakdown for Library 14, displaying the counts for confident IDs, successful HDX, and measurably stable proteins obtained in one single experiment.

**Figure S11:**
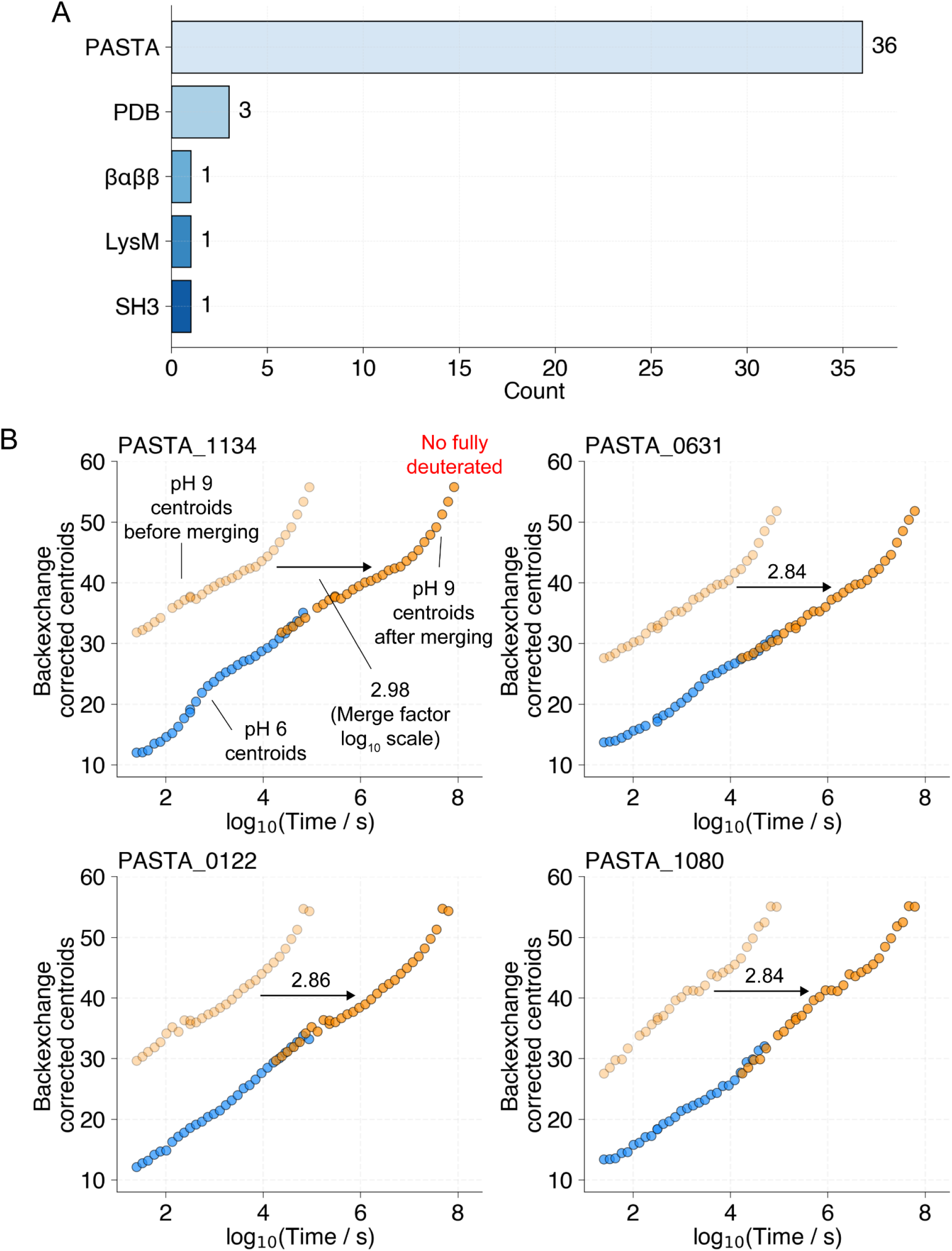
Incomplete deuteration in protein domains. (A) Number of proteins exhibiting a difference greater than 2 Da between the centroid masses measured at the last and fifth-to-last timepoints during the pH 9 experiment, indicating incomplete deuteration. (B) Four PASTA domains with the highest mass differences between the last and fifth-to-last timepoints, suggestive of very slow exchange rates. Blue (pH 6) and transparent orange (pH 9) points indicating the centroid of the mass envelope (after correcting for backexchange and the mole fraction of D_2_O) are shown at the true experimental timepoint. Solid orange points (pH 9) are shown at the inferred time point on a pH 6 scale after merging the pH 6 and pH 9 data together, with the merge factor indicating the adjustment made to the pH 9 timepoints (on a log_10_ scale). A merge factor of 3 (i.e. a 1,000-fold difference in *k*_HX_) is expected between pH 6 and pH 9 in the ideal case where pH only modulates *k*_chem_ according to (Connelly et al. 1993; Nguyen et al. 2018; Bai et al. 1993).

**Figure S12.**
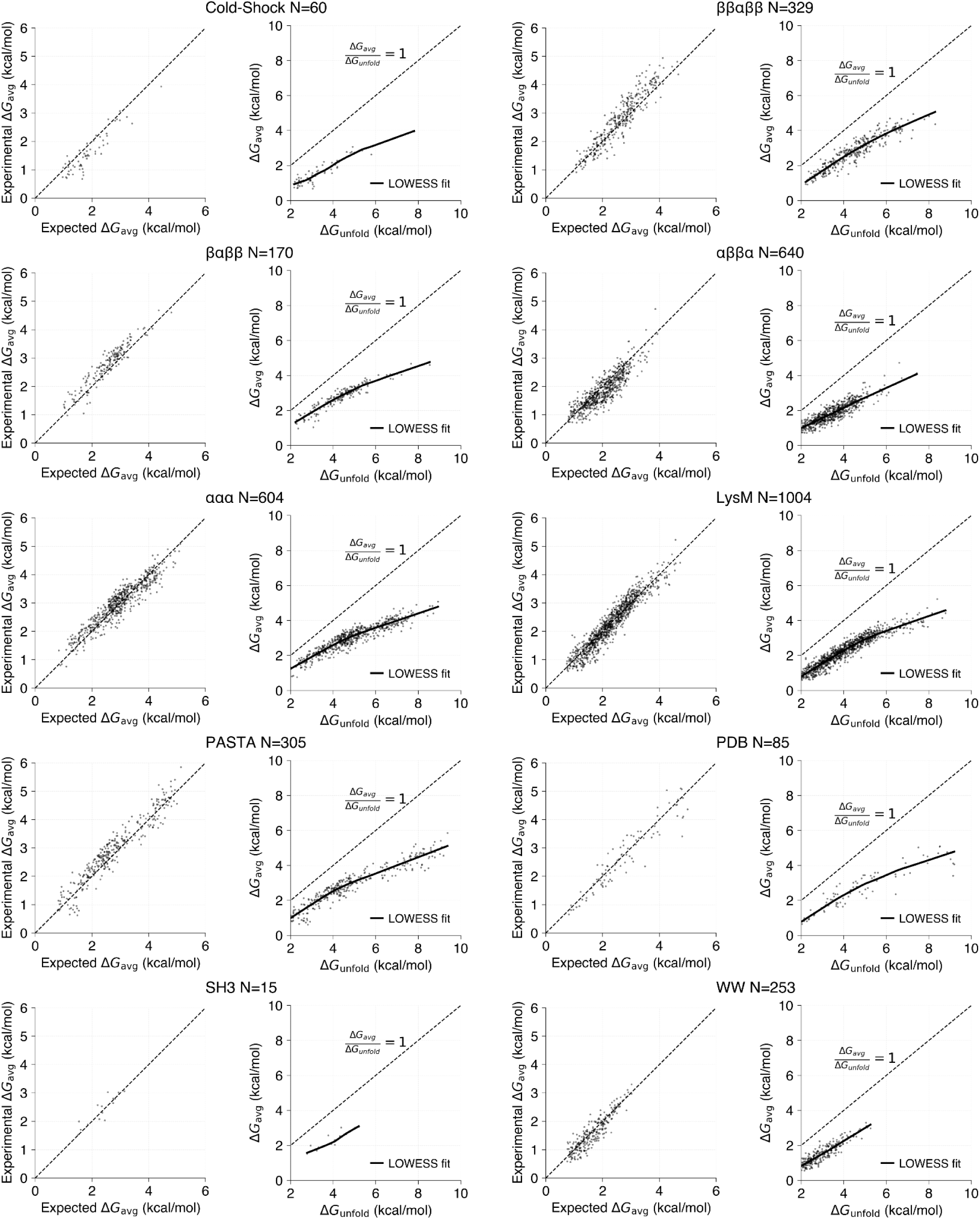
Empirical modeling of normalized cooperativity and family-normalized cooperativity. For each protein family, we fit an empirical model to predict the expected average opening free energy (Δ*G*_avg,expected_) as a function of global stability (Δ*G*_unfold_), the fraction of backbone hydrogen bonds (fxn_hb), and net charge. The model is defined as Δ*G*_avg,expected_ = *a* · (Δ*G*_unfold_ – *b*)*ᶜ* · (fxn_hb)*ᵈ* + *e* · netq, where the parameters *a*, *b*, *c*, *d*, and *e* are estimated from non-informative priors. Family-normalized cooperativity is then computed as the z-scored residual between the experimentally determined Δ*G*_avg_ and Δ*G*_avg,expected_, effectively decoupling cooperativity from stability. Additionally, for each protein family, we illustrate the sub-linear relationship between Δ*G*_unfold_ and Δ*G*_avg_, with a diagonal dashed line indicating a linear dependency for reference.

**Figure S13.**
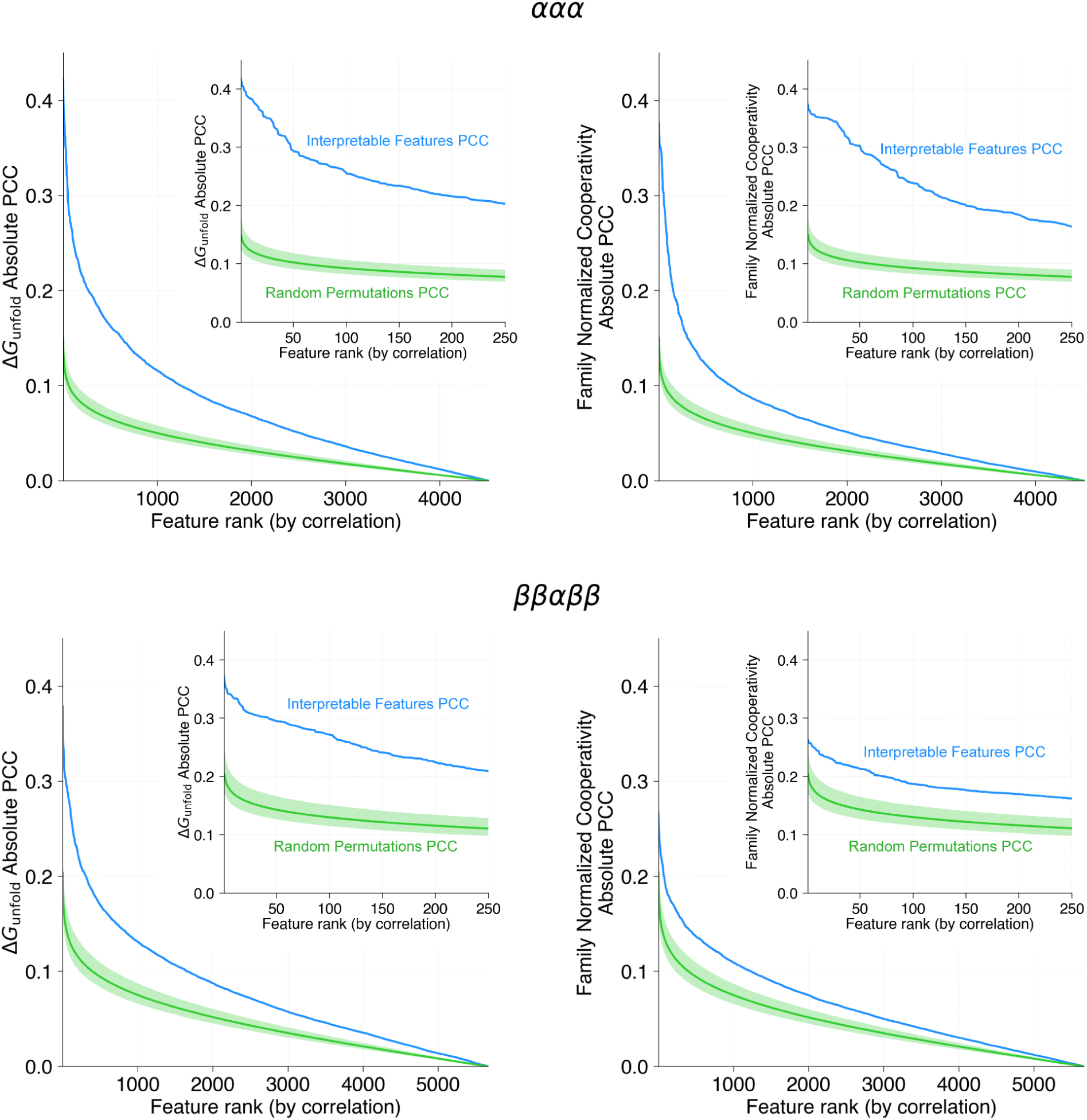
Permutation experiment for feature importance. In this analysis, 10,000 random permutations of the target variables—global stability (Δ*G*_unfold_) on the left and family-normalized cooperativity on the right—were generated for the ααα (top panels) and ββαββ (bottom panels) topologies. For each permutation, the Pearson correlation coefficients (PCCs) were computed between each manually engineered interpretable feature and the randomized target, and the features were then ranked according to these PCCs. This procedure essentially derives the order statistics of the random correlations, with the highest correlation value (first order statistic), the second highest, and so on, recorded to form a null distribution for each rank. Blue curves indicate the PCCs computed from the actual interpretable features, while green curves show the mean and 95% confidence intervals (shaded) of the corresponding null distributions. These comparisons demonstrate that the manually engineered features yield significantly higher correlations than expected by chance.

**Figure S14.**
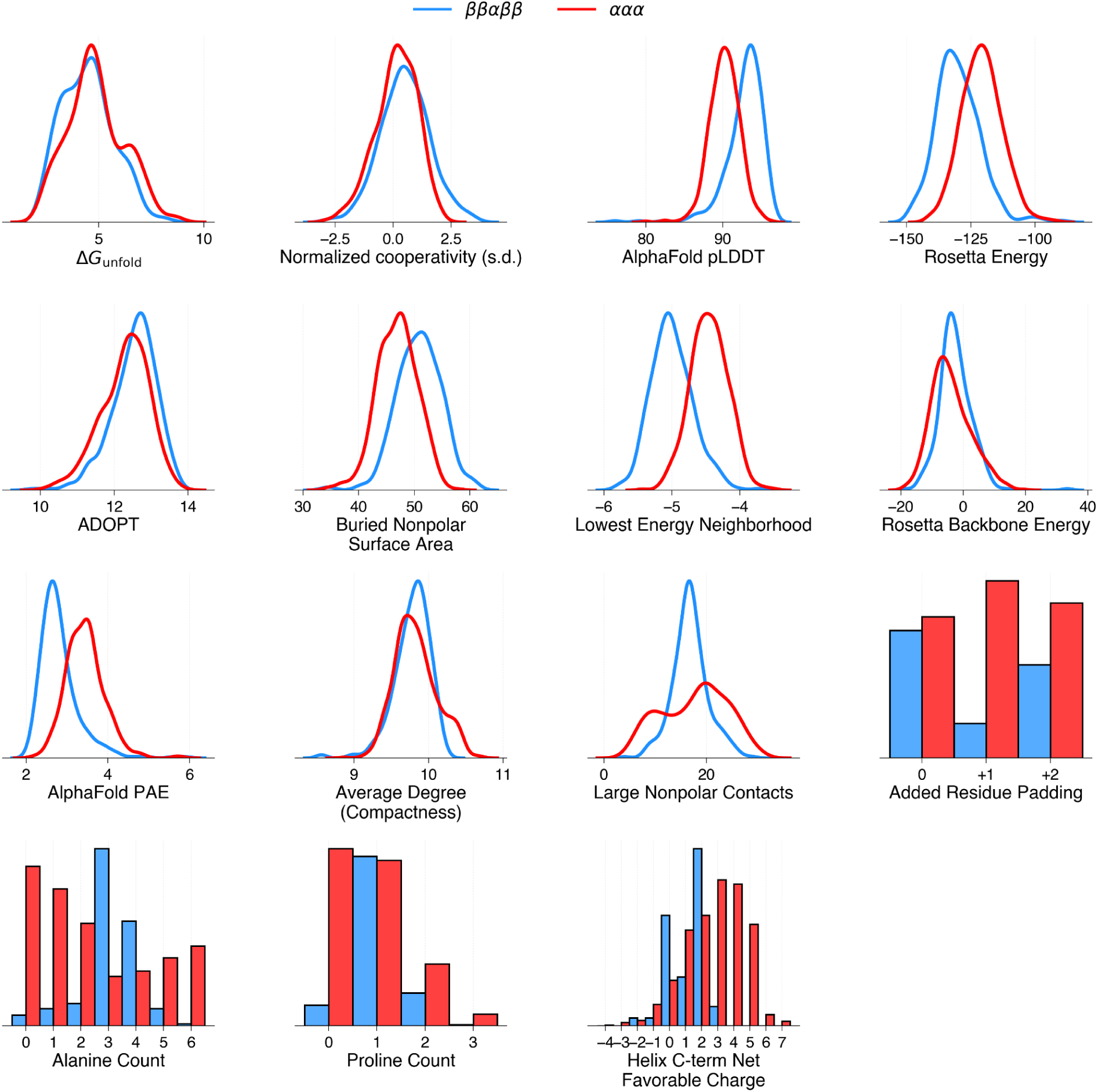
Feature distributions. Distributions for target variables and notable features highlighted in Figure 4 are shown for ααα (red) and ββαββ (blue) topologies. Kernel density plots are shown for continuous variables and histograms are shown for discrete variables.

**Figure S15:**
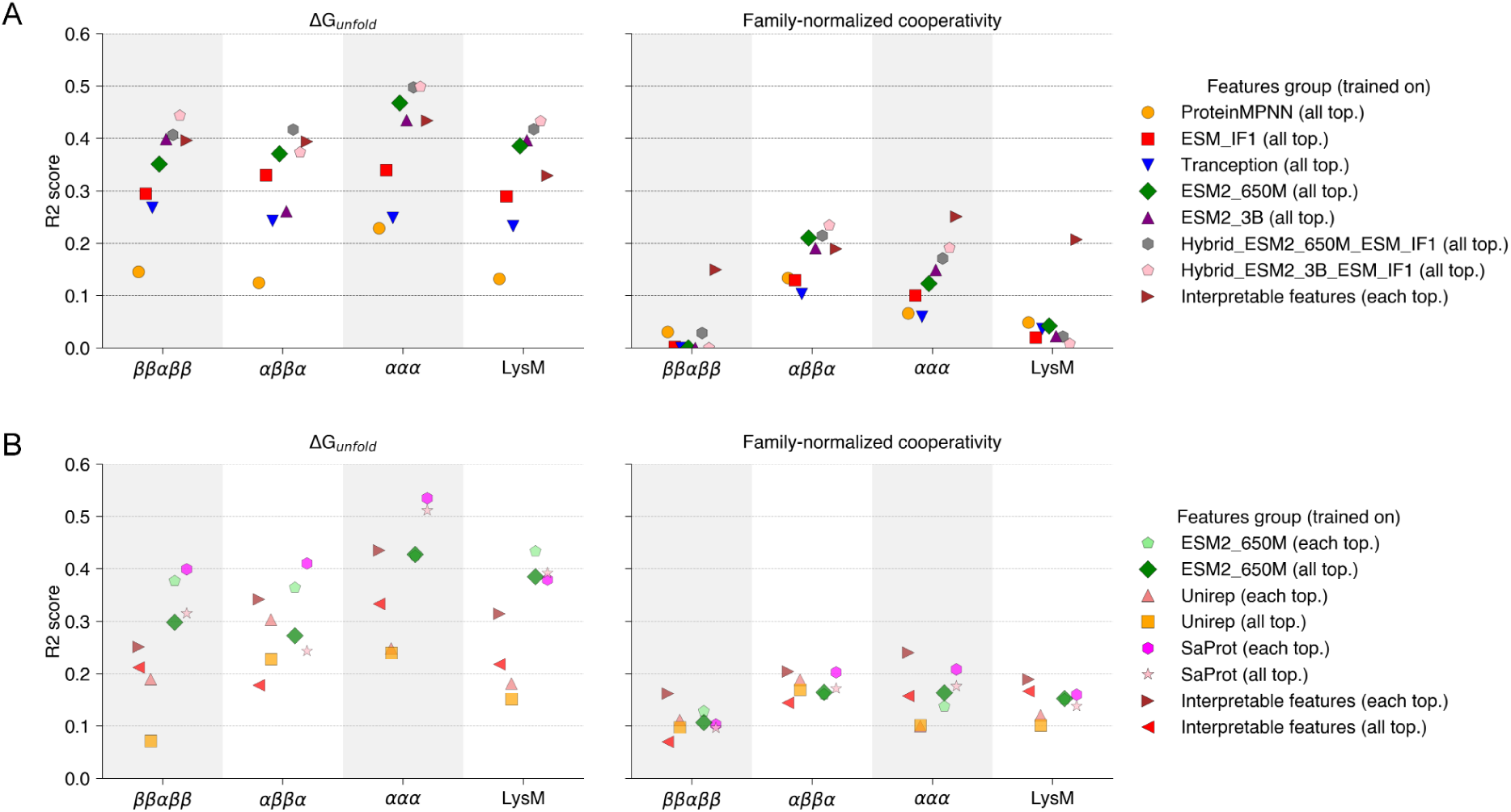
Performance of machine learning models. (A) First-generation models: R² scores for predictions of Δ*G*_unfold_ (left) and family-normalized cooperativity (right) are shown for models trained using protein language model embeddings derived from ProteinMPNN, ESM_IF1, Tranception, and ESM2 (650M and 3B parameters), as well as hybrid combinations, and also using interpretable features. In this initial approach, sequences were randomly assigned to 5-folds without identity-based clustering. (B) Second-generation models: R² scores for the same targets are presented for models incorporating dynamic clustering via MMseqs2 to group similar sequences, ensuring that each fold is as distinct as possible while maintaining balanced representation across protein families. In both cases, models were trained using 5-fold cross-validation over three independent random splits, and performance was evaluated in the aggregated predictions within each family-specific predictions. Models were trained either using examples from all four topologies (labeled as ‘’all top.’) or specifically with examples from the targeted topology (labeled as ‘each top.’).

**Figure S16.**
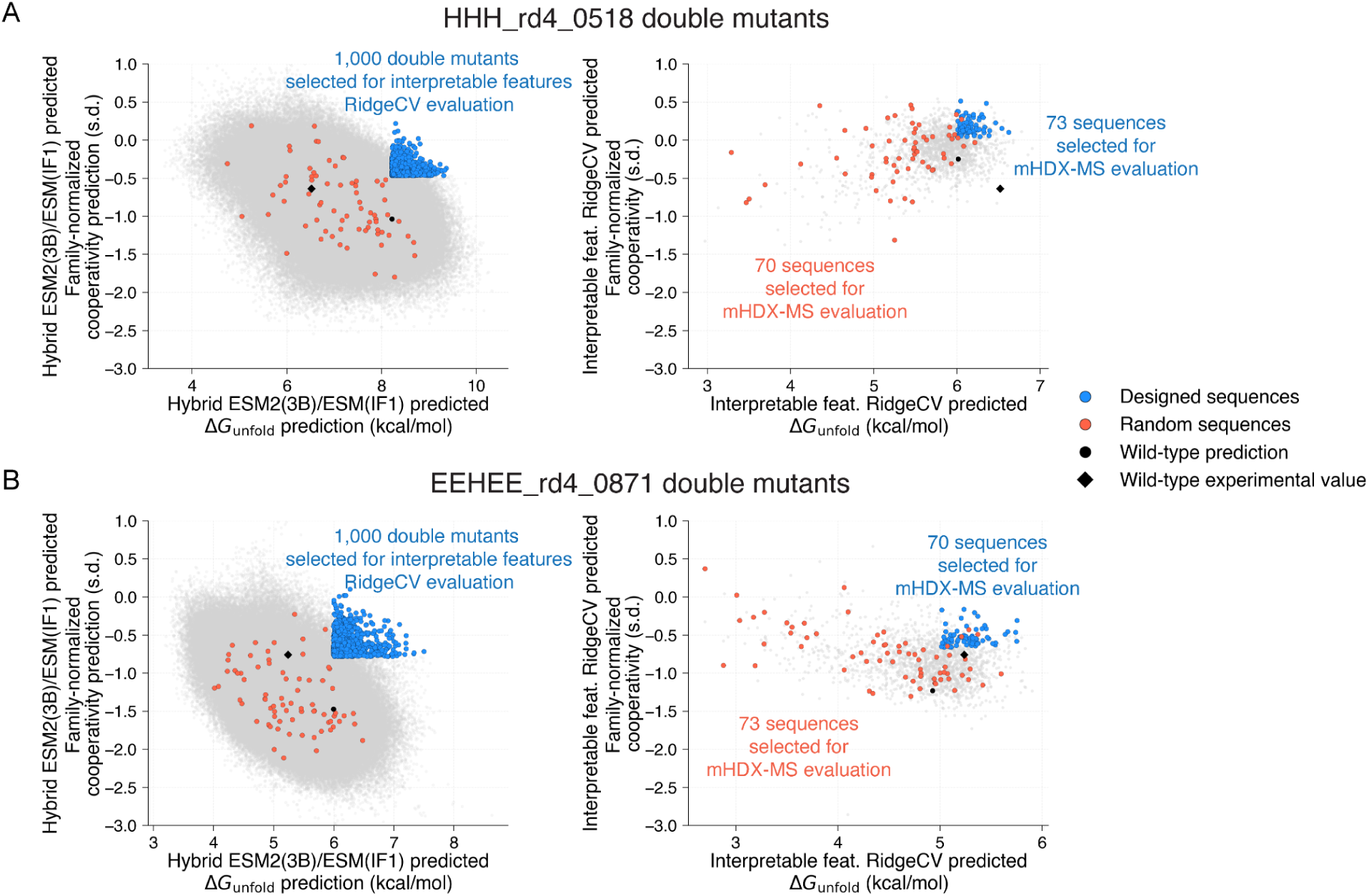
Selection of double mutants for experimental testing. (A) For HHH_rd4_0518, the best-performing first-generation protein language model was used to evaluate all possible double mutants (grey dots, left panel). The top 1,000 candidates were then selected (blue dots, right panel) and further evaluated using RidgeCV trained on interpretable features. Among these, 73 double mutants exhibiting both higher family-normalized cooperativity and Δ*G*_unfold_ compared to the wildtype (indicated by the black circle) were chosen for experimental testing; an additional 70 mutants were randomly selected (red dots, shown in both panels). (B) A similar workflow was applied to EEHEE_rd4_0871: all possible double mutants were initially evaluated (grey dots, left panel), and the top 1,000 candidates were filtered using the RidgeCV model trained on interpretable features (blue dots, left panel). From these, 73 double mutants (blue dots, right panel) with predicted properties exceeding those of the wildtype (black circle) were selected for testing, with another 70 mutants chosen at random (red dots, shown in both panels).

**Figure S17.**
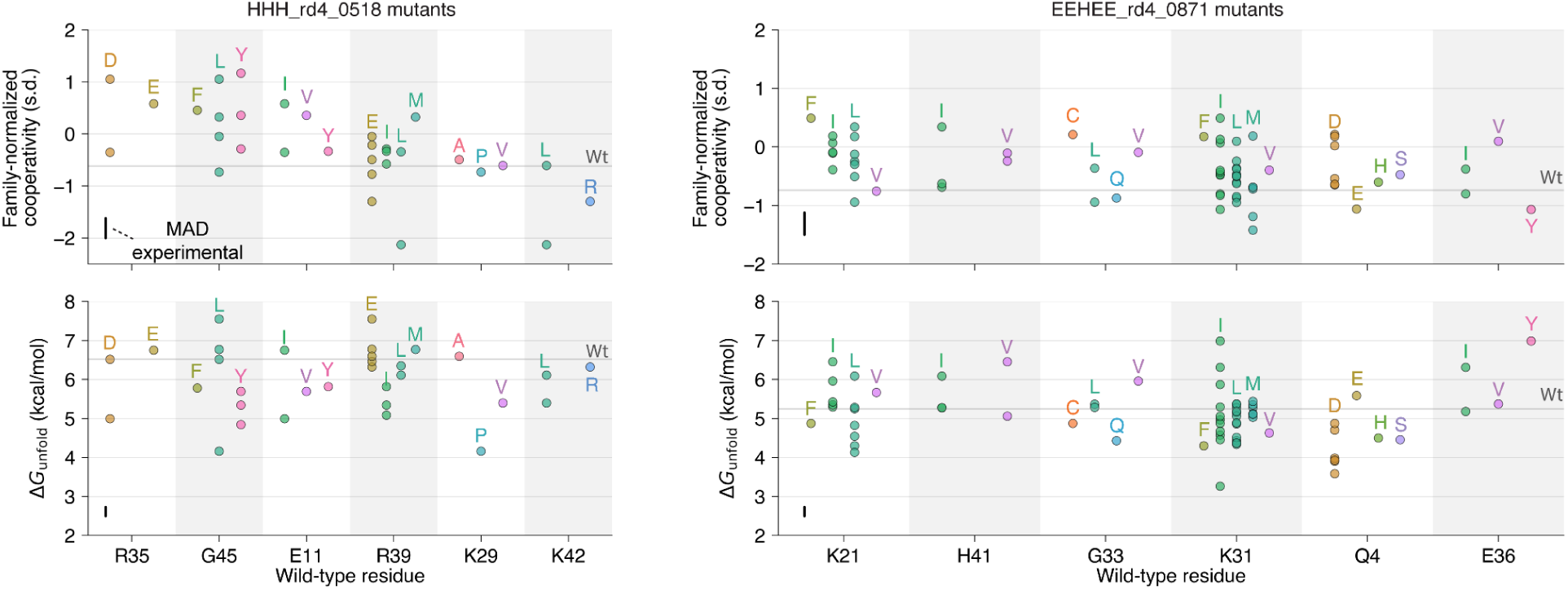
Impact of designed double mutants on protein Δ*G*_unfold_ and family-normalized cooperativity. (A) For HHH_rd4_0518, the top panel shows the changes in family-normalized cooperativity and the bottom panel shows the corresponding changes in Δ*G*_unfold_ for designed double mutants at the six most frequently mutated positions. Wild-type residues at these positions are ranked from highest to lowest average family-normalized cooperativity, providing a reference for assessing the impact of the mutations. Vertical scale bars show experimental mean absolute deviations (MAD) between replicates (of other proteins) measured in multiple libraries (**Fig. S5**). (B) As in A, for EEHEE_rd4_0871.

**Figure S18.**
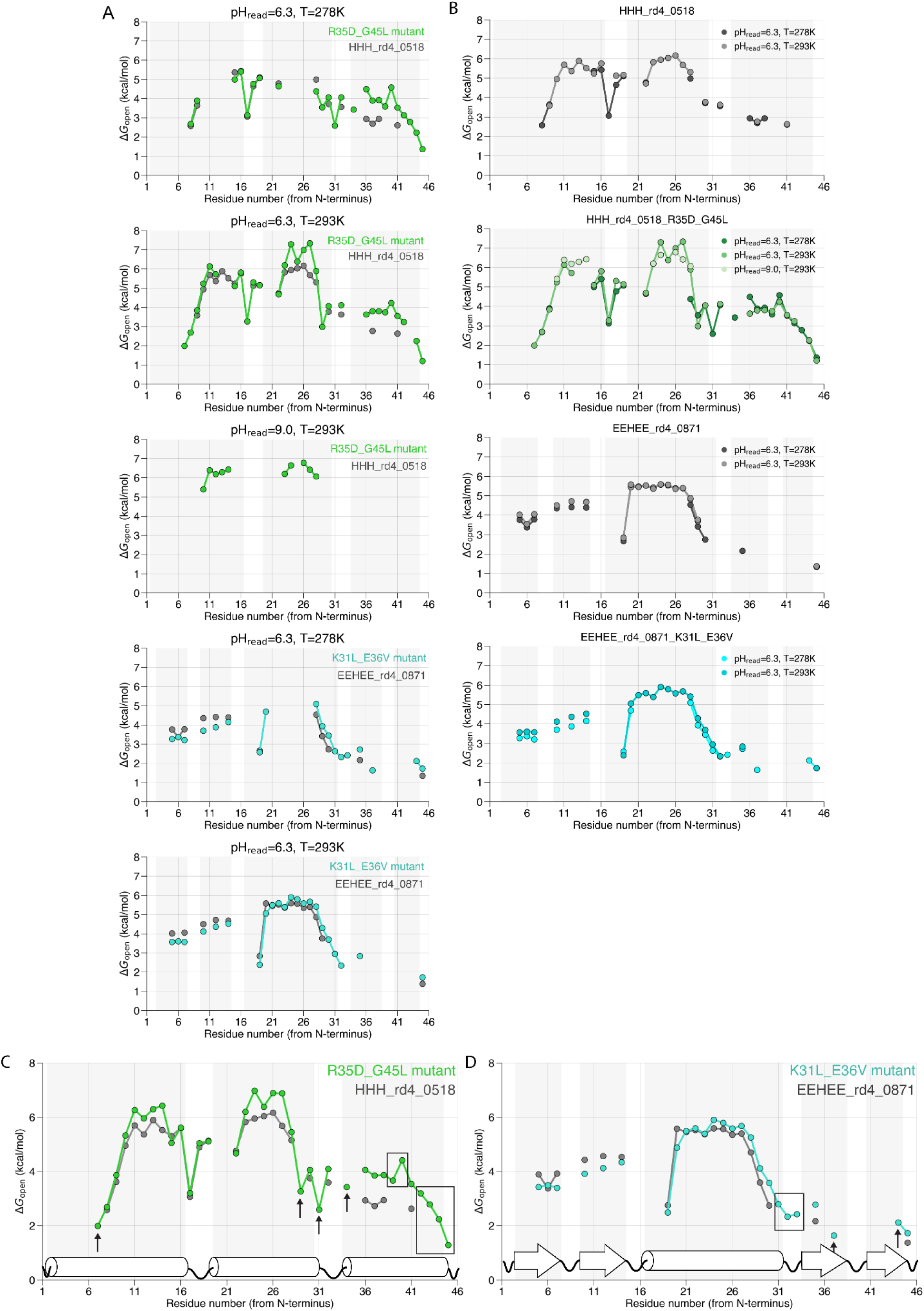
Residue-level HDX NMR analysis of wildtype and selected mutant proteins. (A) HDX NMR collected under multiple experimental conditions for HHH_rd4_0518 (wildtype) and its mutant, labeled as R34D_G45L for simplicity, and for EEHEE_rd4_0871 (wildtype) alongside its mutant K31L_E36V. (B) Four sub-panels display the HDX NMR data for each protein, with different shades representing the different experimental conditions. (C) Average residue opening free energies (Δ*G*_open_) for HHH_rd4_0518 and the R34D_G45L mutant are shown; for residues with protection measured under multiple conditions, the values are averaged. Boxes indicate groups of residues while arrows highlight individual residues where protection was observed in the mutant but not in the wildtype. (D) A similar comparison is presented for EEHEE_rd4_0871 and the K31L_E36V mutant.

**Figure S19.**
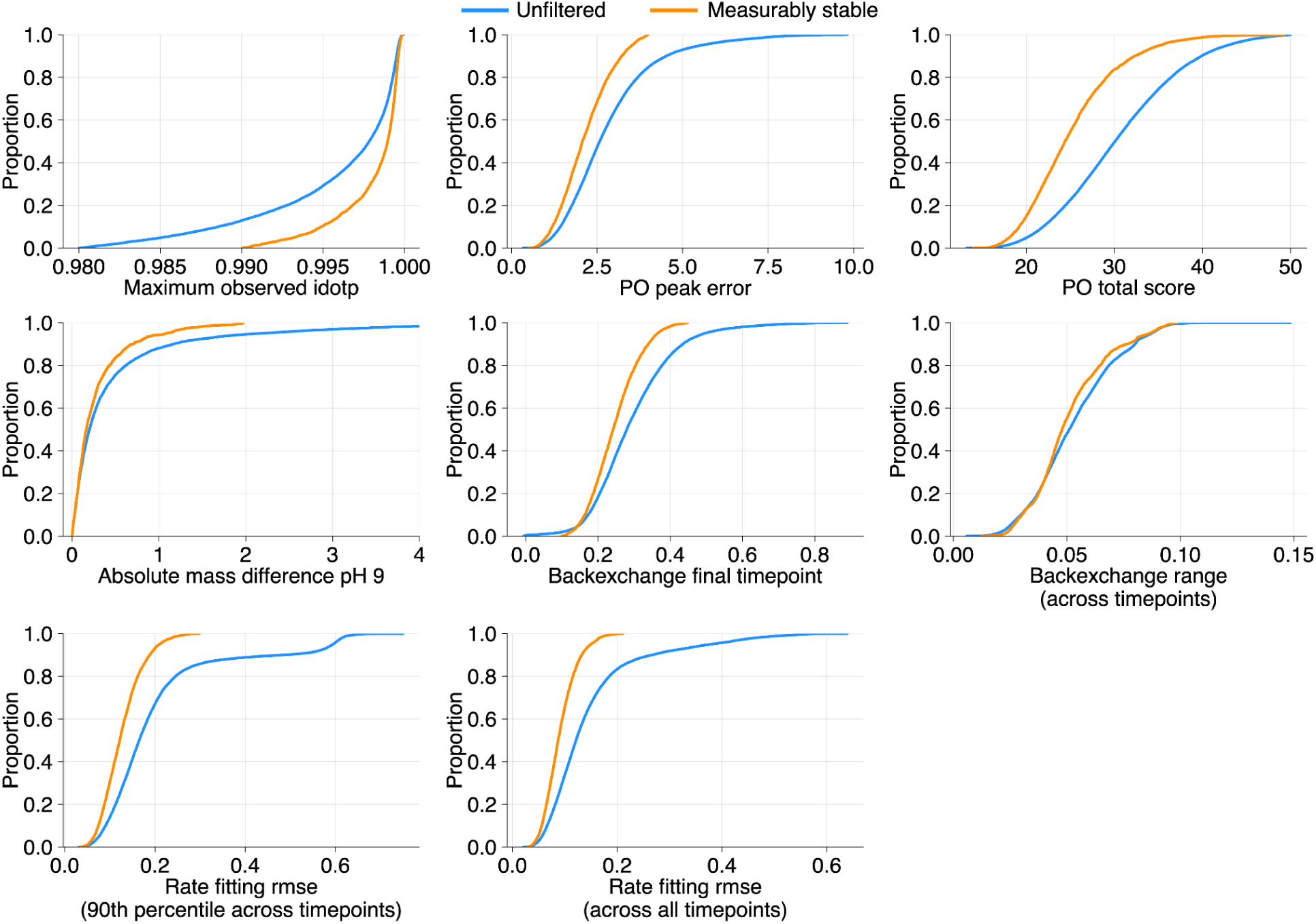
Cumulative distributions of quality control metrics. Blue curves show the cumulative distributions for 28,107 successful HDX measurements (note that individual proteins can be represented multiple times due to multiple retention times) that pass only the initial filtering criteria based on mass accuracy (<10 ppm), minimum isotopic distribution matching (idotp > 0.98), and a path optimizer (PO) total score (<50). Orange curves represent the cumulative distributions for the unique proteins selected to form the final dataset of 3,590 measurably stable proteins after applying additional, stricter quality control filters. These comparisons illustrate the impact of the combined filtering criteria on overall dataset quality. Notes: 1) the “absolute mass difference” is computed as the difference between the centroid mass at the last timepoint and that at the fifth-to-last timepoint after timepoint-specific back exchange correction; 2) the “back exchange range” is defined as the difference between the maximum and minimum back exchange values observed across all timepoints after applying timepoint-specific back exchange correction and, in the case of combined pH 6 and pH 9 data, experiment-wide back exchange normalization; 3) the rate fitting RMSE is computed as the root-mean-squared error between the predicted isotopic distribution and the experimentally observed isotopic distribution; we report either the 90th percentile of RMSE values across timepoints (second-to-last panel) or the average RMSE across timepoints (last panel).

**Figure S20.**
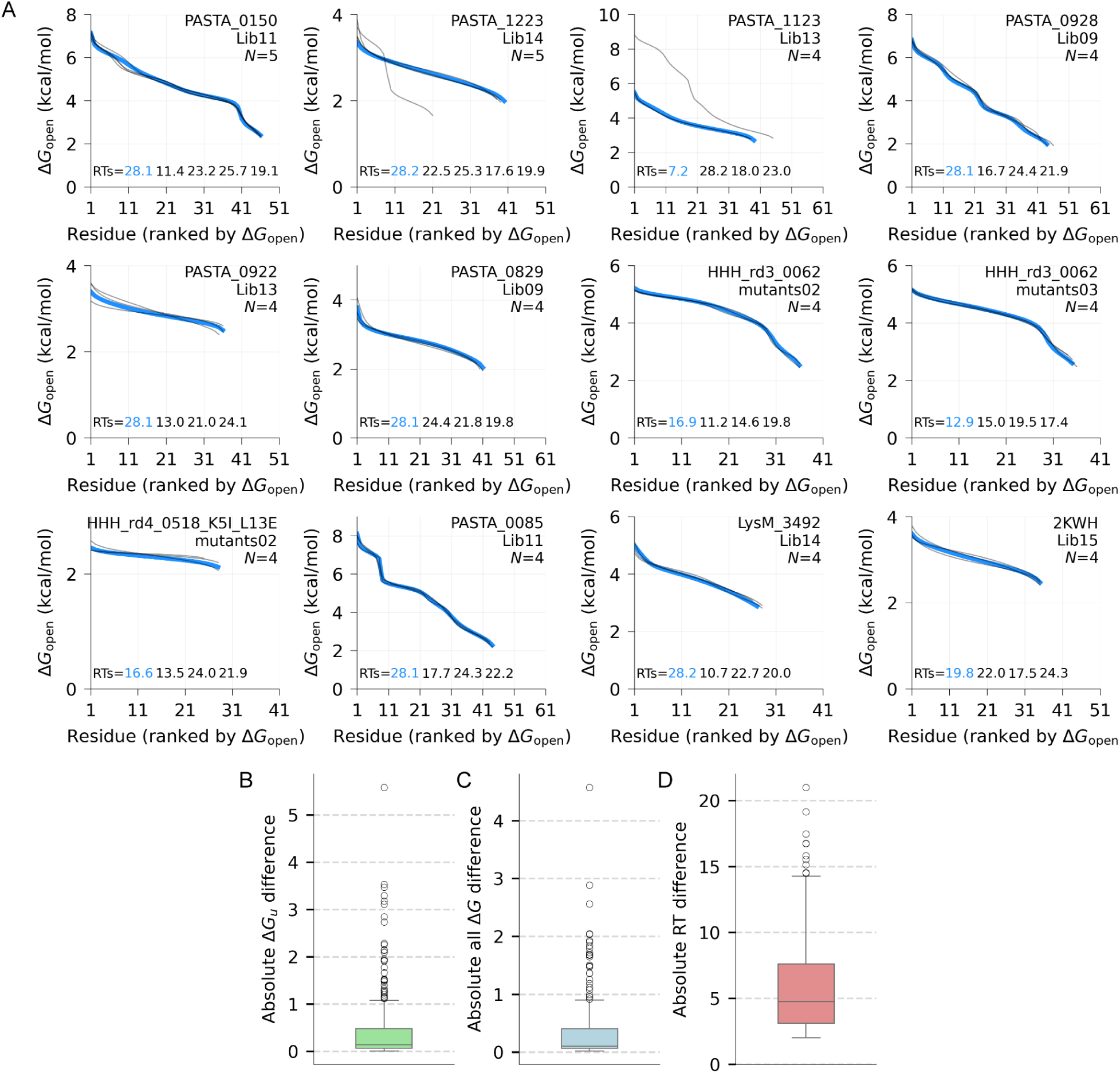
Reproducibility of mHDX-MS measurements across multiple retention times within the same library. (A) Examples of proteins exhibiting multiple retention times within a single experiment, yielding multiple successful HDX replicates. In total, 180 proteins were observed with at least two distinct retention times separated by more than 2 minutes. (B) Distribution of absolute differences in global stability (Δ*G*_unfold_) between replicates. (C) Distribution of all measured Δ*G* values across replicates. (D) Distribution of retention time differences for proteins with replicates that differ by at least 2 minutes.

**Table S1:** Datasets available at https://forms.gle/RwJwvfw6WN4gjXaD9

| Dataset name | Description | # of unique sequences |
| --- | --- | --- |
| Dataset_0_InitialOrder<br>(Fig. S9) | Initial DNA from all libraries | 15,715 |
| Dataset_1_UnfilteredData<br>(Fig. S19) | HDX results minimally filtered based on confident IDs and PO score < 50 | 8,293 |
| Dataset_2_SuccessfulHDX<br>(Fig. 1G, Fig. S5, Fig. S6, Fig. S11, Fig. S19, Fig. S20) | Successful HDX: proteins passing quality metrics described in the methods (includes EX1 kinetics and not fully deuterated proteins) | 5,778 |
| Dataset_3_MeasurablyStable<br>(Fig. 1C, Fig. 2A-F, Fig. 5B-D, Fig. S7, Fig. S12, Fig. S17) | Measably Stable: proteins that reached full deuteration, $\Delta G_{\text{unfold}}$ was greater than 2 kcal/mol, EX1 kinetics removed | 3,590 |
| Dataset_4_HDXNMR<br>(Fig. 1H, Fig. 3C and Fig. 5F, Fig. S3, Fig. S7, Fig. S18) | HDX NMR results: comprehensive dataset per condition, and average $\Delta G_{\text{open}}$ per position are provided as separate rows | 16<br>(13 proteins including some with/without His-tags) |
| Dataset_5_MesophilicThermophilic<br>(Fig. 2G) | Subset of proteins from natural domains from Dataset_3 which we classified as mesophilic or thermophilic (opt. growth temperature > 40°C) | 1,637 |
| Dataset_6_splits_interpretable<br>(Fig. 4, Fig. S13, Fig. S14) | Splits used for machine learning along with set of interpretable features | 3,193 |
| Dataset_6_splits_esm2 | Splits used for machine learning along with set of ESM2-derived features | 3,465 |
| Dataset_6_splits_unirep | Splits used for machine learning along with set of Unirep features | 3,465 |
| Dataset_6_splits_saprot | Splits used for machine learning along with set of SaProt features | 3,465 |
| Dataset_7_mHDX_cDNA<br>(Fig. S8) | Subset of proteins from Dataset_2 (best PO scored representative, excluded EX1 kinetics) that overlaps with available data from cDNA proteolysis assay (Tsuboyama et al. 2023) | 4,464 |
| Dataset_8_PDFs | Comprehensive set of plots generated by mhdX_pipeline and hdxrate_pipeline to evaluate time dependent mass distributions and their fits to rates. A jupyter-notebook is provided to help navigate through the pdfs. | - |
| Dataset_9_AlphaFoldModels | Set of modeled structures from Dataset_2_SuccessfulHDX | 5,778 |

**Table S2:** Parameters of cooperativity model fitting (Table S1: Dataset_3): Δ*G*_avg, expected_ = *a* (Δ*G*_unf_ - *b*)*^c^* · (fxn_hb)*^d^* + *e* (netq)

| PF | N | a | b | c | d | e | Pearson correlation | RMSE |
| --- | --- | --- | --- | --- | --- | --- | --- | --- |
| ααα | 604 | 1.7614 | 1.6552 | 0.5245 | 0.2341 | -0.0172 | 0.9368 | 0.2628 |
| WW | 253 | 0.7137 | 0.7551 | 1.0544 | 0.1236 | -0.0285 | 0.9160 | 0.2280 |
| LysM | 1,004 | 1.5436 | 1.6800 | 0.6364 | 0.2597 | -0.0241 | 0.9504 | 0.2584 |
| ββαββ | 329 | 1.6249 | 1.5824 | 0.6792 | 0.4643 | 0.0303 | 0.9358 | 0.3152 |
| βαββ | 170 | 1.5762 | 1.5393 | 0.5918 | 0.0607 | -0.0184 | 0.9474 | 0.2267 |
| PASTA | 305 | 1.9200 | 1.6959 | 0.5624 | 0.4146 | -0.0192 | 0.9561 | 0.3152 |
| αββα | 640 | 0.4650 | -0.0533 | 1.1308 | 0.0808 | -0.0093 | 0.8810 | 0.2744 |
| Cold-Shock | 60 | 1.2081 | 1.5092 | 0.6229 | 0.1626 | -0.0215 | 0.8478 | 0.3567 |
| PDB | 85 | 1.9810 | 1.8122 | 0.5657 | 0.6322 | -0.0181 | 0.9531 | 0.3381 |
| all | 3,465 | 1.4729 | 1.5818 | 0.5657 | 0.2569 | -0.0082 | 0.9401 | 0.3142 |

**Table S3:**
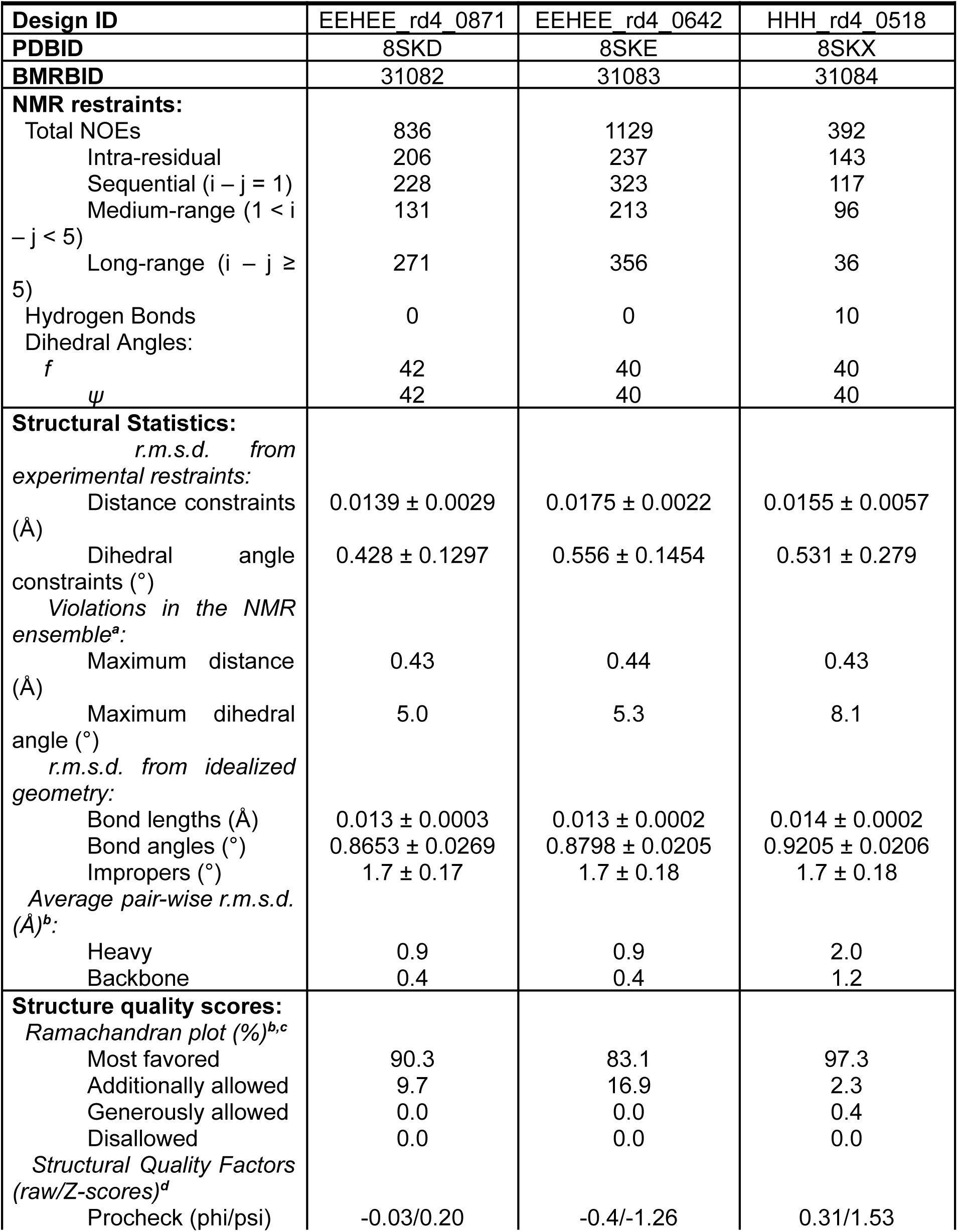
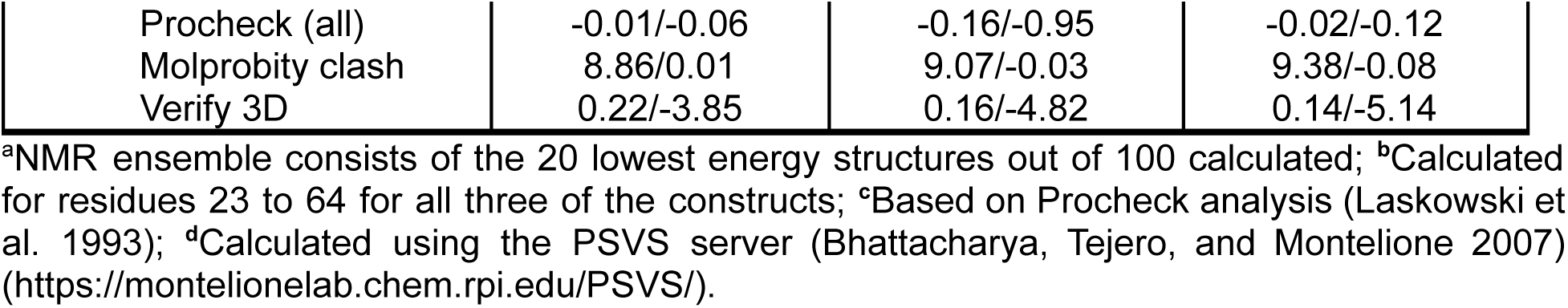
NMR restraints, structural statistics and quality scores for the miniproteins EEHEE_rd4_0871, EEHEE_rd4_0642 and HHH_rd4_0518.

